# Dynamic phosphoproteomics and proteomics uncover *Leishmania donovani*-driven ferritin hijacking, contributing to the control of iron homeostasis and iron-related oxidative stress

**DOI:** 10.64898/2026.02.25.707907

**Authors:** Sharvani Shrinivas Shintre, Florent Dingli, Nicolai Meyerhöfer, Olivier Gorgette, Catherine Thouvenot, David B. Blumenthal, Damarys Loew, Anne Silvestre, Najma Rachidi

**Affiliations:** Institut Pasteur, Université Paris Cité and INSERM U1347, Groupe de signalisation et interactions hôte-parasite, Paris, France; Institut Pasteur, Université de Paris and INSERM U1201, Unité de Parasitologie moléculaire et Signalisation, Paris, France; Laboratoire de Spectrométrie de Masse Protéomique (LSMP), Centre de Recherche, Institut Curie, PSL Research University, Paris, France; Biomedical Network Science Lab, Department Artificial Intelligence in Biomedical Engineering, Friedrich-Alexander-Universität Erlangen-Nürnberg, Erlangen, Germany; Institut Pasteur, Université Paris Cité, Ultrastructural Bioimaging Unit, 75015 Paris, France; INRAe, université de Tours, ISP, Nouzilly, France

**Keywords:** Phosphoproteome, Proteome, Signal transduction, Ferritin light chain 1, Iron homeostasis, *Leishmania*

## Abstract

*Leishmania donovani*, the causative agent of visceral leishmaniasis, survives within the parasitophorous vacuole (PV) of mammalian macrophages by extensively rewiring host cellular pathways. Although transcriptional and proteomic changes in infected macrophages have been characterized, the impact on the host phosphoproteome, a pivotal, reversible regulator of signalling, remains largely unexplored. To address this gap, we combined time-resolved quantitative phosphoproteomics and proteomics to map the dynamic response of murine macrophages to *L. donovani* infection. Early after infection, the parasite rapidly attenuates the macrophage signaling cascades normally triggered by phagocytosis. Between 24 h and 48 h post-infection we observed a progressive de-phosphorylation of proteins involved in innate immunity, apoptosis and other stress-responsive pathways, consistent with a partial shutdown of multiple host kinases. From 24 h onward, a global decline in protein abundance was also detected, most notably within lysosomal network. Strikingly, only five host proteins were consistently up-regulated, suggesting their importance for parasite survival. Ferritin light chain (Ftl1) displayed the largest increase. Immunofluorescence and transmission-electron microscopy revealed that Ftl1 accumulates within the parasitophorous vacuole, colocalizes with the ferritin receptor Ncoa4, and yet fails to undergo degradation, indicating that *Leishmania* co-opts ferritin as its own intracellular iron-storage compartment. Ferritin is not confined to the PV; it is also internalized by the parasite. Within *L. donovani*, ferritin accumulates not only in the cytoplasm but also in the nucleus, where it may function both as an iron-storage depot and as an iron buffer that protects the parasite from oxidative damage. Finally, we demonstrate that *Leishmania donovani* pre-conditions its host macrophage for the iron-rich environments of the liver and spleen by driving the assembly of ferritin particles enriched in ferritin-light chain. This contrasts with *L. amazonensis*, which, residing in the iron-poor skin, induces ferritin predominated by ferritin-heavy chain (Fth1). Knock-down of *Ftl1* did not decrease parasite survival because macrophages compensated for its loss by up-regulating *Hspb1*, which encodes a protein that limits lipid peroxidation by inhibiting the Fenton reaction. Although this response protects the host cell, it also diminishes iron import, imposing a metabolic cost on *Leishmania*. *Hspb1* represents only one example; additional, as yet unidentified, compensatory pathways are likely activated by the parasite to mitigate the loss of ferritin-light chain. Collectively, our data uncover a previously unknown strategy whereby *L. donovani* hijacks host ferritin trafficking to create a protected iron reservoir, thereby preventing ferroptosis. This mechanism sets *Leishmania*, a eukaryotic intracellular parasite, apart from the canonical iron-acquisition tactics employed by bacteria and fungi.

## Introduction

*Leishmania* parasites are part of a group of pathogens, including *mycobacteria*, that establish their niche within immune cells, particularly macrophages, and effectively disarm the immune system. Gaining insight into how these pathogens survive in such hostile environments is essential for developing next-generation therapies that will overcome pathogen resistance. In this context, *Leishmania*, serves as an excellent model for studying host-pathogen interactions under these extreme conditions, as a representative of eucaryotic pathogens. *Leishmania donovani* is the causative agent of visceral leishmaniasis, also known as kala-azar, which infects 30,000 people annually. If untreated, visceral leishmaniasis is fatal in over 95% of the cases (WHO, 2023). The parasite completes its lifecycle in two hosts, the female phlebotomine sandfly, which acts as the vector, and the mammalian host. During a blood meal, an infected sandfly transmits infective metacyclic promastigotes to the mammalian host. Once phagocytosed by macrophages, these parasites reside within the parasitophorous vacuole, where acidification triggers their differentiation into amastigotes ^1^.

To create a niche within the macrophage and proliferate, the parasite alters various host cell pathways including metabolism^2,3,4^ translation and immune response. For instance, *Leishmania* attenuates host immune responses by activating the host protein tyrosine phosphatase Shp-1 through its direct interaction with *Leishmania* elongation factor-1 alpha (EF1α), leading to the inhibition of macrophage antimicrobial functions (NO, TNFα or ROS) ^5, 6^. *Leishmania donovani* remodels host cell pathways on multiple regulatory levels, host apoptosis is a striking example ^7^. At the transcriptomic level, the parasite inhibits apoptosis by downregulating pro-apoptotic gene expression while upregulating anti-apoptotic genes, such as Bcl2 ^8, 9^. At the proteomic level, *L. donovani* further inhibits apoptosis by reducing the abundance of annexin A5, a marker of early apoptosis, and increasing levels of vimentin, a known anti-apoptotic factor ^10, 11^. Beyond transcriptional and proteomic regulation, *L. donovani* also reprograms the host epigenome, altering cytosine methylation patterns to modulate the expression of key genes within the Mapk and Jak/Stat signalling pathways ^12^. These multi-tiered strategies highlight the parasite sophisticated manipulation of host cell death pathways to ensure its survival and persistence.

Numerous host pathways have been shown to be modulated by *Leishmania* through unbiased approaches, such as transcriptomics, and, to a lesser extent, proteomics^13^. These studies have provided valuable snapshots of the regulatory landscapes altered during infection. However, one critical layer of cellular regulation remains largely unexplored in the context of *Leishmania* infection: post-translational modifications (PTMs). PTMs are rapid, reversible, and dynamic changes that serve as key regulators of protein function and cellular behaviour. Among them, protein phosphorylation stands out as one of the most abundant and functionally significant in humans, orchestrating essential physiological processes such as protein activity, stability, protein–protein interactions, and gene transcription^14^. Notably, phosphorylation is a fundamental mechanism that regulates both innate and adaptive immune responses ^15,16^. A clear illustration of this is the suppression of host protein kinase C activity during *Leishmania* infection, thereby dampening downstream host immune responses ^17,18^. These examples clearly illustrate that modulation of host signalling, particularly through the regulation of kinases and phosphatases, might be a central strategy employed by *Leishmania* to rapidly and extensively rewire host cellular pathways, ultimately ensuring its survival within the host ^19, 20^. This is further supported by the observation that *Leishmania* parasites actively secrete parasite-derived kinases into host cells ^21–24^.

Despite the importance of host signalling regulation for parasite survival, an assessment of the host phosphoproteome alterations induced by *Leishmania donovani* infection, and their dynamic evolution over time, has not yet been undertaken. To address this critical gap, our study investigates the modulation of the host phosphoproteome over the first 48 hours of *L. donovani* infection. This time-resolved, systems-level approach offers a more comprehensive understanding of host-pathogen interactions than previously available, enabling us to uncover early signalling events and cellular pathways that the parasite targets as it establishes its intracellular niche. This study demonstrates that *Leishmania donovani* infection induces far more extensive changes on the host phosphoproteome than on the host proteome, highlighting post-translational modifications as a key mechanism through which the parasite reprograms host cell function. These phosphorylation dynamics are closely linked to the activation and inactivation of host kinases. In contrast, the host proteome undergoes only modest changes. Only a very small subsets of proteins are still up-regulated at 48h post infection, likely reflecting their importance for parasite survival; among them are ferritin and its autophagy adaptor Ncoa4, both of which regulate host iron homeostasis. We show that *Leishmania* specifically recruits host ferritin to the parasitophorous vacuoles via Ncoa4-mediated interactions, while preventing ferritinophagy. Knockdown of Ftl1 alone does not markedly impair *Leishmania donovani* infection, because the parasite directly or indirectly activates compensatory mechanisms, such as the up-regulation of Hspb1, which can mitigate lipid peroxidation by inhibiting the Fenton reaction. Our study identifies a novel mechanism by which *L. donovani* hijacks ferritin trafficking and degradation to sequester host iron, creating a localized reservoir while avoiding host lipid peroxidation. This mechanism underscores the distinct strategies by which prokaryotes and eukaryotes regulate iron homeostasis.

## Results

### *Leishmania donovani* attenuates the host responses to phagocytosis

To elucidate the impact of *Leishmania donovani* infection on the host phosphoproteome and investigate host signalling dynamic evolution over the course of infection, we performed a temporal phosphoproteomic profiling of infected macrophages. Murine bone marrow-derived macrophages (BMDMs) were generated, and either maintained uninfected or infected with *L. donovani* amastigotes at a multiplicity of infection (MOI) of 10 for 4, 24, or 48 hours post-infection (hpi). Given that infection by *L. donovani* involves phagocytic uptake of the parasite ^25^, a phagocytosis control was included wherein BMDMs were exposed to zymosan particles for 4 hours. The experimental workflow is illustrated in Supplementary Figure S1A. At each timepoint, cells were lysed, and protein extracts were subjected to a phosphoproteomic profiling by mass spectrometry, using five biological replicates per condition. Datasets were retrieved from MyProms ^26^ and filtered based on biological reproducibility. Four comparisons were made to assess phosphorylation changes in four conditions (1) 4h Zymosan versus 4h uninfected (Zymo 4h/UI 4h, 8849 phosphosites), (2) 4hpi versus 4h uninfected (Inf 4h/UI 4h, 8294 phosphosites), (3) 24hpi versus 24h uninfected (Inf 24h/UI 24h, 8863 phosphosites), and (4) 48hpi versus 48h uninfected (Inf 48h/UI 48h, 9751 phosphosites). The corresponding volcano plots are shown Figure S1B. All phosphosites, quantified in at least three biological replicates, are reported Table S1, while a filtered dataset quantified in at least three biological replicates and including only phosphosites with statistical significance (p < 0.05) is provided in Table S2 for all conditions.

We first investigated the impact of *Leishmania* phagocytosis on the host phosphoproteome, comparing Zymo-4h/UI-4h to Inf-4h/UI-4h. Upon zymosan phagocytosis, 2,276 phosphosites were differentially regulated, of which 1,533 with a fold change above 2 or below 0.5. In contrast, only 200 phosphosites were differentially regulated during *Leishmania* phagocytosis, of which 143 with a fold change above 2 or below 0.5 (Figure 1A). This is a 10 fold reduction compared to the Zymo-4h/UI-4h condition. These findings indicate that macrophage signalling responses to *L. donovani* phagocytosis strongly differ from those triggered by classical phagocytosis, exemplified by zymosan. These results also suggest that upon *L. donovani* infection, the differentially regulated phosphosites might have a significant importance for parasite survival.

**FIGURE 1:**
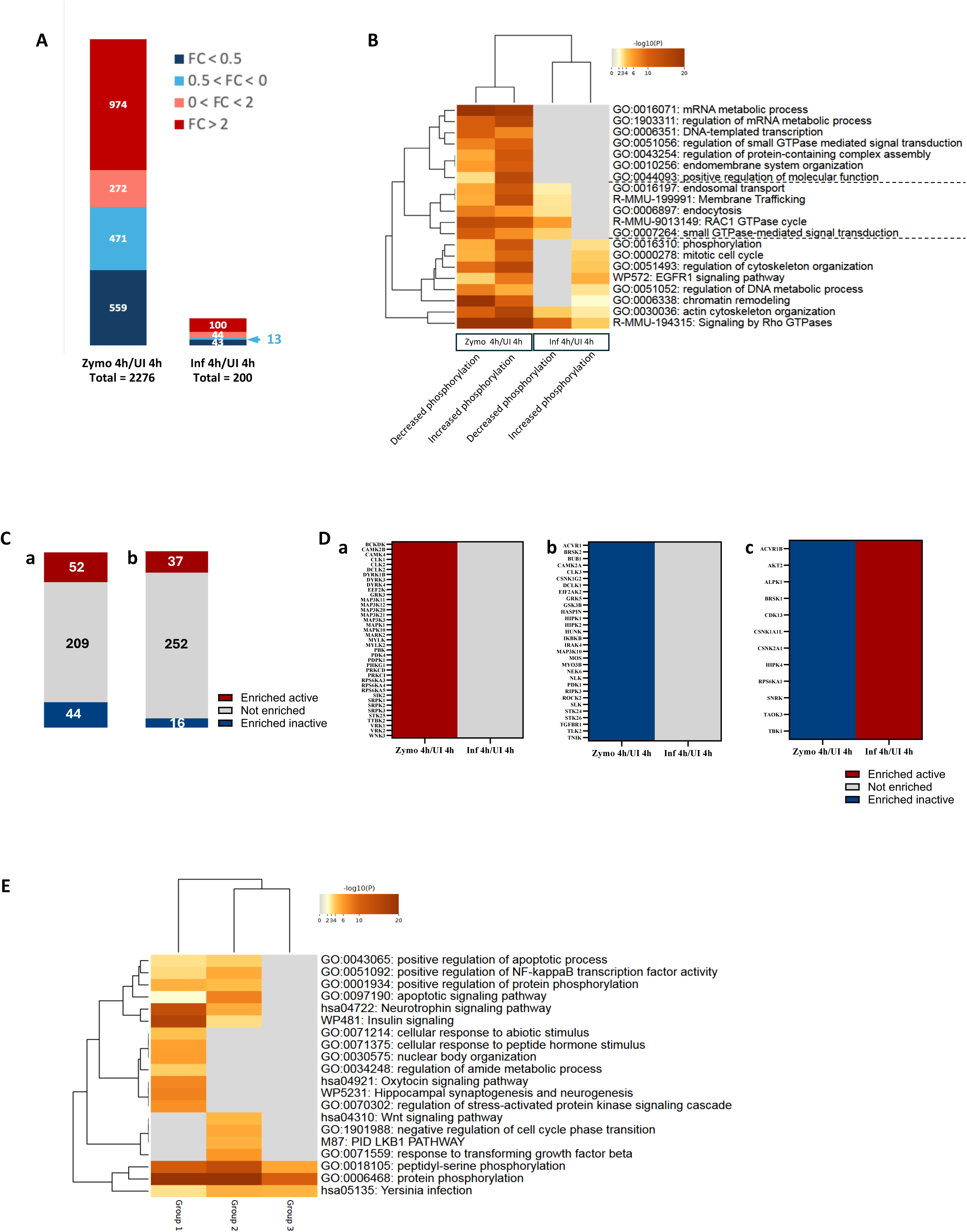
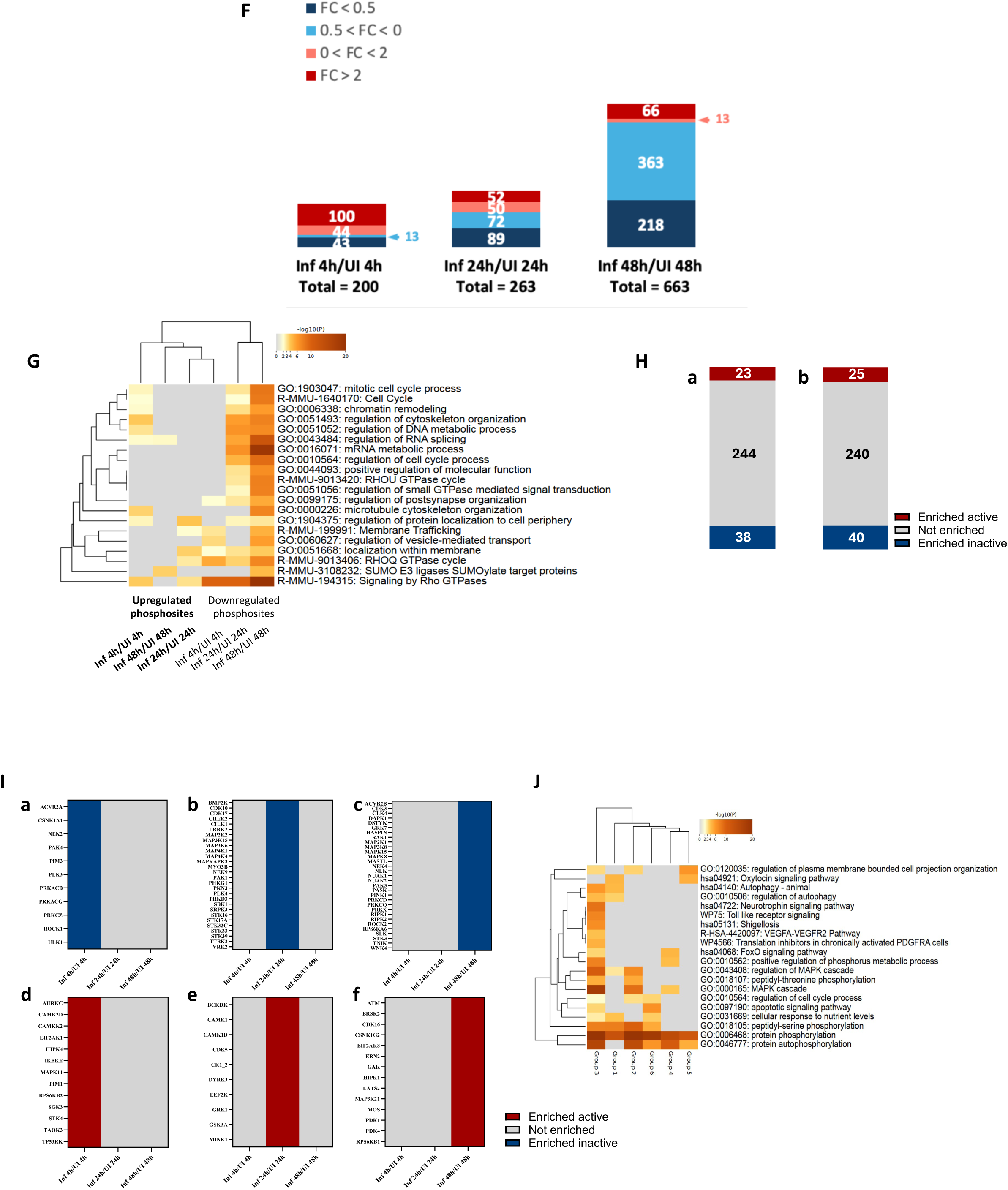
Modulation of host phosphoprotein by *L. donovani*. **A.** Distribution of fold changes in the phosphorylation levels of differentially regulated phosphosites (p<0.05, any FC, quantified in at least three replicates) is given for 4h Zymosan and 4hpi. The total number of phosphosites is presented below each plot. The color code for the four brackets for fold change values is given in the legend. The number of phosphosites present in each of the fold change brackets is given inside the plot. The phosphoproteome has been done in five replicates. **B.** Functional impact of phosphorylation modulation by *L. donovani*. GO term enrichment of 4h Zymosan and 4hpi for phosphosites with increased (Log2FC>0) and decreased (Log2FC<0) phosphorylation. The intensity of orange provides the level of significance of the enrichment. Grey indicates that the pathway was not enriched. **C.** Histograms describing the number of enriched active and inactive kinases during a) zymosan treatment and b) 4h infection. **D.** Kinases with their activities modulated during infection were divided into three groups based on their regulation: a) Active in 4h zymosan, not enriched in 4hpi, b) inactive in 4h zymosan, not enriched in 4hpi, c) inactive in 4h zymosan but active in 4hpi. **E.** Functional enrichment of the three groups of kinases using Metascape. The intensity of orange provides the level of significance of the enrichment. Grey indicates that the pathway was not enriched. The intensity of orange provides the level of significance of the enrichment. Grey indicates that the pathway was not enriched. **F.** Distribution of fold changes in the phosphorylation levels of differentially regulated phosphosites (p<0.05, any FC, quantified in at least three replicates) is given for 4h, 24h and 48h post infection. The 4hpi is the same as that presented in Figure 2 but has been repeated here for the reader comfort. The total number of phosphosites is presented below each plot. The color code for the four brackets for fold change values is given in the legend. The number of phosphosites present in each of the fold change brackets is given inside the plot. The phosphoproteome has been done in five replicates. **G.** Functional impact of phosphorylation modulation by *L. donovani*. GO term enrichment of 4h Zymosan and 4hpi for phosphosites with increased (Log2FC>0) and decreased (Log2FC<0) phosphorylation. The intensity of orange provides the level of significance of the enrichment. Grey indicates that the pathway was not enriched. **H.** Histograms describing the number of enriched active and inactive kinases in a) 24hpi and b) 48hpi conditions. **I.** Kinases showing time-dependent activities divided into 6 groups depending on the timepoint and activity status: a) Inactive at 4hpi, b) Inactive at 24hpi, c) Inactive at 48hpi, d) Active at 4hpi, e) Active at 24hpi, f) Active at 48hpi. **J.** Functional GO term enrichment of the six groups using Metascape. The intensity of orange provides the level of significance of the enrichment. Grey indicates that the pathway was not enriched.

We next performed a pathway enrichment, using the "Multiple Gene List" feature in Metascape ^27^ with default parameters. The resulting enrichment map comparing Zymo-4h/UI-4h to Inf-4h/UI-4h is presented Figure 1B. In contrast to Zymo-4h/UI-4h, the pathways enriched in the Inf-4h/UI-4h up-regulated phosphosite dataset, such as ‘chromatin remodelling’, ‘EGFR1 signalling pathway’ and ‘regulation of DNA metabolism process’, are different from those enriched in the Inf-4h/UI-4h down-regulated phosphosite dataset, including ‘small GTPase-mediated signalling’, ‘endosomal transport’, ‘membrane trafficking’ and ‘endocytosis’. The only common regulated pathways were ‘actin cytoskeleton organisation’ and ‘Rho GTPases signalling’. Additionally, several pathways were not enriched in the Inf-4h/UI-4h dataset compared to Zymo-4h/UI-4h, such as ‘mRNA metabolic process’, ‘DNA-templated transcription’ or ‘endomembrane system organisation’. This finding suggests that infection with *L. donovani* might selectively suppress the regulation of processes linked to transcription and mRNA processing, which might explain the low level of transcriptional regulation detected during *Leishmania* infection (mostly below 2 fold change) ^28^.

### Regulation of host kinases partly affect phagocytosis signalling

Interpreting phosphoproteomic data is notoriously challenging. presence or absence of phosphorylation can be associated with a wide spectrum of cellular outcomes, protein degradation, activation or inactivation, and conformational changes, yet the literature that directly connects specific phosphosites to these functional consequences is still sparse. To gain insight into host signaling dynamics during *Leishmania* infection, we conducted motif enrichment analysis of our phosphoproteomic datasets. These analyses employed an established experimental-based pipeline developed by Johnson *et al.* ^29^. Serine/threonine kinase activities from Zymo-4h/UI-4h and Inf-4h/UI-4h inferred from motif-based enrichment obtained experimentally were visualized using volcano plots, displaying motif frequency against adjusted p-values (Figure S2A). Kinases exhibiting an enrichment value Log₂FC < 0 and an adjusted p-value ≤ 0.05 (blue, Figure S2B) were interpreted as inactivated kinases (lower activity compared to the basal level represented by the uninfected controls, whereas those exhibiting an enrichment value Log₂FC > 1 and an adjusted p-value ≤ 0.05 (red, Figure S2B) were considered as activated kinases (higher activity compared to the basal uninfected activity). The number of both activated and inactivated kinases declined during the first 4 hours post-infection (Figure 1C-b) compared to the 4 hours post Zymosan, returning to the basal activity levels observed in the uninfected control (Figure 1C-a). These results suggest that *Leishmania* blocks the up- or down-regulation of the kinases normally differentially regulated during phagocytosis.

To identify the functional characteristics of these kinases, they were grouped into three categories: **Group 1**: Kinases activated during zymosan phagocytosis but at basal level during *Leishmania* phagocytosis (39, Figure 1D-a). **Group 2**: Kinases inactivated during phagocytosis but not during *Leishmania* phagocytosis (30, Figure 1D-b). **Group 3**: Kinases inactivated during phagocytosis but activated during *Leishmania* phagocytosis (12, Figure 1D-c). Only two kinases, Pim3 and Prkacb, were activated after Zymosan phagocytosis and inactivated after *Leishmania* phagocytosis, so will not be considered for further analysis. We next conducted a functional enrichment analysis of each group, using Metascape (Figure 1E). Kinases in Groups 1 and 2, which are up- or down-regulated after Zymosan phagocytosis but not regulated after *Leishmania* phagocytosis, were associated with ‘positive regulation of apoptosis’ ^7^ and ‘NF-κB signalling’ ^30, 31^, ‘Wnt signalling pathways’ ^32, 33^ and ‘response to Tgfβ’ ^34, 35^. These pathways, which participate in immune and cellular stress responses, are detrimental to parasite survival. Thus, the apparent deregulation of these pathways by *L. donovani* likely represents a deliberate strategy to interfere with the host-defence programs that are normally triggered during phagocytosis. In contrast, Group 3, including kinases such as Ck1 and Akt2 are inactivated during zymosan phagocytosis, while activated during *Leishmania* phagocytosis. This group is highly heterogeneous and limited in size, precluding robust GO-term enrichment analysis. The only GO terms that emerge are ‘protein phosphorylation’ and ‘*Yersinia* infection’, which are shared across all three groups. The putative activation of Group 3 kinases in the context of *Leishmania* infection is corroborated by previous studies; for example, *Leishmania* has been shown to activate host Akt, which in turn enhances macrophage resistance to apoptosis^36^.

### Global Phosphoproteomic Profiling Reveals Temporal Modulation of Host Signalling by Leishmania donovani

To characterise the temporal dynamics of the phosphoproteome of *Leishmania* infected macrophages (LIMs), we compared phosphosite abundances at 4, 24, and 48 hpi (Figure 1F). In the course of the analysis, the number of significantly regulated phosphosites increased markedly from 200 at 4 hpi to 663 at 48 hpi (Figure 1F). This expansion was driven predominantly by an increase in downregulated phosphosites, as their percentage increased from 28% at 4 hpi, to 61% at 24 hpi and 87% at 48 hpi, reflecting a global and progressive dephosphorylation over the first 48h of infection. Pathway enrichment analysis of the down regulated phosphosites (log₂FC < 0, Figure 1G) revealed biological processes that were exclusively enriched in this dataset such as ‘mRNA metabolism’, ‘regulation of small-GTPase-mediated signal transduction’, and ‘regulation of vesicle-mediated transport’, with the number and significance of enriched terms increasing from 4 h to 48 hpi. The pathway enrichment corresponding to up-regulated phosphosites (log₂FC > 0) revealed distinct temporal signatures (Figure 1G). At 4 hpi, the most enriched GO terms were ‘chromatin remodelling’, ‘regulation of RNA splicing’, ‘DNA metabolic processes’, and ‘cytoskeleton organization’, indicating early parasite-driven re-programming of epigenetic, post-transcriptional mechanisms and cellular structure. By 24 hpi, the enrichment shifted toward ‘membrane trafficking’ and ‘membrane localization’, suggesting a re-wiring of vesicular transport. At 48 h pi only two categories remained significantly enriched: ‘regulation of RNA splicing’ and ‘SUMO-E3 ligase-mediated SUMOylation’, consistent with previous reports showing that *Leishmania* up-regulates host SUMOylation machinery to dampen immune signalling and promote intracellular survival ^37^. Together, these results suggest that *Leishmania* orchestrates a temporally ordered phosphorylation landscape: an early regulation of chromatin and RNA-processing pathways, a mid-phase regulation of membrane dynamics, and a late-phase regulation of RNA splicing and SUMOylation, while simultaneously driving a broad dephosphorylation that may attenuate multiple cellular functions.

### Host kinases partly shapes the dynamic phosphoproteome during *Leishmania* infection

To assess whether the observed dephosphorylation pattern during *Leishmania* infection aligned with changes in host kinase activity, we performed kinase enrichment analyses. These analyses compared kinase activity under infected conditions to uninfected conditions at 4 hpi (Figure S2A-b), 24 hpi (Figure S2A-c) and 48 hpi (Figure S2A-d). The results, presented in the bar plots (Figure 1C-b and 1H), revealed a decrease in the number of active kinases: from 37 at 4 hpi to 23 at 24 hpi, and 25 at 48 hpi. Concurrently, inactive kinases increased: from 16 at 4hpi to 38 at 24hpi and 40 at 48hpi. This progressive inactivation of host kinases during the course of infection correlates with the observed dephosphorylation of phosphosites. Additionally, these results suggest that the parasite modulates host signalling, at least in part, by restoring host kinase activity to the baseline levels observed in uninfected cells. To explore temporal dynamics in kinase activity, six groups of kinases were created based on their activity at specific infection time points, as shown in Figure 1I. **Group 1**: Inactive only at 4hpi (Figure 1I-a). **Group 2**: Inactive only at 24hpi (Figure 1I-b); **Group 3**: Inactive only at 48hpi (Figure 1I-c); **Group 4**: Active only at 4hpi (Figure 1I-d); **Group 5**: Active only at 24hpi (Figure 1I-e); **Group 6**: Active only at 48hpi (Figure 1I-f).

The functional enrichment analysis of these groups revealed distinct pathway associations linked with temporal modulations of kinase activities. Groups 1 and 4 (4 hpi) were linked to autophagy regulation, MAPK signalling, response to nutrient levels, and protein phosphorylation. Groups 2 and 5 (24 hpi) were associated with the MAPK cascade, peptidyl-threonine phosphorylation, and membrane-bound cell projections, while groups 3 and 6 (48 hpi) were enriched in a broad range of pathways, including autophagy, Toll-like receptor signalling, FoxO signalling, MAPK cascade, apoptosis, and phosphorylation. However, most of these regulations appeared to be associated with inactivated kinases at 48h (group 3), suggesting widespread suppression of host kinase activity late in infection, which is consistent with the strong number of down-regulated phosphosites at 48h. Thus, the dynamic regulation of the host phosphoproteome during *Leishmania* infection can be attributed, in part, to the modulations of host kinase activity.

Based on our analysis, *L. donovani* might suppress important immune and stress-related signalling pathways through the down-regulation of host kinases. For instances, Ikbke, which phosphorylates Stat3 and NF-kB to promote pro-inflammatory signalling, is only activated at 4 hpi and return to basal uninfected level from 24 hpi (Figure S2E, blue arrow). Similar finding for the MAPK signalling cascade (Figure S2E, green arrows), activator of th1-regulated processes such as NF-kB pathway important for innate immunity^38^, which is active at 4h but returned to basal level from 24h. This is in line with the literature, as MAPK signalling is found to be activated in the first few hours of infection and then subsequently inhibited ^39, 40^. Our findings also align with the known activation of host phosphatases, such as Shp-1 and Ptp-1B, by *Leishmania* parasites through effectors like *Leishmania* elongation factor 1α, that inhibit MAPK activation^5^. These examples demonstrate that kinase-activity inference from the phosphoproteome generally agrees with previously reported findings. Consequently, such inference represents a valuable first step toward the biological interpretation of phosphoproteomics datasets, especially given the lack of functional annotations for many phosphorylation sites. Nevertheless, these predictions must be treated with caution. Although the computational approach can, in principle, discriminate among closely related kinase paralogs, the similarity of their consensus motifs often renders such discrimination unreliable and of limited biological relevance.

Altogether, our results show that by selectively tuning host phosphorylation events, *Leishmania* restricts activation of potentially harmful host pathways while enabling those essential for successful infection, such as endocytosis, endosomal trafficking, GTPase signalling, and cytoskeletal reorganization. These processes are known to be crucial for the parasite entry and establishment within macrophages ^41–43^.

### *Leishmania donovani* is involved in the reduction of host protein abundances in a temporal manner

To determine whether *L. donovani* also modulates host protein abundances, we performed a proteomic analysis on the samples used for phosphoproteomics analysis, 4 hours post Zymosan phagocytosis, 4 hpi, 24 hpi and 48 hpi (Figure S3A). The corresponding volcano plots, representing the distribution of protein abundance changes across the four conditions, are shown in Figure S3 (see also Table S3). Figure 2A (see also Table S4) shows that the overall protein abundance changes only marginally during phagocytosis, indicating that this process exerts a minimal effect on the proteome. These results reinforce the hypothesis that the downstream consequences of phagocytosis are primarily mediated by signalling-dependent phosphorylation rather than by changes in protein abundance. Noticeably, at 24 hpi, the number of proteins exhibiting reduced abundance peaked at 1,247 (68% of total proteins, Figure 2A), which is a significant shift from 4hpi (196 down-regulated proteins, 44% of total proteins). This trend became even more pronounced at 48hpi, where 567 proteins showed a reduced abundance (94% of total proteins, Figure 2A, Inf 48h/UI 48h), indicating a progressive reduction of host protein abundance as infection progresses. This pattern likely reflects a parasite-driven, time-dependent reprogramming of host functions as the infection progresses. This early-stage decline in protein abundance is easier to interpret than the changes observed in the phosphoproteome, because a reduction in protein levels directly signals down-regulation of the corresponding pathways. To gain insights into the processes regulated by these host proteins, we conducted a metascape pathway enrichment analysis on the lists of proteins with downregulated abundance for all three infection conditions (Figure 2B, the complete metascape analysis is available in Fig. S4). Nearly half of the enriched pathways, including ‘endocytosis’, ‘Rho GTPase signalling’, and ‘regulation to biotic stimulus’, were consistently enriched across all infection timepoints (Figure 2B), as well as after zymosan phagocytosis (Figure S4B). in contrast, pathways including ‘membrane organisation’, ‘carbon metabolism’ and ‘mitochondrial protein degradation’, were specifically enriched at 24hpi and 48hpi, suggesting a parasite-specific regulation.

**FIGURE 2:**
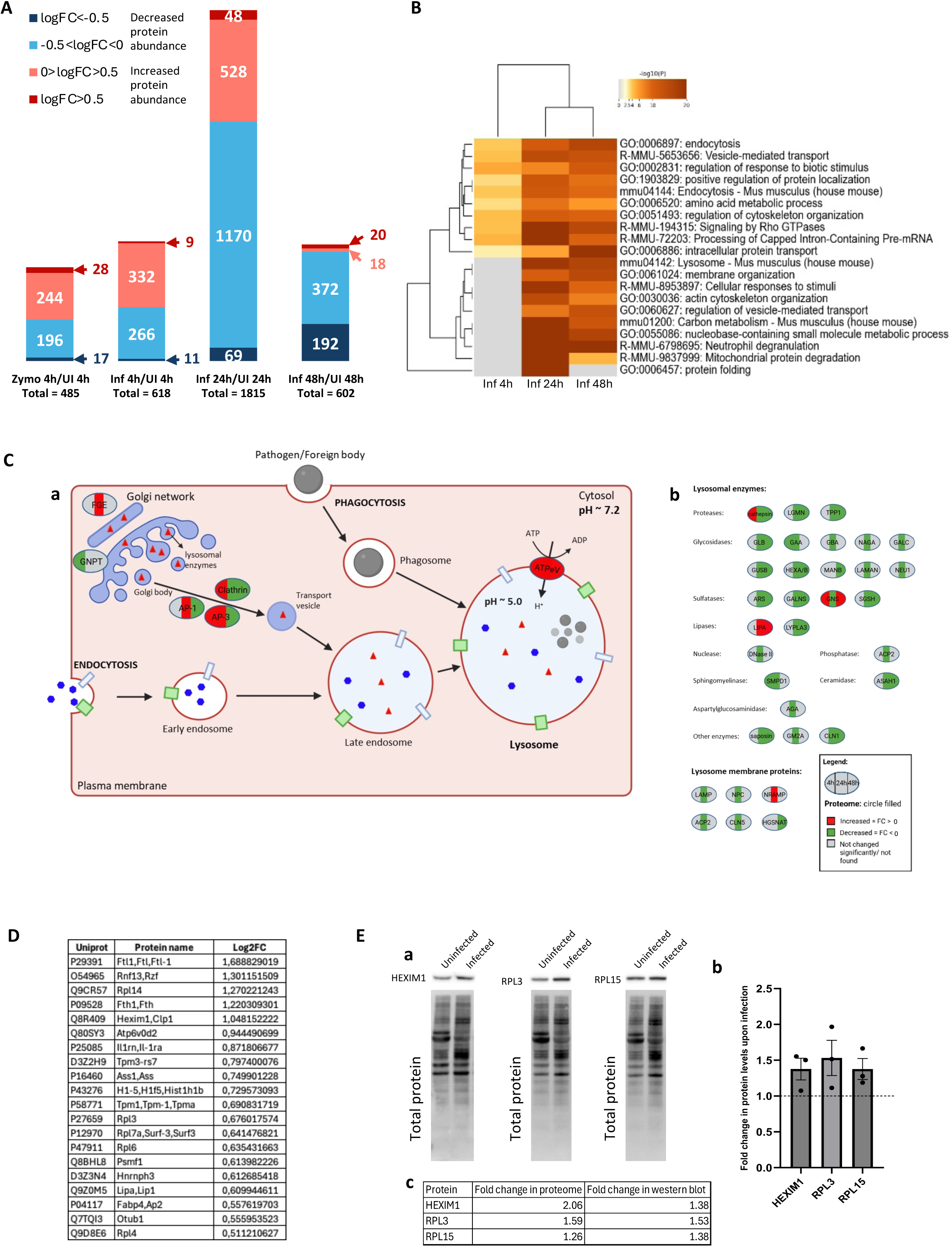
Modulation of host protein abundances by *L. donovani*. **A.** Distribution of fold changes in protein abundance (p<0.05, any FC, quantified in at least three replicates) is given for 4h Zymosan, 4 hpi, 24 hpi and 48 hpi. The total number of proteins is presented below each plot. The color code for the four brackets for fold change values is given in the legend. The number of proteins present in each of the fold change brackets is given below the plot. The proteome has been done in five replicates. **B.** Functional impact of the host proteome modulation by *L. donovani.* GO term enrichment of 4h Zymosan, 4 hpi, 24 hpi and 48 hpi for proteins with increased (Log2FC>0) and decreased (Log2FC<0) abundance. The intensity of orange provides the level of significance of the enrichment. Grey indicates that the pathway was not enriched. **C.** Modulation of protein abundances in the lysosome pathway during infection progression. A pathway illustration was adapted with BioRender based on the KEGG lysosome pathway (Pathway ID: mmu04142). Proteins were color-coded according to their expression status - Red: FC > 0, Green: FC < 0, Grey: Not detected in the dataset. Circle divided in three: 4 hpi, 24 hpi and 48 hpi. **D.** Table with the list of twenty proteins having FC>1.5 at 48 hpi. **E.** Validation of some of the highly abundant proteins by Western blot analysis using 48h uninfected and infected BMDM lysates. (a) Western blot images for HEXIM1, RPL3 and RPL15. (b) The total protein lane intensity was used for normalizing the corresponding band intensities before performing any further analysis. Fold changes in HEXIM1, RPL3 and RPL15 levels upon infection. Calculated from the Western blot by the ratio of infected/uninfected BMDMs. The graph represents three biological replicates shown as dots in each graph and the horizontal dotted line represents a ratio infected/uninfected of 1. The error bars represent standard error. (c) Table comparing the fold change values for HEXIM1, RPL3 and RPL15 in the proteomic analysis and western blot. The experiment was conducted with three biological replicates. Band intensities were normalized to total protein levels, and the ratio of normalized intensity between infected and uninfected samples was calculated.

### *Leishmania donovani* down-regulates host lysosomal enzyme abundance, thereby limiting parasite digestion

One striking example of coordinated protein down-regulations are lysosomal proteins. Although the parasite-containing PV fuses with host lysosomes ^44^, the parasite is not digested by lysosomal hydrolases. Our data might provide an explanation, as we mapped the proteomic data onto the lysosomal pathway, integrating protein modulation at 4hpi, 24hpi, and 48hpi (Figure 2C). Vesicles containing lysosomal enzymes are transported from the Golgi network to the late endosomes via proteins such as clathrin, adaptor proteins AP-1 and AP-3. These proteins were found to be upregulated until 24hpi and then downregulated at 48hpi (Figure 2C-a). Moreover, this mapping revealed a progressive decrease in the abundance of lysosomal enzymes: proteases, glycosidases, sulfatases, lipases, nuclease, phosphatase, sphingomyelinase, ceramidase, aspartyl-glucosaminidase, lysosomal membrane proteins from 24h to 48hpi (Figure 2C-b). Such a reduction of lysosomal hydrolases likely enables PV-lysosome fusion while minimizing lysosomal degradation of parasite components, ensuring intracellular persistence. Interestingly, at 48hpi, only two lysosomal enzymes were upregulated: lipase A (Lipa) and glucosamine (N-acetyl)-6-sulfatase (Gns). These enzymes may serve parasite-specific functions. Currently, there is no evidence linking Gns to *Leishmania* infection, making it an intriguing candidate for further investigation to determine its potential importance for the parasite. In contrast, previous studies have shown that *L. donovani* depends on host-derived cholesterol for successful infection ^45^.

Lipase A degrades the cellular lipoproteins and releases free cholesterol and fatty acids for further uses ^46^. Thus, upregulation of Lipa at 48 hpi may represent a strategic mechanism employed by the parasite to secure its cholesterol supply and maintain a favourable intracellular niche. Overall, our quantitative proteomics data indicate that rather than possessing intrinsic resistance to hydrolytic enzymes ^47, 48^, amastigotes might delay phagosome maturation to allow the degradation of these enzymes while preserving the acidification of the PV. The resulting acidic, hydrolase-depleted phagolysosome provides a permissive niche that supports parasite survival and replication.

### *Leishmania donovani* selectively up-regulates the abundance of a limited subset of host proteins at 48 h post-infection

The effect of *Leishmania* on host proteome is a massive decline in protein abundance. Consequently, the relatively few proteins that become more abundant are particularly outstanding. At 48 hpi, only 6 % of detected proteins are up-regulated, compared with 55 % at 4 hpi and 32 % at 24 hpi (Figure 2A). This pronounced shift suggests that *Leishmania* exerts tight, time-dependent control over the subset of host proteins whose levels are increased, implying that these molecules may be crucial for parasite survival. We identified only twenty proteins displaying a log2FC greater than 0.5 (Figure 2D). To experimentally validate the proteomic results, we selected three of these proteins: Hexim1, a nuclear protein that inhibits RNA polymerase II mediated transcription ^49^, Rpl3 and Rpl15, both structural components of the 60S ribosomal subunit, important for protein translation^50^. Western blotting was performed on whole-cell lysates from BMDMs infected or uninfected with *L. donovani* amastigotes at MOI 10 for 48 hours (Figure 2E-a). These quantifications (Figure 2E-b) corroborate the proteomic changes, supporting the validity of the dataset (Figure 2E-c, Table S4). We then carried out Gene Ontology enrichment analysis on this highly restricted protein set to identify the pathways that are regulated by these proteins. Compared to the 4-h and 24-h time points, where up-regulated proteins were associated with numerous pathways, such as immune response and vesicle-mediated transport (Figure S4C), at 48 hpi, up-regulated proteins were only significantly associated with ‘mRNA processing’, ‘neutrophil degranulation’ and ‘eukaryotic translation initiation’ (Figure S4C). This finding suggests that only a very limited subset of pathways remained active 48 hours post-infection, excluding GO terms such as ‘innate immune response’ or ‘cellular response to stimuli’. Overall, this limited number of proteins up-regulated during *Leishmania* infection might be essential for parasite survival in the host cell. To test this hypothesis, we selected host Ferritin light chain 1 (Ftl1), the protein exhibiting the highest fold-change (infected vs non-infected) at 48 h post-infection, for detailed functional characterization.

### *Leishmania donovani* actively sequesters host ferritin, a key player of iron homeostasis

Ferritin is a cytosolic macromolecule composed of 24 subunits that assemble into a heteropolymeric complex. The H-subunit (Fth1) provides the ferroxidase activity, while the L-subunit (Ftl1) promotes the incorporation and storage of iron within the ferritin core ^51^. Ferritin is the main intracellular iron-storage protein complex in cells and can sequester roughly 4,500 iron atoms ^52, 53^. Iron is a critical micronutrient for all living cells, essential for oxygen transport, energy metabolism, enzymatic catalysis, and immune regulation. Both iron deficiency and overload are detrimental for cells in general, and the effect is particularly pronounced for *Leishmania*, underscoring the necessity for tight regulation of iron homeostasis ^54^ ^55, 56^. Ferritin levels are regulated in response to cellular iron status, either through translational control mediated by iron response proteins ^57^ or via lysosomal degradation to mobilize stored iron ^58, 59^, thereby maintaining intracellular iron homeostasis.

To validate the ferritin upregulation observed in the proteomic analysis of mouse BMDMs at 48 hpi, we performed a Western blot analysis using whole-cell protein lysates of mouse BMDMs infected or uninfected with *L. donovani* amastigotes. Ftl1 levels were assessed at 4 and 48 hpi (Figure 3A). While Ftl1 level was not changed at four hours post infection compared to uninfected cells (FC 1.39, Figure 3B), a significant increase was observed at 48 hpi (FC 2.71, Figure 3B), confirming our proteomics result. To determine whether the observed increase in Ftl1 abundance was accompanied by changes in its subcellular localization, mouse bone marrow–derived macrophages (BMDMs) were either left uninfected or infected with *Leishmania donovani* amastigotes for 4 hours, 48 hours, or 6 days, followed by immunofluorescence staining with an anti-Ftl1 antibody. Ftl1 consistently localized around intracellular *L. donovani* amastigotes at all examined time points, whereas it exhibited a uniform cytoplasmic distribution in uninfected macrophages (Figure 3C-a & 3C-b uninfected and infected columns and Figure 3D). This is a long-lasting process, as quantitative analysis showed that over 80% of intracellular parasites were still surrounded by Ftl1 at 6 days post-infection (Figure 3D). Since *L. donovani* parasites reside inside PVs in macrophages, this localization could be a consequence of phagocytosis and not specific to *Leishmania* infection ^60^. To address this hypothesis, mouse BMDMs were treated with Zymosan particles for four hours, and Ftl1 localization was assessed by immunofluorescence. As judged Figure 3C-c, Ftl1 did not localise around Zymosan particles in any of the analysed cells, a finding that was consistently reproducible (see quantitative analysis, Figure 3D), indicating that the Ftl1 localisation is specific to *L. donovani* infection and not the consequence of phagocytosis. This localisation was also dependent on parasite viability, as only 10% of paraformaldehyde-fixed parasites retained Ftl1 accumulation in their periphery (Figure 3C-d & 3D). This finding indicates that Ftl1 recruitment to the parasite location is an active, dynamic, parasite-driven process during infection.

**FIGURE 3:**
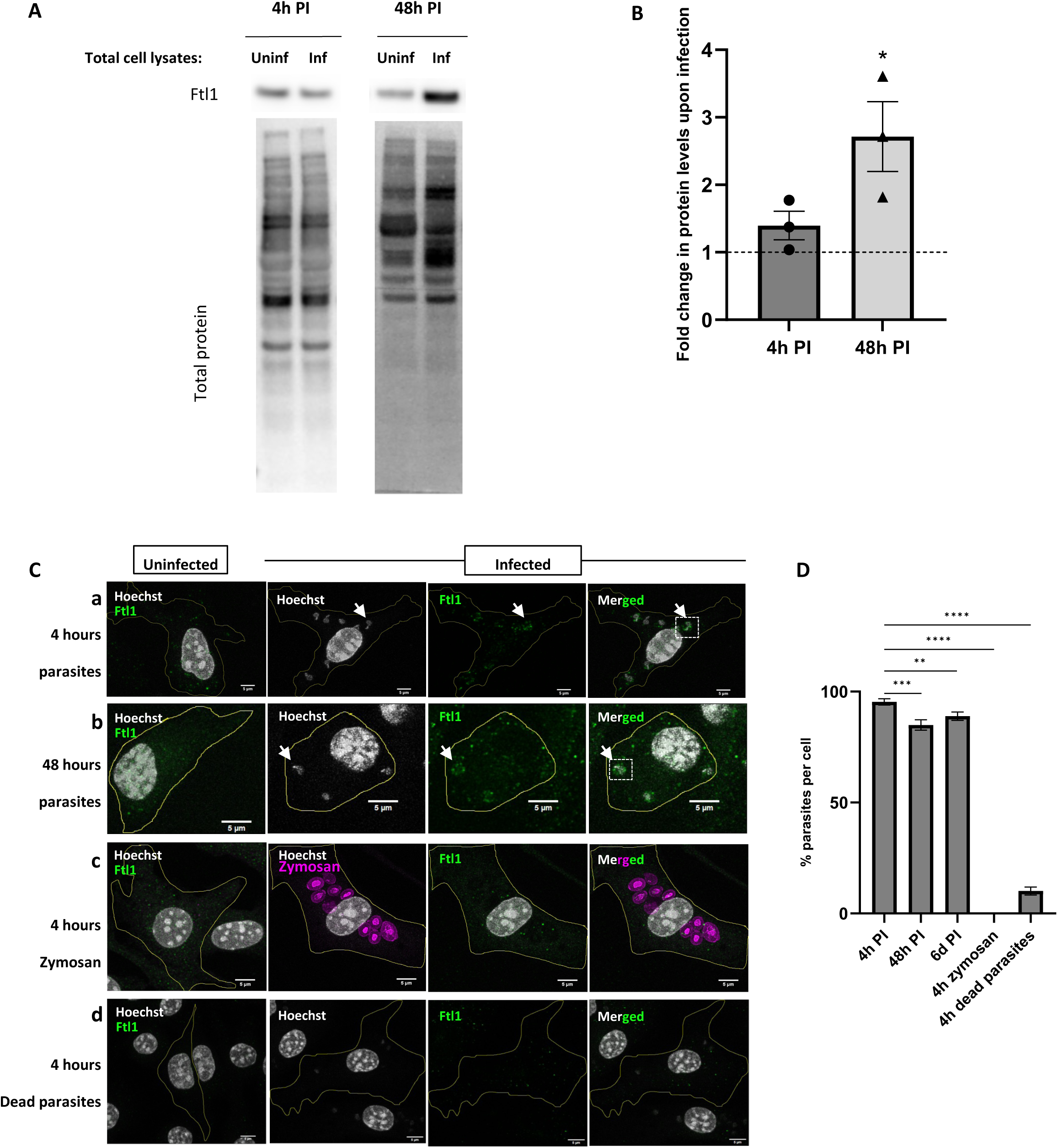
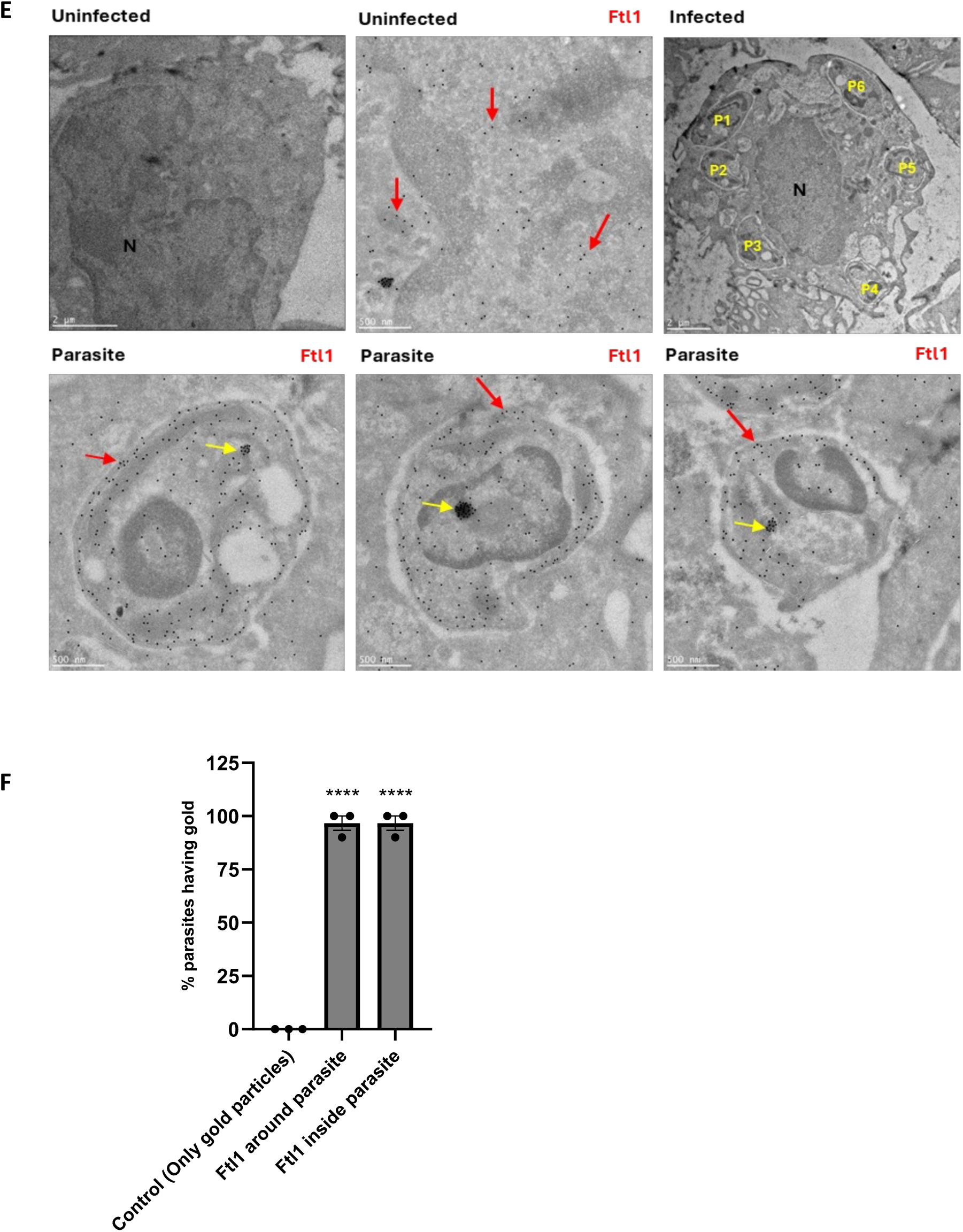
Ftl1 is upregulated at 48hpi and localises in the vicinity of the parasite. **A.** Western blot showing the Ftl1 levels in uninfected and infected cells at 4h and 48h PI. The total protein lane intensity was used to normalise the corresponding band intensities. **B.** Quantification of the fold changes in Ftl1 levels upon infection. The quantification has been done on three biological replicates. The horizontal dotted line represents a ratio infected/uninfected of 1. The statistical significance analysis was performed with respect to FC=1 for each timepoint. The error bars represent standard error. **C.** Representative immunofluorescence images showing the localisation of Ftl1 in uninfected and infected cells with Hoechst (grey) and Ftl1 (green) for 4hpi (a), 48hpi (b), 4h zymosan treatment (c) and 4hpi with dead parasites (d). The white arrows and box indicate the location of a parasite and shows Ftl1 localised around it at 4 hpi and 48 hpi. Scale bar, 5 µm. **D.** Quantification of % of parasites having Ftl1 localised around and inside them per cell. The analysis was performed by counting at least 100 cells, in three replicates. The error bars represent standard errors and the significance level between conditions is indicated. **E.** Representative electron microscopy images showing the presence of Ftl1 (black dots) in the macrophages. In the infected cell, six parasites labelled P1 to 6 can be seen. In the zoomed images, the red arrows indicate where Ftl1 is present. The presence of Ftl1 aggregates is also indicated by yellow arrows. Scale bar, 500 nm. **F.** Quantification of the percentage of parasites having Ftl1 around and inside for the control and the Ftl1 immunolabelling conditions. Thirty eight parasites were counted in total in three biological replicates. The error bars represent standard errors and statistical significance between conditions is indicated.

To determine whether Ftl1 localizes to the membrane of the parasitophorous vacuole, within the PV lumen, or directly at the parasite surface, we applied 4× expansion microscopy to visualise BMDMs infected or uninfected with *Leishmania donovani* amastigotes for 4 hours. Ftl1 was detected in the cytoplasm of uninfected and infected BMDM, at the parasite plasma membrane and within the parasite itself (Figure S5A). To confirm these localizations, we performed a Ftl1 gold immunolabelling followed by Electron Microscopy under the same infection conditions (Figure 3E, top panel: macrophages, bottom panel: zoom in on parasites; Figure S5B). Ftl1 did not seem to accumulate at the PV membrane but rather at the parasite plasma membrane itself (Figure 3E & S5B) in 100% of the imaged parasites across three biological replicates (Figure 3F). Ftl1 was also observed within all the observed parasites (Figure 7D). To further validate this finding, we purified *L. donovani* amastigotes from the liver and the spleen of a hamster, fixed them with 4% PFA and subjected them to immunofluorescence staining using the anti-Ftl1 antibody. Consistent with the electron microscopy data, Ftl1 was detected within 80% of isolated parasites (Figure S5C and S5D), confirming that the ferritin subunit is also transferred to the parasite *in vivo*. Notably, within the infected macrophages, aggregates of Ftl1 were observed in 37% of the parasites, in the cytoplasm (Figure 3E, bottom left panel, yellow arrow), nucleus (Figure 3E, bottom middle panel, yellow arrow) and kinetoplast (Figure 3E, bottom right panel, yellow arrow). Additional examples are provided in Fig. S5B. Consistent with prior reports that nuclear ferritin protects chromosomal DNA from reactive-oxygen-species–induced damage ^61, 62^, the dense ferritin aggregates we observed within the parasite nuclei (Fig. 3E, bottom middle panel, yellow arrow & Fig. S5B) might constitute a dedicated defence mechanism that mitigates macrophage-derived oxidative stress. Collectively, these data indicate that host-derived Ftl1 is translocated into the parasite, including its nucleus, not merely as a nutritive iron source but also to carry out dedicated functions, namely, to mitigate oxidative stress and protect DNA from ROS-induced damage.

To assess whether the entire ferritin complex, rather than only Ftl1, associates with the parasite, we localised Ferritin heavy chain (Fth1), the second subunit of the host ferritin heteropolymer in infected host cells. Mouse BMDMs were infected with *L. donovani* amastigotes for 48 hours followed by immunofluorescence staining using an anti-Fth1 antibody. We observed the localisation of Fth1 around 97.8% of intracellular parasites at 48 hpi (Fig. 4A & 4B). To validate this observation, we performed immunogold labelling of Fth1 in BMDMs infected with *Leishmania donovani* amastigotes for 4 h and examined the samples by transmission electron microscopy. Fth1 was detected both around the parasite and inside the parasites as shown in Figure 4C, red arrow. These findings demonstrate that both subunits of host ferritin complex are sequestered around and internalized by the parasite as early as four hours post infection.

**FIGURE 4:**
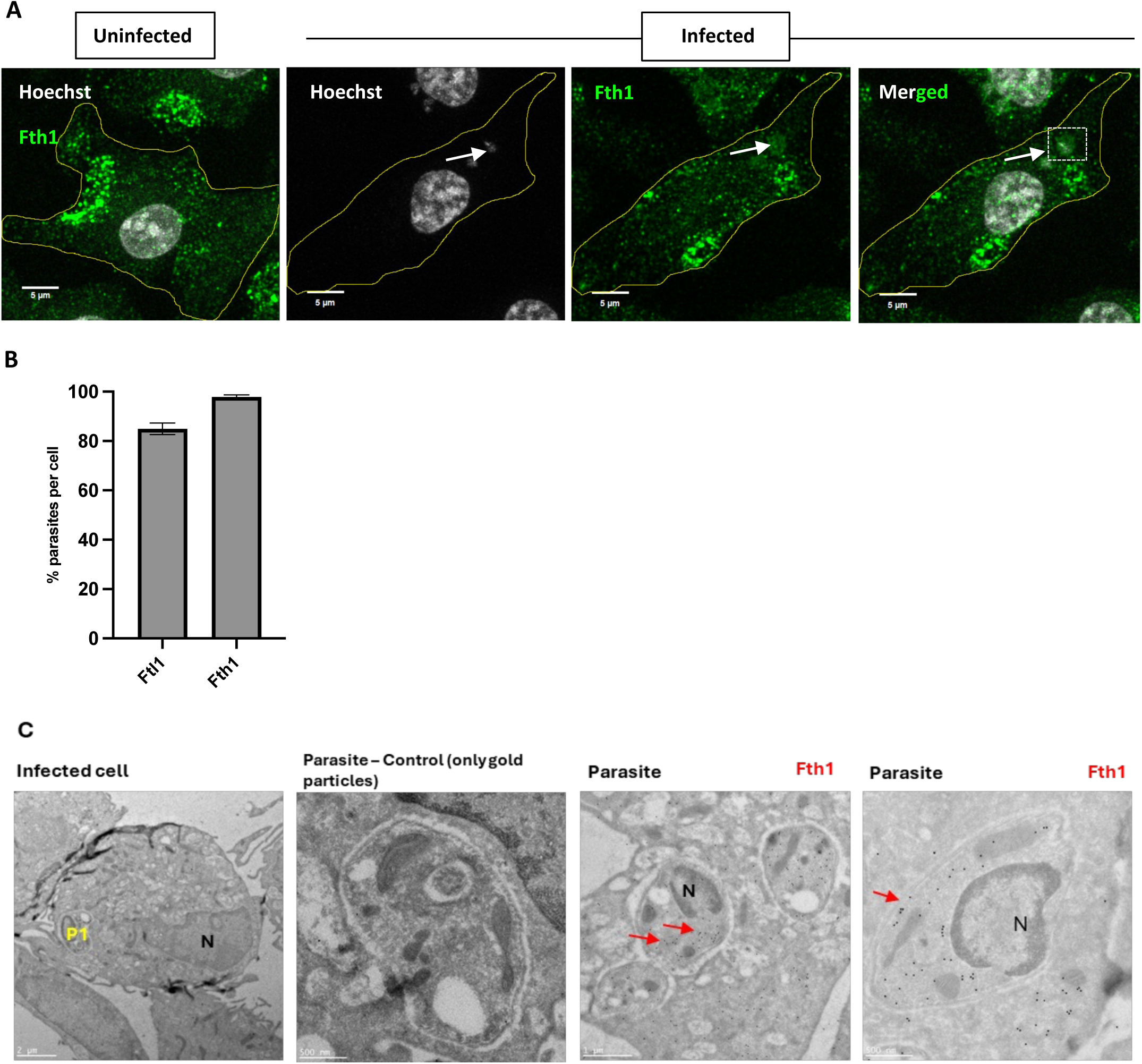
Fth1 localisation in the vicinity of parasites during infection. **A.** Representative immunofluorescence images showing uninfected and infected cells at 48 hpi with Hoechst (grey) and Fth1 (green). The white arrow and box indicate the location of a parasite and Fth1. Scale bar, 5 µm. **B.** Quantification of the % of parasites having Ftl1 and Fth1 around them per cell at 48 hpi. Quantifications were performed by counting around 100 cells in three biological replicates. The error bars represent standard error. **C.** Representative electron microscopy images for infected cells showing the presence of Fth1 inside the parasite (red arrows). Scale bar, 1 µm and 500nm respectively.

Altogether, these data demonstrate that host ferritin is actively recruited into the parasite in a parasite-driven manner, revealing ferritin as a host factor co-opted by *Leishmania donovani*.

### *Leishmania donovani*, but not *L. amazonensis* modulates host ferritin composition

The ratio of Ftl1 to Fth1 within the ferritin complex varies depending on the cellular environment ^63^. Cells in iron-rich organs such as liver and spleen typically exhibit a ferritin composition enriched in light chain subunits ^63, 64^. Since *L. donovani* predominantly resides in these organs ^65^, the parasite might affect Ftl1/Fth1 ratio in infected macrophages. To test this hypothesis, we conducted Western blot analyses examining the protein levels of Ftl1 and Fth1 in whole-cell lysates from BMDMs that were either uninfected or infected with *L. donovani* amastigotes for 48 hours (Western blot presented in Fig. S6A). The ratio Ftl1/Fth1 was calculated for each condition to assess the impact of infection on the composition of ferritin complexes. At 48 hpi, a significant six-fold increase of the Ftl1/Fth1 ratio (3.8±1.1) is observed in infected cells relative to uninfected controls (0.6±0.02), indicating a significant shift toward a light chain-rich ferritin composition (Figure 5A).

**FIGURE 5:**
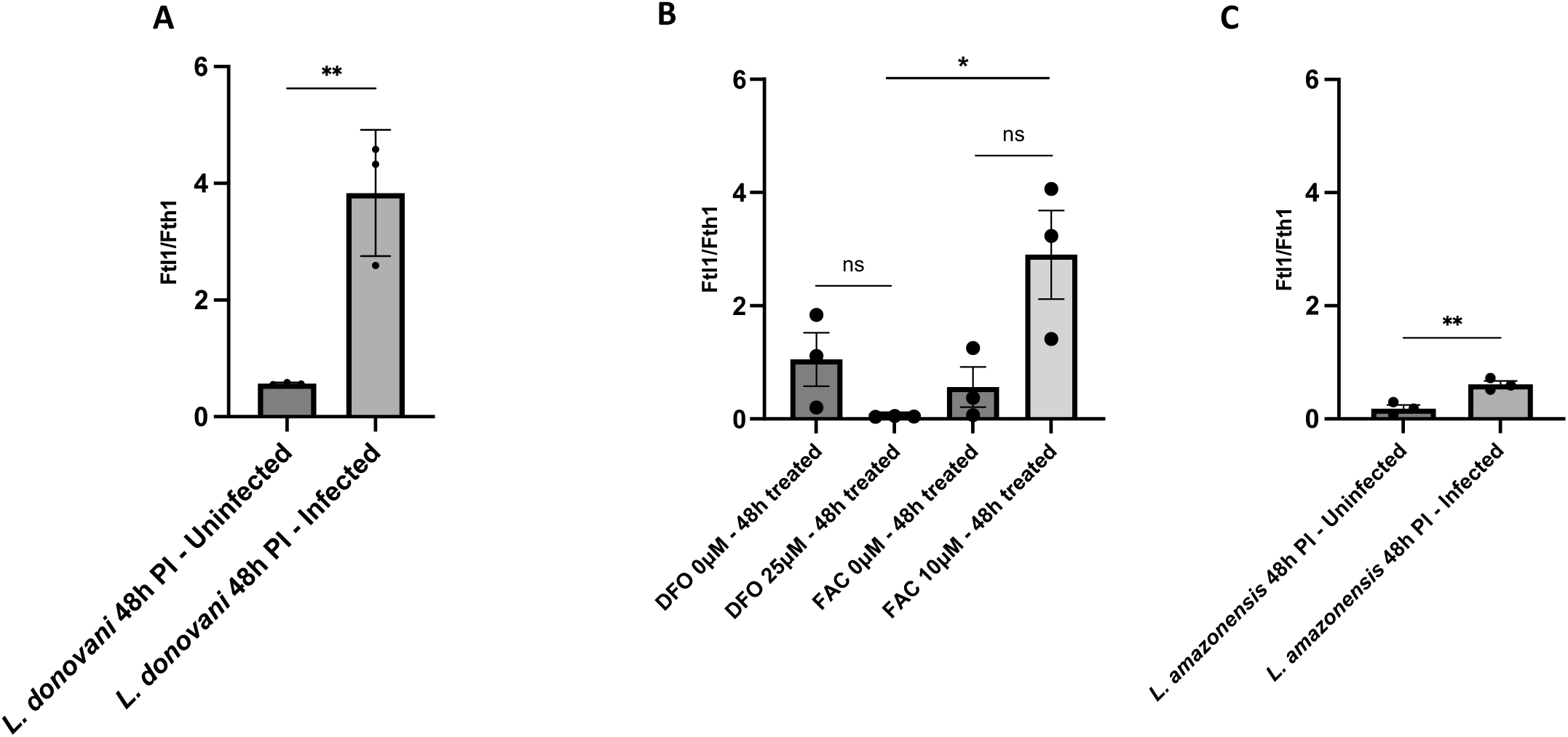
The Ftl1/Fth1 ratio is different between *L. donovani* and *L. amazonensis*. **A.** Quantification of Ftl1/Fth1 ratio for BMDMs infected or not with *L. donovani* for 4 hpi and 48 hpi. **B.** Quantification of Ftl1/Fth1 ratio for BMDMs treated or not with 25 µM DFO or 10 µM FAC for 48 hpi. **C.** Quantification of Ftl1/Fth1 ratio for BMDMs infected or not with *L. amazonensis* for 48 hpi. All quantifications were performed with three biological replicates and the statistical significance between conditions is indicated with an asterisk: below 0.05 (*), below 0.01 (**) and below 0.001 (***). The error bars represent standard error.

Altering iron concentration also drives a change in the ferritin composition, as demonstrated by the following experiment. We treated mouse BMDMs for 48 hours with either 10 µM ferric ammonium citrate (FAC) to simulate iron-rich conditions or 25 µM deferoxamine mesylate (DFO) to deplete iron and then subjected the whole cell lysates to Western blotting analyses using anti-Ftl1 and anti-Fth1 antibodies (Figure S6B). In presence of DFO, Ftl1/Fth1 ratio decreases markedly from 1.1±0.8 to 0.04±0.006, suggesting a predominance of Fth1-rich ferritin complexes. Conversely, treatment with FAC increased this ratio from 0.6±0.6 to 2.9±1.4, resulting in Ftl1-rich ferritin complexes, mirroring the changes observed in *L. donovani*-infected cells (Figure 5B & S6B). There is a significant difference between the Ftl1/Fth1 ratio in absence (+ DFO) or in presence of iron (+ FAC, p-value < 0.05, *, Figure 5B), indicating that iron supplementation with FAC creates an iron-rich extracellular environment similar to that of the liver and spleen, which in turn alters the Ftl1/Fth1 ratio. These findings suggest that *L. donovani* might actively modulates host ferritin composition to mimic the cellular iron-storage profile characteristic of an iron-rich environment, even in the absence of external iron stimuli. This mechanism might be a pre-adaptation strategy of the parasite to its future localization in the liver and spleen. If true, *Leishmania amazonensis*, a parasite species that causes cutaneous leishmaniasis and resides in the skin, a tissue poor in iron, should not increase the Ftl1/Fth1 ratio of BMDMs above 1 ^66^. To test this hypothesis, BMDMs were left uninfected or infected with *L. amazonensis* amastigotes. At 48 hours post-infection, the protein levels of Ftl1 and Fth1 were assessed and the ratio calculated (Figure 5C & S6C). Unlike *L. donovani*, in BMDM infected with *L. amazonensis*, the composition of ferritin remained heavy chain-rich as the Ftl1/Fth1 ratio increased from 0.2±0.1 to 0.61±0.1. Given that all other experimental conditions, including the use of mouse BMDMs and infection protocols, were identical, the observed differences are due to the parasite itself. These findings demonstrate that *Leishmania* preadapts host cells by modulating ferritin composition to match the iron levels of its future target organ, even in the absence of external iron cues.

### Ncoa4 is recruited to the intracellular parasite and colocalises with Ftl1

Our findings reveal that ferritin is actively recruited to the parasite during infection. Given that the parasite resides within the PV, ferritin must first be trafficked to this lysosome-like compartment ^67^. This observation raises a fundamental question: what host-driven mechanism mediates ferritin recruitment to the PV? Ferritin is normally targeted to lysosomes for degradation, particularly under low cytoplasmic iron conditions, to facilitate iron release into the cytosol ^58^. *Leishmania* may exploit this host degradation pathway to redirect ferritin to the parasitophorous vacuole. A key protein involved in this process is nuclear receptor coactivator 4 (Ncoa4), which binds to and delivers ferritin to lysosomes for degradation ^68, 69^. The analysis of Ncoa4 levels during infection, revealed a significant upregulation of the protein at 48 hours, but not at 4 hours post-infection (Figure 6A). Quantitative analysis revealed a two-fold increase in Ncoa4 abundance at 48 hours post-infection in infected macrophages (2±0.4) compared to uninfected controls (1±0.08, Figure 6B). The upregulation of both Ftl1 and Ncoa4 during infection, despite their typical inverse relationship ^70^, together with the known interaction between Ncoa4 and ferritin ^69^, supported the hypothesis that Ncoa4 may also localise in the vicinity of the parasites. To test this hypothesis, we examined the subcellular localization of Ncoa4 during infection. Immunofluorescence analysis of mouse BMDMs infected with *L. donovani* amastigotes revealed that Ncoa4 accumulates around the parasite at 4 hpi (Figure 6C-a, living parasites & Figure S7), 48 hpi (Figure S7) and 6 days post infection (Figure S7), mirroring Ftl1 localization. A quantitative colocalization analysis was performed at 4 hpi using Huygens Professional software to determine the pixels from Ftl1 and Ncoa4 that colocalise (Figure 6C-b, ROI, light green staining). A strong positive correlation was observed between Ncoa4 and Ftl1 at 4 hpi (Pearson’s coefficient > 0.5, Figure 6C-b live parasites), persisting up to 6 days post-infection (Figure 6C-c). Given our earlier observation that dead parasites fail to recruit Ftl1, we investigated whether Ncoa4 recruitment was also dependent on parasite viability. Interestingly, Ncoa4 still localized around 98% of dead parasites (Figure S7A), despite the absence of colocalization with Ftl1 (Figure 6C, dead parasites, Pearson’s coefficient = 0.348), suggesting that Ncoa4 recruitment is independent of that of Ftl1, and unlike Ftl1, may occur through a mechanism independent of active parasite signalling. Since Ncoa4 appears to be recruited independently of Ftl1 and parasite viability, we asked whether this could be a general response of phagocytosis. To test this hypothesis, mouse BMDMs were treated with Zymosan particles for 4 hours and Ncoa4 localization was assessed via immunofluorescence. Ncoa4 did not accumulate around Zymosan particles (Figure S7B), indicating that its recruitment to the PV is linked to the presence of the dead or alive parasite but is not due to phagocytosis.

**FIGURE 6:**
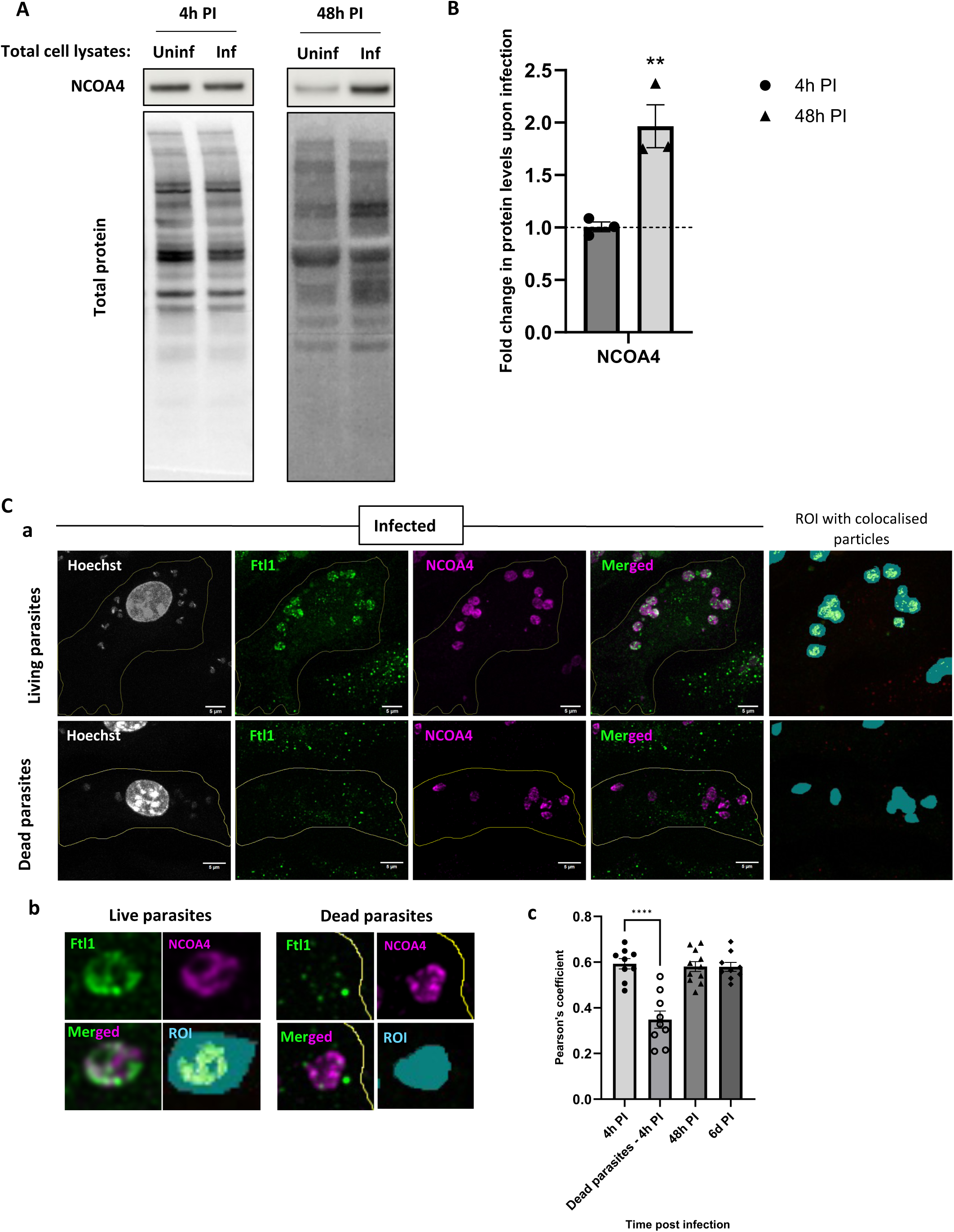
NCOA4 protein is up-regulated and localises in the vicinity of the parasite during infection. **A.** Representative Western blot of NCOA4 levels in uninfected and infected BMDMs at 4hpi and 48hpi. The total protein lane intensity was used to normalise the corresponding band intensities for quantification. **B.** Quantification of NCOA4 levels at 4h and 48h PI comparing infected/uninfected BMDMs. The horizontal dotted line represents a ratio infected/uninfected of 1. The statistical significance analysis was assessed compared to a ratio infected/uninfected of for each timepoint. **C.** Co-localisation analysis for Ftl1 and NCOA4. (a) Representative immunofluorescence images showing Hoechst (grey), Ftl1 (green), NCOA4 (magenta) and ROI with colocalised particles for 4h PI with live and dead parasites. (b) Representative parasites showing the Ftl1, NCOA4 localisation and the ROI with colocalised particles for live and dead parasites. (c) Pearson’s coefficients for 4 hpi, 24 hpi, 48 hpi and 4 hpi dead parasites. All the quantifications were performed with three biological replicates. The error bars represent standard errors. Scale bar, 5 µm.

Our findings demonstrate that *L. donovani* infection induces directly or indirectly the upregulation and recruitment of host Ncoa4 to the vicinity of the parasites, where it co-localizes with Ftl1. The accumulation of both proteins is unexpected as under normal conditions, stabilized Ncoa4 facilitates ferritin degradation via lysosomal trafficking, leading to a reduction in ferritin levels ^68, 69^. Conversely, the accumulation of ferritin in response to elevated cytosolic iron depends on the degradation of Ncoa4^68,70^. Their co-localization at a lysosome-like site would typically reinforce this outcome. However, we observe ferritin accumulation instead, suggesting that despite the recruitment of ferritin to the parasitophorous vacuole for degradation, this process might be compromised.

### *Leishmania* prevents the recruitment of NCOA4 partners involved in ferritin degradation

To verify this hypothesis, we investigated whether the three pathways known to be involved in Ncoa4-mediated degradation of ferritin were functional ^71^. First, ferritinophagy is a canonical autophagy process, where Ncoa4-ferritin complex is sequestered into autophagosomes and fuse with lysosomes leading to the degradation of ferritin ^69^. To assess whether this pathway is active during infection, we analysed electron microscopy images of infected macrophages for evidence of autophagosome-like membrane structures surrounding Ftl1 near the parasites or anywhere in the macrophage. No such structures were observed (Figure 7A and 3E, 4C, S5B), suggesting that canonical ferritinophagy is not functional during *Leishmania* infection. The second pathway is a non-canonical, autophagy-independent mechanism in which NCOA4-ferritin complexes are directly delivered to lysosomes via proteins, such as Atg9A, Vps34, and the ESCRT machinery ^72^. Immunofluorescence staining of Atg9A was performed on mouse BMDMs infected with *L. donovani* amastigotes for 48 hours. Atg9A was not detected in the vicinity of the PV (Figure 7B-a), and colocalisation analysis with Ncoa4 yielded a low Pearson coefficient of 0.336 (Figure 7C), indicating minimal spatial overlap and suggesting limited interaction between the two proteins during infection. The final known mechanism for ferritin degradation occurs in response to iron repletion. In the early phase of iron repletion, iron binds to Ncoa4 leading to the formation of condensates, which transiently sequester Ncoa4 away from ferritin. With prolonged repletion, Ncoa4 condensates interact with Tax1bp1 to deliver ferritin to lysosomes via a Tax1bp1 binding-dependent, noncanonical autophagy pathway, preventing relative iron deficiency caused by excessive storage ^73^. If this pathway were active during infection, Ncoa4 and Tax1bp1 would be expected to colocalize near the parasite. However, Tax1bp1 was not detected at the PV (Figure 7B-b) and did not colocalise with Ncoa4, as judged by the low Pearson coefficient of 0.347 (Figure 7C), indicating that this pathway is not activated during infection. Beyond the three pathways highlighted above, our proteomic data revealed an Ncoa4-independent mechanism that also contributes to ferritin stabilization. Ferritin is normally degraded by lysosomal enzymes^74^ ; however, we observed a pronounced decline in the abundance of these lysosomal hydrolases between 24 h and 48 hpi (Figure 2C). The reduced lysosomal proteolysis of ferritin likely underlies the ferritin accumulation we observe after 24 h post-infection.

**FIGURE 7:**
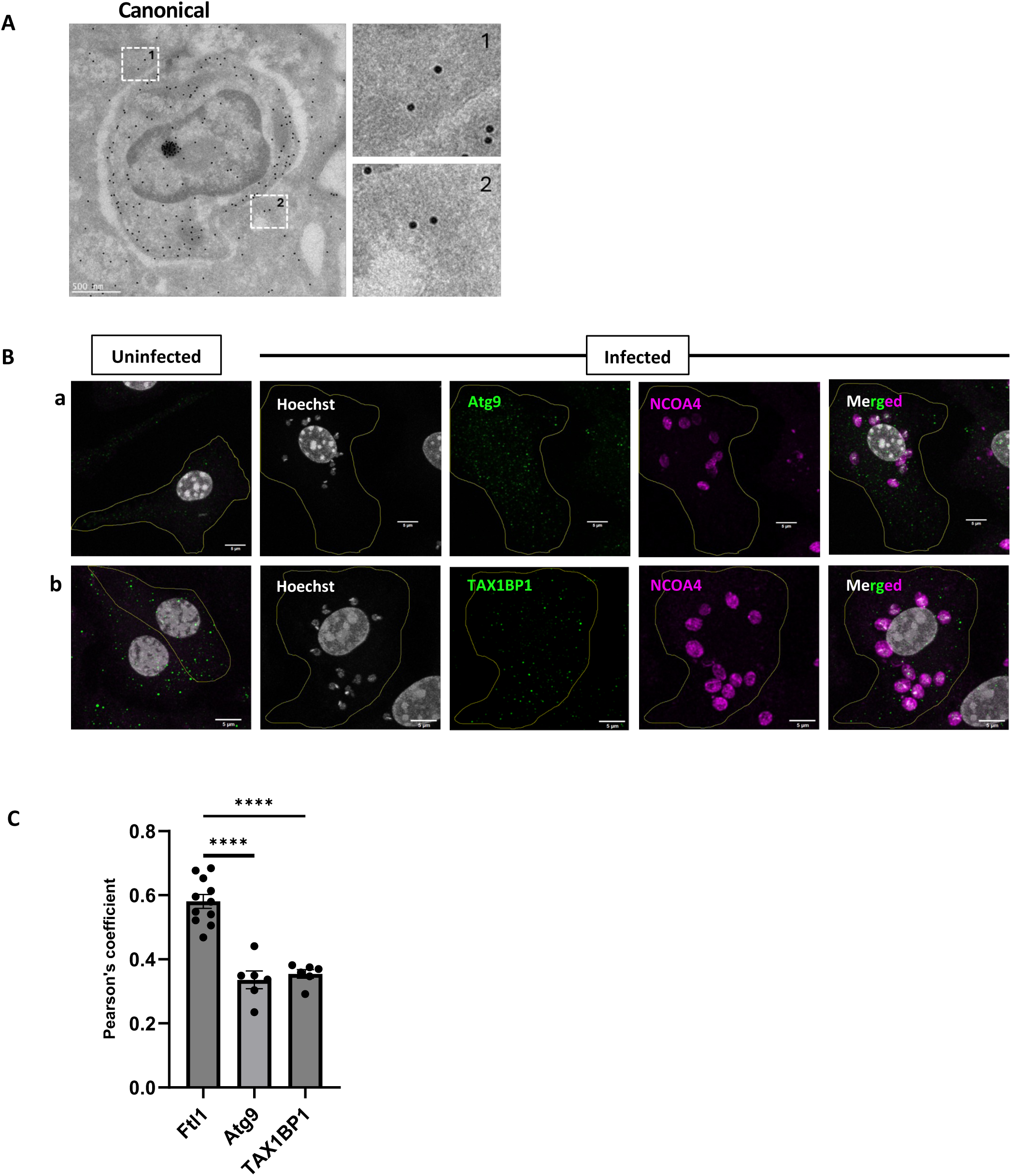
Assessing the ferritin degradation pathways are used during infection. **A.** Canonical autophagy: Representative electron microscopy image of the lack of membrane around Ftl1 as seen in fields 1 and 2 represented by white boxes and the corresponding zoomed view. Scale bar, 500 nm. **B.** Representative immunofluorescence images of Atg9 and TAX1BP1 colocalization with NCOA4. (a) Localisation of Atg9 (green) and NCOA4 (magenta) in uninfected and infected cells. (b) Localisation of TAX1BP1 (green) and NCOA4 (magenta) in uninfected and infected cells. Scale bar, 5 µm. **C.** Pearson’s coefficient for colocalization of NCOA4 with Ftl1, Atg9 and TAX1BP1. The quantifications were performed with three biological replicates and the error bars represent standard errors. The statistical significance is indicated.

Taken together, our findings indicate that none of the known Ncoa4-dependent or -independent ferritin degradation pathways active in mammalian cells are engaged during *Leishmania* infection. Instead, ferritin, by binding to Ncoa4, is redirected to the parasitophorous vacuole, while the associated degradation mechanisms are uncoupled, and the lysosomal hydrolases degraded. This decoupling may represent a novel strategy by which *L. donovani* subverts host iron homeostasis to establish a permissive intracellular niche for its survival.

### The knockdown of *Ftl1* does not affect *Leishmania* survival in the macrophage

To investigate the functional role of ferritin on iron homeostasis and parasite survival during *L. donovani* infection, we knocked down *Ftl1* gene in mouse bone marrow-derived macrophages using siRNA. Mouse BMDMs were treated with either a universal negative siRNA control or a Ftl1 targeting siRNA for 4 hours, and the treated cells remained uninfected or were infected with *L. donovani* amastigotes. The knockdown of Ftl1 was validated 48-hours post-treatment at the transcript (Figure S8A) and the protein levels (Figure S8B). Treated BMDMs were fixed at 4 hours, 48 hours and 6 days post infection, and infection efficiency and progression were quantified by calculating the percentage of infected cells at these timepoints (Figure 8A) and counting the number of parasites per hundred BMDMs (Figure 8B). The percentage of infection and the parasite burden of the si-*Ftl1*-treated cells were similar to those of the si-control conditions at any timepoint. These results indicate that knockdown of *Ftl1* does not impair infection, at least during the first 6 days, suggesting that the parasite, to sustain infection, has likely evolved compensatory mechanisms to tolerate the loss of *Ftl1*. This hypothesis is consistent with the critical role of iron in parasite survival, serving as an essential nutrient for *Leishmania* while posing a risk of cell death if in excess ^75, 76^.

**FIGURE 8:**
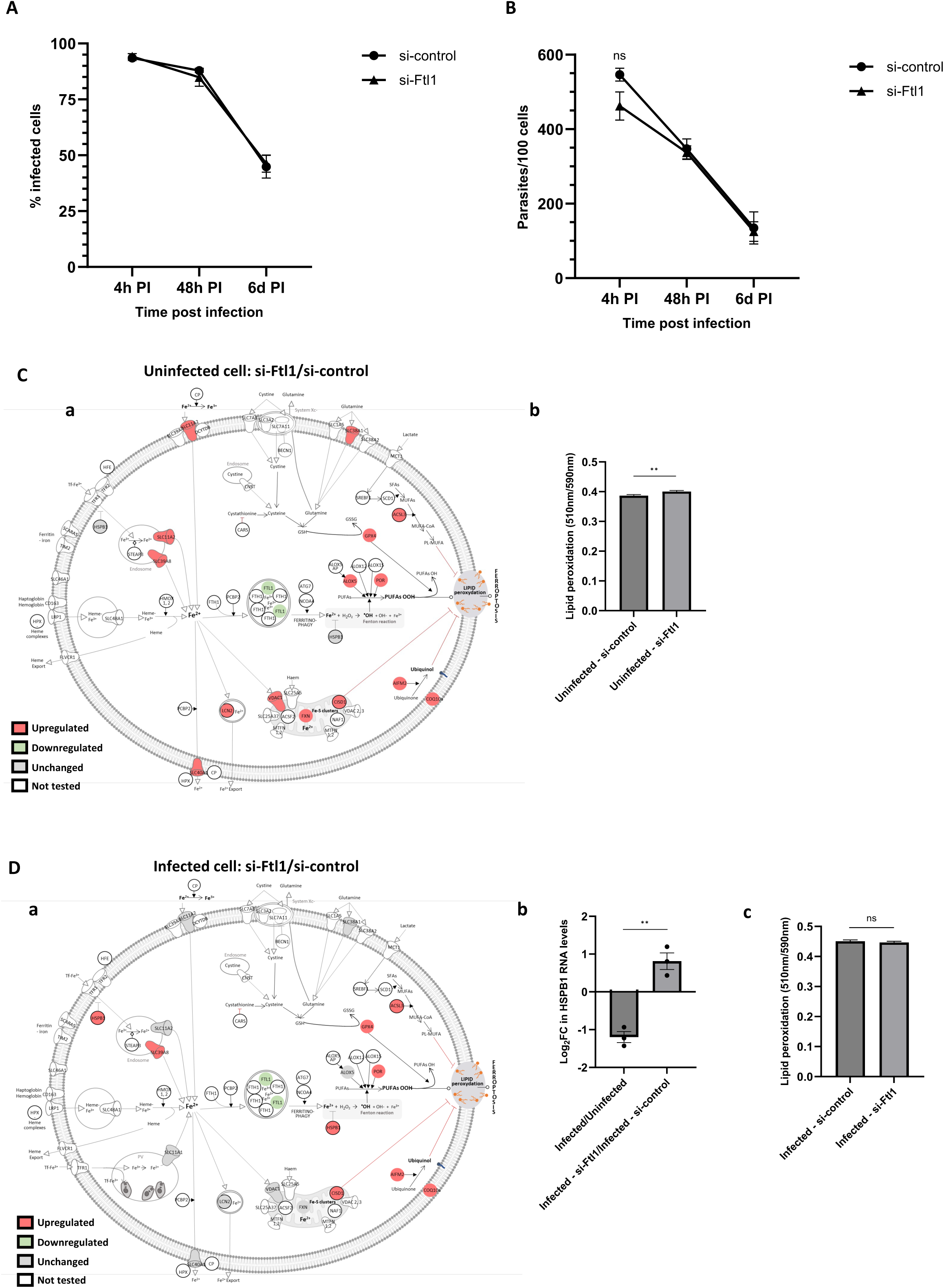
Impact of Ftl1 knockdown on *Leishmania* infection and iron homeostasis pathway. **A.** Percentage of infected cells at 4h, 24h and 48 hpi for the negative siRNA control (si-control) and Ftl1 siRNA (si-Ftl1). **B.** Quantification of parasite burden (number of parasites per 100 cells) at 4h, 24h and 48hpi for the negative siRNA control (si-control) and Ftl1 siRNA (si-Ftl1). All quantifications were performed with three biological replicates by counting around 300-400 cells in each condition. The error bars represent the standard deviation. **C.** In uninfected cells at 48h post siRNA treatment, (a) RT-qPCR analysis of genes in the ferroptosis pathway. The differential expression is shown using a color-coded illustration: red for significantly upregulated targets (Log2FC including error bars > 0), green for significantly downregulated targets (Log2FC including error bars < 0), and grey for unchanged targets (error bars encompassing zero). (b) Quantification of the lipid peroxidation level calculated using BODPIY C11 staining for si-control and si-Ftl1. C) In infected cells 48h hours post siRNA treatment, (a) RT-qPCR analysis of genes in the ferroptosis pathway. The differential expression is shown using a color-coded illustration: red for significantly upregulated targets (log2FC including error bars > 0), green for significantly downregulated targets (log2FC including error bars < 0), and grey for unchanged targets (error bars encompassing zero). (b) HSPB1 gene expression levels during infection and upon Ftl1 KD in infected cells. (c) Quantification of the lipid peroxidation level calculated using BODIPY C11 staining for si-control and si-Ftl1. All quantifications were done using three biological replicates. The error bars represent standard deviations.

### *Leishmania* has evolved multiple parallel mechanisms to regulate iron homeostasis

Ferritin, by sequestering excess intracellular iron, plays a key role in preventing increase of free cytosolic ferrous iron, which can trigger iron-dependent oxidative stress and activate ferroptosis, a regulated form of cell death characterized by iron-induced lipid peroxidation ^77^. This sequestration of iron is also used to prevent pathogens from accessing freely the pool of intracellular iron. We showed that *Leishmania* exploits this defence mechanisms to its advantage by recruiting the ferritin first to the PV and ultimately inside the parasite. Given the critical role of ferritin in iron homeostasis and the relevance of iron to *Leishmania* survival, we next explored potential compensatory mechanisms employed by the parasite to mitigate the effects of *Ftl1* loss. To this end, we selected 16 genes representing various aspects of the iron homeostasis pathway (Table S5) and assessed their expression by RT-qPCR. To first understand the basal impact of Ftl1 deficiency on uninfected mouse BMDMs, we compared si-*Ftl1* to si-control in uninfected BMDMs (Figure 8C-a & S8C). In these cells, *Ftl1* knockdown led to an increased expression of genes involved in iron transport (*Slc40a1*, *Slc11A2* and *Lcn2*), mitochondrial iron flux and storage (*Vdac1*, *Cisd1*, *Fxn*) and genes promoting lipid peroxidation (*Slc38a1*, *Alox5*, *Por*), which might lead to an increase in intracellular iron and ROS levels (Table S5). Several inhibitors of ferroptosis such as *Gpx4* which detoxifies peroxidised lipids; *Acsl3* which replaces polyunsaturated fatty acids (PUFA) with monounsaturated fatty acids (MUFA) to reduce the susceptibility of plasma membrane lipids to oxidation; and *Aifm2* and *Coq10a*, which inhibit lipid peroxidation, were also upregulated likely in response to the increased ROS stress. These findings indicates that upon *Ftl1* depletion, both positive and negative regulators of lipid peroxidation are upregulated in the uninfected macrophage. Our data suggest that *Ftl1* depletion, by triggering an increase of labile iron pool, activates pro–lipid-peroxidation pathways, thereby driving lipid peroxidation. In parallel, the uninfected macrophage mount compensatory antioxidant responses, upregulating lipid-peroxidation suppressors. To assess how these compensatory pathways impact lipid peroxidation observed in uninfected macrophages, uninfected si-control and si-Ftl1 cells were stained with the BODIPY™ 581/591 C11, and the 510/590 nm fluorescence ratio was compared between the two conditions. For each image, cells were segmented using Ilastik and these segmentation masks were then used to calculate the intensity ratios using Fiji. As shown in Figure 8C-b, we detected a slight, but statistically significant increase in lipid peroxidation level 48 hours after *Ftl1* siRNA treatment, indicating that the compensatory mechanisms are only partially effective.

Next, we examined whether *Ftl1* knockdown affects the regulation of iron homeostasis in *L. donovani*-infected BMDMs. To this end, we compared infected BMDMs treated with si-*Ftl1* or with the si-control using RT-qPCR for the same 16 targets (Table S5). In infected macrophages, about fifty percent of the tested targets, including iron exporters (*Slc40a1*, *Lcn2* and *Slc11a1*), mitochondrial iron flux (*Vdac1* and *Fxn*), showed no significant changes in the absence of *Ftl1* compared to the control (Figure 8D-a & S8D). These results are consistent with the tight regulation of iron homeostasis pathways observed during *Leishmania* infection, including the downregulation of *Slc40a1* expression, the host iron exporter ^78^. Noticeably, while iron export is upregulated in uninfected macrophages, its expression is still down regulated in infected cells, likely reflecting the parasite absolute requirement for intracellular iron. This observation further suggests that *Leishmania* activates alternative mechanisms to compensate for *Ftl1* loss and mitigate cytosolic iron accumulation as well as its deleterious consequences, without resorting to iron export. Among the pro-oxidant genes analysed, only *Por* was upregulated in the absence of *Ftl1*, while *Alox5* and *Slc38a1* levels remained unchanged, suggesting that *Leishmania* may be unable to alter the regulation of *Por* expression (Figure 8D-a).

Strikingly, in the absence of *Ftl1*, the only gene upregulated in infected macrophages, but not in uninfected controls, was *Hspb1* (Figure 8D-a). The encoded HSPB1 protein inhibits erastin-induced ferroptosis potentially by interfering with ROS production via the Fenton reaction ^79^, indicating that the parasite actively engages host defences to counteract oxidative stress. Interestingly, *Hspb1* is downregulated during normal infection (Figure 8D-b), likely because of its role in reducing cellular iron uptake ^80^ that could hinder a process essential for parasite proliferation. We next assessed the impact of *Ftl1* knockdown on lipid peroxidation using BODIPY™ 581/591 C11 staining. Lipid peroxidation levels remained unchanged in both conditions (Figure 8D-c), indicating that parasite-driven compensatory mechanisms are sufficient to prevent oxidative damage under these conditions, unlike in uninfected macrophages.

## Discussion

Although extensive work has documented the interference of *Leishmania* with key signalling cascades, including MAPK ^81^ ^2^, JAK-STAT ^82, 83^, and PKC ^84^, a comprehensive, global and temporal view of the host phosphoproteome during infection has been lacking, preventing access to the regulation of host signalling by *Leishmania*. By combining quantitative phosphoproteomics with total-proteome profiling, we demonstrated that *Leishmania* infection elicits a rapid, time-dependent reshaping of the macrophage phosphoproteome that far exceeded any alterations in protein abundance. Thus, phosphorylation emerges as the main regulatory switch exploited by the parasite to reshape host signalling networks and to block host cell defence mechanisms.

The swift remodelling of the macrophage phosphoproteome observed during *Leishmania donovani* infection is likely achieved through at least three distinct strategies. First, *L. donovani* manipulates host kinases directly or indirectly. Our phosphoproteomic profiling, obtained using the framework established by Johnson *et al.*, suggests that *Leishmania* does more than merely perturb substrate phosphorylation, it seems to hijack host kinases, especially those that drive pro-inflammatory signalling, by blocking their normal activation (e.g., MAPKs) or by preventing their inactivation (e.g., IRAK4) during phagocytosis, consistent with earlier observations ^81, 85, 2^. This dual interference locks these signalling enzymes in a basal, “uninfected” state, thereby blunting the host innate immune response. Second, *Leishmania* directly modulates host signalling proteins, through the release of extracellular vesicles (EVs) by the parasite. These EVs have been shown to contain at least twenty-three kinases and five phosphatases that can be delivered into the host cytoplasm and modified host substrates ^6, 21, 23, 24, 86, 87^. For example, we and Liu *et al.* demonstrated that the *Leishmania*-derived kinase CK1.2 directly phosphorylates host substrates ^88, 89^, including the host kinases Rps6ka1, Slk and Tnik, whose activities were differentially regulated in infected macrophages in the present study. This demonstrates that *Leishmania* can directly re-wire the signalling network of specific host enzymes^89^. Third, *L. donovani* directly activates host phosphatases^90^, suppressing multiple immune-signalling kinases and ultimately dampening the host inflammatory response ^81, 91^. Our quantitative data also reveal that protein tyrosine phosphatase 1B (Ptp-1B) and the serine/threonine phosphatase Pp2a are up regulated in macrophages following *Leishmania* infection. Both enzymes have been implicated in the dampening of NF κB driven transcription and the dephosphorylation of central signalling hubs such as Erk1/2, thereby further impairing the host immune response ^19^.

In contrast to the widespread phospho-rewiring that occurs during infection, changes in the overall abundance of host proteins are modest and follow a progressive decline. This progressive loss of many host proteins likely reflects two non-exclusive mechanisms. First, targeted proteolysis contributes to the depletion of specific factors such as Stat1^82^ and Ifnar1^88^, suggesting that *Leishmania* actively accelerates protein turnover to blunt host defence pathways. Second, the parasite suppresses de novo host protein synthesis, as shown by our proteomic analysis and corroborated by previous reports. Indeed, *Leishmania* selectively modulates the mTor pathway ^92, 93^ to inhibit translation. GP63, the major *Leishmania* secreted protease, cleaves mTorc1, thereby preventing 4e-bp1 phosphorylation and thus suppressing cap-dependent translation^90^. Our quantitative proteomics confirmed a marked down-regulation of mTor protein levels, reinforcing the view that *Leishmania* blocks host protein synthesis.

This coordinated reduction in host protein abundance, potentially through degradation and inhibition of translation, complements the parasite-driven phospho-reprogramming of the host cell, together constituting a strategy to attenuate antimicrobial defences. In this context, it is striking that only a very small subset of proteins is up-regulated ≥ 2-fold. These outliers are likely host factors that the parasite deliberately co-opts to facilitate its own survival. We selected Ftl1 to investigate this hypothesis, especially since earlier studies showed that *Leishmania* infection raises the intracellular iron pool of host cells by simultaneously enhancing iron import and decreasing iron export ^78, 94^. Our results confirmed the increase of Ftl1 levels during infection, unlike Fth1, and demonstrate its active sequestration by living parasites (Figure 9). This dual strategy may enable the parasite to circumvent iron toxicity while securing a controlled supply of iron, especially since *Leishmania* lacks a ferritin homolog ^75^. Ftl1 is not transcriptionally regulated during macrophages infection with *L. major* and *L. amazonensis* and *L. donovani* ^8, 28^. Consequently, its elevated abundance most likely reflects a reduction in protein degradation rather than an increase in protein synthesis. This interpretation is reinforced by our proteomic data: despite Ftl1 being delivered to lysosomes (PV) for degradation through the Ncoa4-dependent pathway ^71^, quantitative proteomics reveal a concerted down-regulation of lysosomal hydrolases. The reduced activity of these enzymes would attenuate lysosomal breakdown of Ftl1, thereby accounting for its accumulation during infection. Moreover, trafficking of Ftl1 to the PV by Ncoa4 is uncoupled from the canonical host ferritinophagy machinery, indicating that the parasite hijacks Ncoa4-mediated cargo delivery while bypassing host-cell degradation pathways. Our data further suggest that *Leishmania* blocks Ncoa4 iron-dependent turnover. The up-regulation of both Ftl1 and Ncoa4 correlates with infection status rather than with intracellular iron concentration. Under normal physiological conditions, elevated iron induces ferritin synthesis and triggers Ncoa4 degradation via autophagy and Herc2-mediated proteasomal pathways, while low-iron conditions stabilize and allow Ncoa4 to promote ferritin degradation and iron release ^70^. The simultaneous increase of ferritin and Ncoa4 during *Leishmania* infection deviates from their typical inverse relationship, likely reflecting a parasite-driven strategy to recruit ferritin without triggering its subsequent degradation. This is not the only direct impact of *L. donovani* on host iron homeostasis.

**FIGURE 9:**
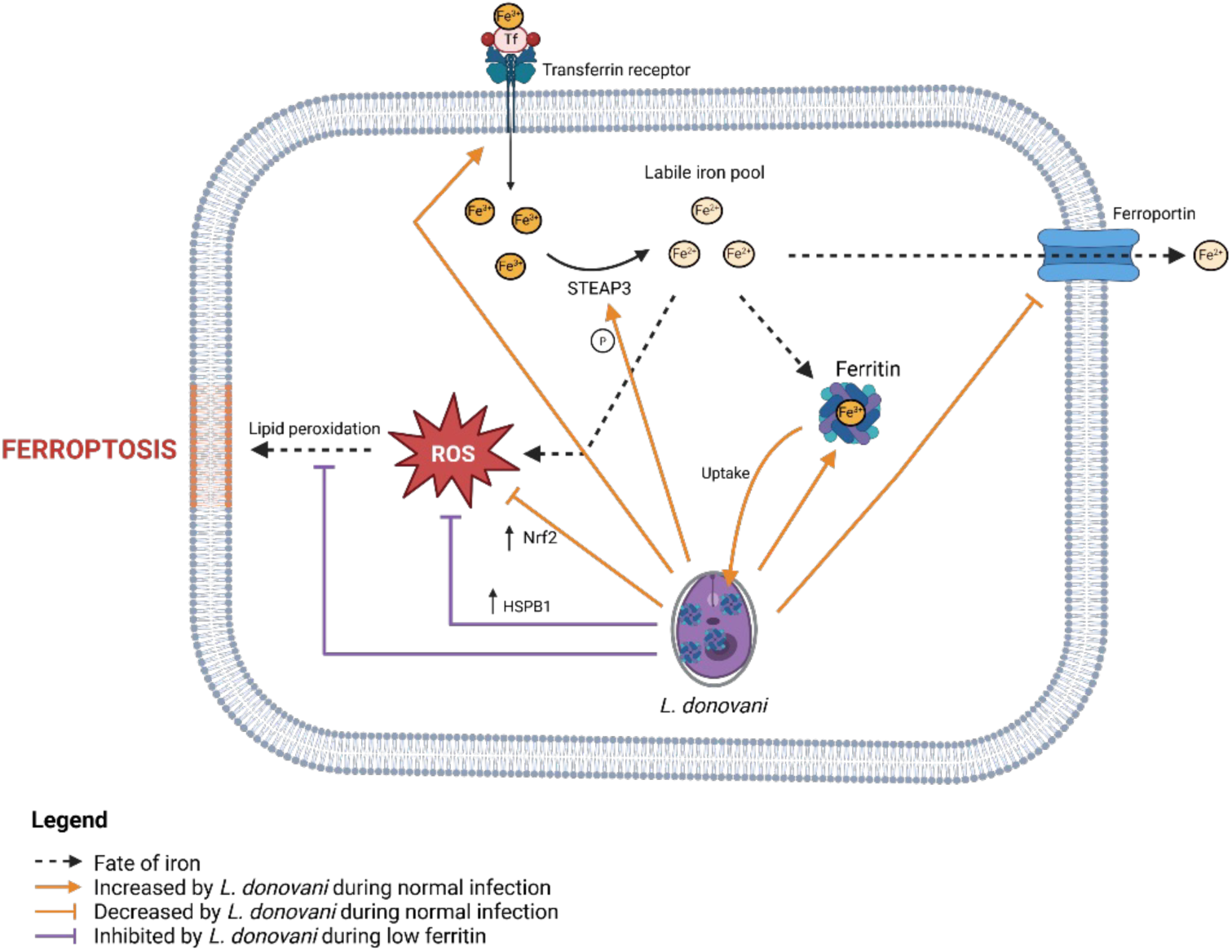
Schematic showing the different potential mechanisms used by *L. donovani* to modulate the host iron homeostasis. It combines previously published data with the new results presented here. The orange arrows present the modulation that occur during normal *Leishmania* infection while the purple arrows described the alteration observed when host Ftl1 is knocked down.

Our findings also reveal that *L. donovani* might prime infected macrophages for iron-rich environments, such as liver and spleen, by skewing ferritin composition toward the light-chain-rich form. This specific composition is characteristic of tissues with high iron turnover and storage capacity, and higher Ftl1/Fth1 ratio is known to enhance iron loading efficiency and structural stability of the ferritin complex ^63, 95^. We presented evidence suggesting that the shift toward an Ftl1-rich ferritin pool occurs independently of any exogenous iron cue. Remarkably, the rise in Ftl1 protein abundance is correlated to an increase in Ftl1 phosphorylation at Ser-169 in *Leishmania donovani*-infected macrophages. By contrast, phosphorylation of Fth1 at Ser-5 remains unchanged during the first 24 h of infection, similar to the pattern observed in uninfected cells, and is undetectable at 48 h, consistent with an absence of Fth1 up-regulation. These findings that require confirmation suggests that post-translational modification may differentially affect the stability or turnover of the ferritin subunits, thereby enabling the parasite to remodel the host ferritin composition in its favour. Notably, this shift to Ftl1-rich ferritin appears to be driven solely by the presence of specific *Leishmania* species, as it is observed during *L. donovani* but not *L. amazonensis* infection, highlighting a niche-specific strategies employed by the parasite to manipulate its host. This finding, as well as highlighting once again that different *Leishmania* species elicit distinct modulatory effects on macrophages ^1^, it provides a possible mechanism for the differential tissue tropism observed among *Leishmania* species. Indeed, species capable of visceralization may pre-adapt their macrophage host to environments encounter in internal organs (eg. high iron concentration), whereas cutaneous species may remain restricted to the skin due to their inability to modulate the macrophage in a similar manner. This diversity in host manipulation might also explain why drugs show variable efficacies across *Leishmania* species.

Our study provides clear evidence that *Leishmania* actively manipulates host cellular processes by recruiting ferritin in its vicinity, but we also demonstrated that the host, in turn, might have an impact on the parasite, as host ferritin is internalised by *Leishmania*, likely to serve as an iron reservoir. This finding, which is consistent with the parasite lacking ferritin orthologs, suggests a bidirectional interplay between host and pathogen during infection, a dimension that has been largely overlooked to date. De Souza *et al.* demonstrated that *Leishmania* amastigotes internalise class II MHC molecules and subsequently degrade them, thereby (i) acquiring nutrients and (ii) compromising antigen-presentation pathways. In contrast, our data demonstrate that the internalised ferritin is observed in the parasite cytoplasm as well as in the nucleus, an observation that argues against ferritin being only internalised as a nutrient. Moreover, ferritin persists within the parasite over time. The parasite pool of ferritin might serve as iron storage and to protect the parasite DNA from ROS-induced damage, as previously shown in mammalian cells ^51^. To our knowledge, this is the first report of a host protein being internalized by *Leishmania* without subsequent degradation, suggesting a novel strategy in which the parasite uses ferritin for iron storage rather than for direct nutrient acquisition. These findings raise several important questions, most notably whether the parasite exploits ferritin to store its own cytoplasmic and nuclear iron. Furthermore, this discovery opens new avenues for investigating whether other host proteins are similarly internalised during infection, thereby adding a novel dimension to our understanding of host-pathogen interactions.

Silencing Ftl1 did not impair intracellular *Leishmania* likely for two reasons: (1) The knockdown of Ftl1 was incomplete, achieving only partial suppression of the protein, and (2) the parasite compensates by up-regulating Hspb1, a small heat-shock protein that protects cells from ferroptosis ^96^. The Hspb1-mediated response is absent in uninfected macrophages and in a typical infection, even though Hspb1 is well known to suppress ROS production by inhibiting the Fenton reaction ^79^. Because Hspb1 also down-regulates surface transferrin-receptor-1 (TfR1), thereby limiting iron uptake ^80^, its activation would be detrimental to the parasite. Consequently, *Leishmania* is directly or indirectly regulating the expression of Hspb1 as a “last-resort” strategy when ferritin-based iron storage is unavailable. In addition to up-regulating Hspb1, *Leishmania* employs multiple, overlapping mechanisms to control lipid peroxidation and iron homeostasis, thereby securing enough iron while avoiding ferroptosis cell death. This redundancy explains why silencing Ftl1 does not affect parasite survival: alternative pathways can compensate when ferritin-mediated iron storage is unavailable. Such multilayered strategies likely arise from the parasite continual co-evolution with its host and the preservation of any effective adaptation ^1^. Nevertheless, each compensatory mechanism has a physiological cost, suggesting that *Leishmania* might prioritize its responses in a hierarchical fashion to preserve an optimal risk-benefit balance.

Control of iron homeostasis is not unique to *Leishmania* parasites. Many pathogens, including bacteria and fungi, also depend on iron for survival and proliferation within host cells. In response to infection, host cells actively restrict pathogen access to iron, a strategy often referred to as nutritional immunity^97^. To limit iron availability, host cells can increase iron export, reduce iron import, and enhance iron sequestration in storage proteins such as ferritin, thereby creating an iron-depleted environment that impairs pathogen growth ^98^. Two main non-haem strategies have been described for salvaging iron from the host, mainly in bacteria. One is based on siderophore production: *Mycobacterium tuberculosis* synthesizes high affinity siderophores, such as mycobactin and carboxymycobactin, that extract iron from host proteins like transferrin and lactoferrin, as well as transport it via the IrtAB system into the bacteria ^99, 100^; *Neisseria meningitidis* can directly or indirectly induce ferritin degradation to release iron that is then captured by its siderophores; or *Bacillus cereus* destabilises ferritin subunits to promote iron release, which is subsequently bound by its bacillibactin siderophore. The other strategy relies on surface- or membrane-anchored proteins that target host iron-containing proteins such as ferritin; for instance, *Listeria monocytogenes* uses a surface-associated ferric reductase system to access iron from host ferritin ^101^. Pathogenic fungi appear to use strategies similar to those of bacteria. For instance, *Candida albicans* hyphae express the cell-surface protein Als3, which acts as a receptor for host ferritin. Once ferritin is bound, local acidification promotes iron release, and the freed iron is then taken up by the fungus via its reductive iron uptake pathway ^102^. These strategies differ from those employed by *Leishmania*. Whereas bacteria and fungi typically degrade ferritin or extract iron directly from it, *Leishmania* instead exploits host ferritin as an iron reservoir, effectively using ferritin itself as a storage unit rather than simply a source to be dismantled.

The main question is therefore how *Leishmania* obtains iron. Several mechanisms have been described in prokaryotes and fungi that may provide clues. Ferritin is known to be unstable at low pH; acidification promotes both ferritin destabilisation and the reduction of Fe³⁺ to Fe²⁺, which is more soluble and can diffuse out of the ferritin shell ^103^. Because *Leishmania* resides in the phagolysosome, where the pH is around 5.5, this acidic environment could facilitate iron release from host ferritin and thus represent one route by which the parasite acquires iron once inside the vacuole. Similar to bacteria and fungi, *Leishmania* also expresses a surface ferric reductase (LFR) that converts Fe³⁺ to Fe²⁺, coupled to a ferrous iron transporter (LIT) that mediates Fe²⁺ uptake ^104^. The conversion of ferric to ferrous iron is thought to be regulated so that excess Fe²⁺ does not enter the parasite and cause toxicity. Our work reveals an additional, parasite-specific strategy that appears to compensate for the absence of siderophores or intrinsic ferritin-like storage proteins, by exploiting host ferritin instead. In this model, the parasite uses host ferritin as a safe iron reservoir in its immediate vicinity and within itself, avoiding iron-mediated toxicity. Together, our data and published evidence suggest that *Leishmania* might combine two broad strategies: one resembling those used by bacteria and fungi, and a second, distinct mechanism that we describe here for the first time, which may be specific to parasites that lack their own iron storage systems.

## Material and methods

### Key resource table

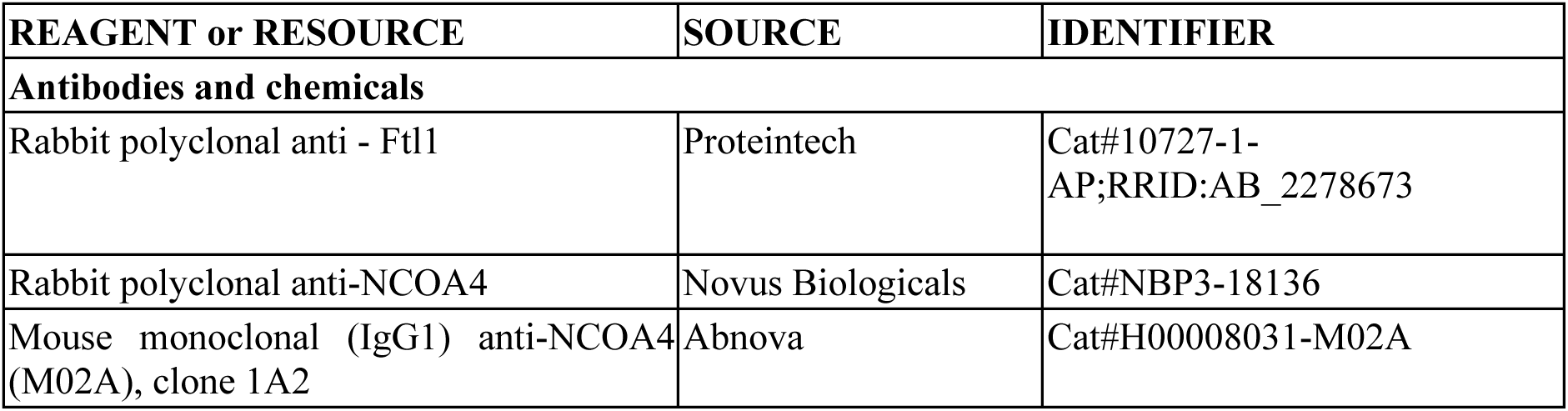
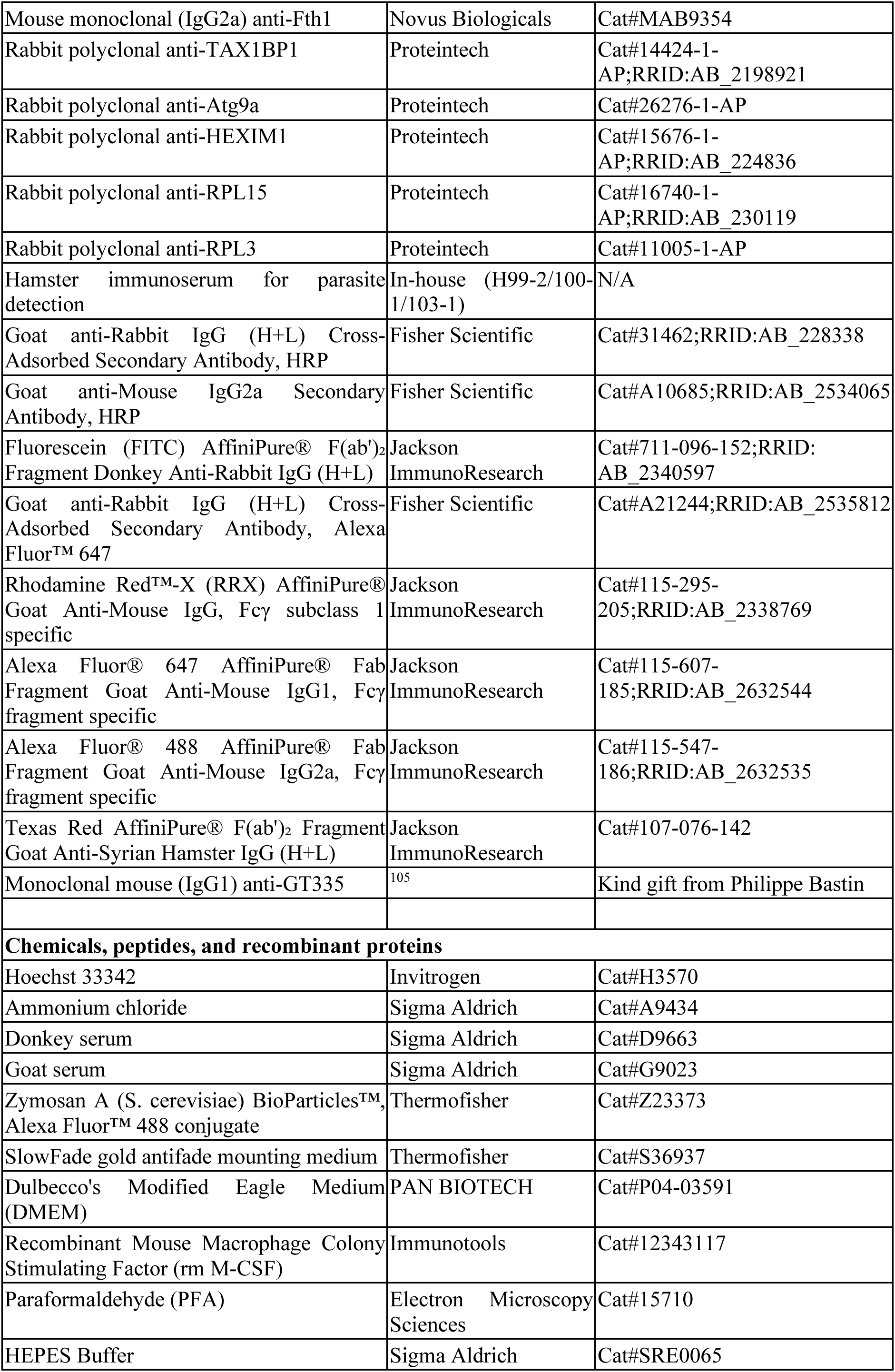
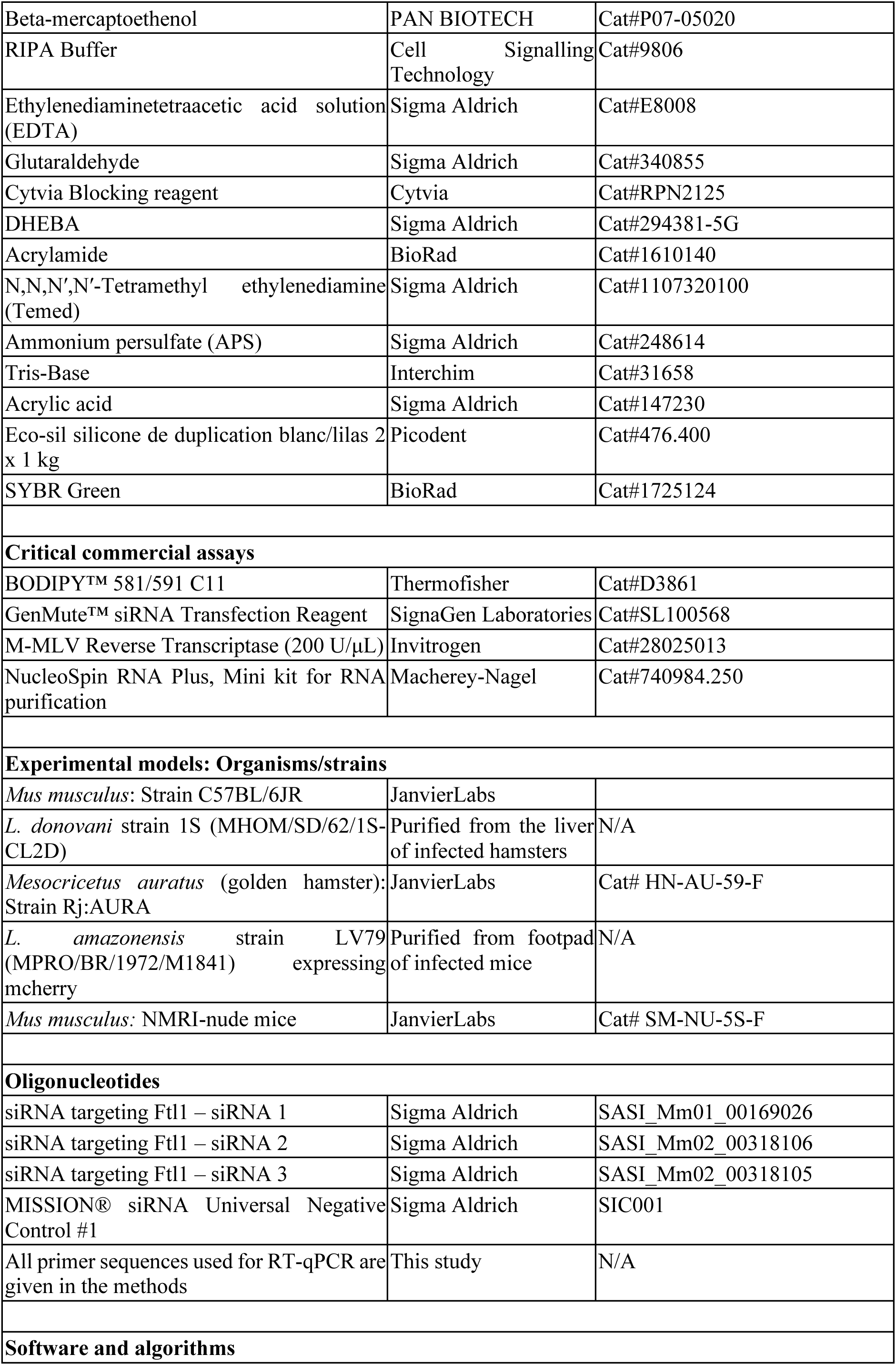
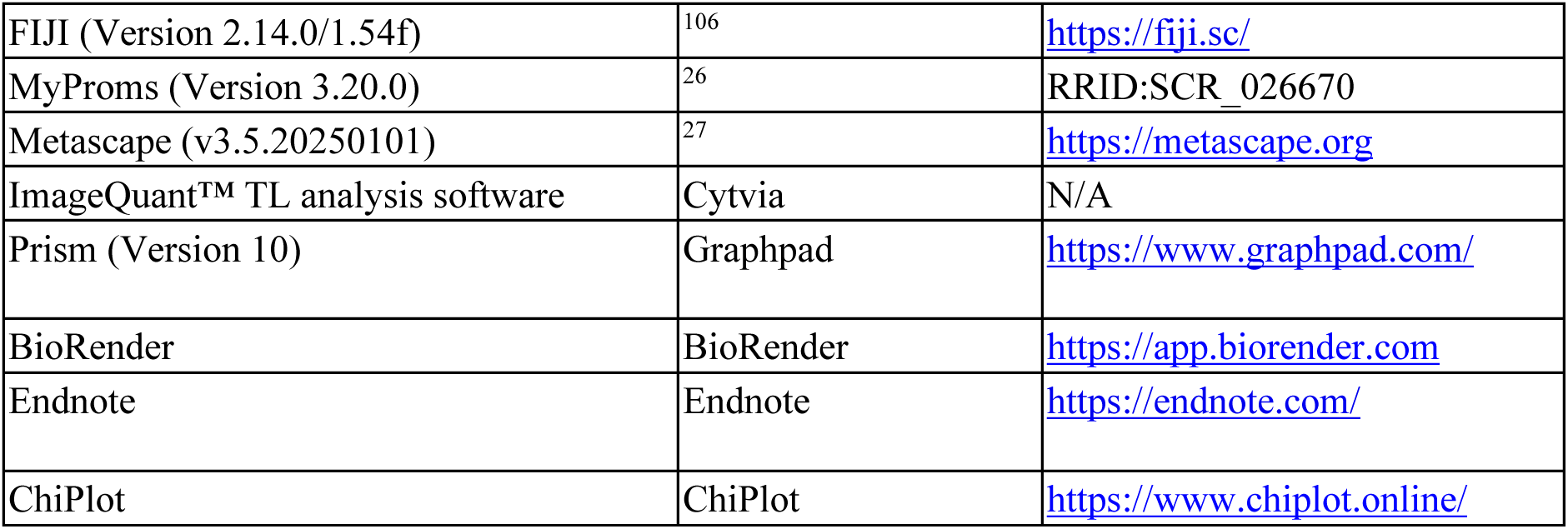

### Ethics statement

Work on animals was performed in compliance with French and European regulations on care and protection of laboratory animals (EC Directive 2010/63, French Law 2013-118, February 6th, 2013). All animal experiments were approved by the Ethics Committee and the Animal welfare body of Institut Pasteur and by the Ministère de l’Enseignement Supérieur, de la Recherche et de l’Innovation (project n°#19683).

### Animals

Four to ten weeks old female Syrian golden hamsters (*Mesocricetus auratus*), C57BL/6 mice (*Mus musculus*) and NMRI-nu mice *(Mus musculus)* were purchased from Janvier Labs and all animals were hosted in A3 animal facility at the Institut Pasteur. All animals were handled under specific, pathogen-free conditions in biohazard level 3 animal facilities (A3) accredited by the French Ministry of Agriculture for performing experiments on live rodents (agreement A75-15-01).

### Production of mouse bone marrow derived macrophages (BMDMs)

Bone marrow cell suspensions were recovered from tibias and femurs of C57-BL/6 mice in DMEM medium (Pan Biotech, #P04-03591) and cultured in medium complemented with Recombinant Mouse Macrophage Colony Stimulating Factor (rm M-CSF, Immunotools #12343117). Precursors were seeded for one night at 37°C in a 7.5% CO2 atmosphere at 3 × 10^7^ cells / 12 ml of complete medium containing 50 ng/ml of rm M-CSF-1, in Petri dishes for tissue culture (Falcon® 100 mm TC-treated Cell Culture Dish ref OPTILUX 353003) for the removal of unwanted adherent cells. Unattached cells were recovered and cultured at one million cells per ml in non-treated Petri dishes (Greiner bio-one 664161) at 37°C in a 7.5% CO2 atmosphere for 6 days with 50 ng/mL rm M-CSF-1.

### Isolation of *L. donovani* amastigotes and BMDM infection

For *L. donovani*, anesthetized hamsters were inoculated by intra-cardiac injection of 5×10^7^ *L. donovani* strain 1S (MHOM/SD/62/1S-CL2D) amastigotes purified from hamster infected spleens as described previously ^107^. The weight of the animals was recorded over time, and the animals were euthanized by CO_2_ asphyxiation before they reached the end-point of infection represented by a 20% loss of body weight. Amastigotes were then purified from the infected hamster liver and spleens with the previously published protocol ^107, 108^.

mBMDMs were seeded in either 6 well or 24 well plates according to the specific experiments in DMEM media containing 30 ng/mL rm M-CSF. *L. donovani* amastigotes, purified from the liver and spleen of infected hamsters, were used to infect BMDMs at the specific multiplicity of infection (MOI). The cells were incubated at 37°C in a 7.5% CO2 atmosphere for the appropriate durations.

### Isolation of *L. amazonensis* amastigotes and BMDM infection

The mCherry expressing *L. amazonensis* strain LV79 (MPRO/BR/1972/M1841) was propagated in NMRI-nude mice and amastigotes were purified from cutaneous lesions on the footpad of infected mice. Mouse BMDMs were plated in 6 well plates with 2.5 million cells per well. Twenty-four hours after plating, the cells were left uninfected or infected at an MOI of 2 with *L. amazonensis* amastigotes purified from footpad of infected NMRI-nude mice. After 48 hours of infection, whole cell lysates were prepared using RIPA lysis buffer (1XRIPA buffer, 1mM PMSF, Benzonase, cOmplete EDTA-free protease inhibitor cocktail). The samples were then sonicated for 5 min with 30sec ON/30sec OFF setting and centrifuged at 14000g for 15 min at 4°C. The supernatant was collected and quantified using Bradford quantification. The samples were then used for performing western blot analysis.

### Zymosan A treatment

Mouse BMDMs were seeded in 24 well plates on coverslips at a concentration of 1.5*10^5^ cells per well or 6 well plates at 2.5 million cells per well. Zymosan A particles (Thermofisher) were diluted in 1XPBS, counted and concentration of 10 beads per cell was prepared in DMEM and added to cells. The cells were incubated with Zymosan A particles for 4 hours at 37°C in a 7.5% CO2 atmosphere. Following incubation, for 6 well plates, the cells were washed twice with 1XPBS and lysed for mass spectrometry analysis. For 24 well plates, the cells were fixed with 4% paraformaldehyde (PFA).

### Sample preparation and analysis for label-free, quantitative phosphoproteomic and proteomic analyses

Mouse BMDMs were seeded in 6 well plates at the concentration of 2.5 million cells per well. Two wells were seeded for each condition. 24 hours after seeding, the cells were either left uninfected or infected with *L. donovani* amastigotes at the MOI of 10. For zymosan treatment, the cells were treated with Zymosan A particles (Thermofisher) for four hours. After the appropriate timepoints of infection/Zymosan A treatment, cells were washed twice with 1XPBS. Cell lysates were prepared by adding the lysis buffer (8M Urea/ 200mM ammonium bicarbonate) and scraping the cells to ensure homogenous lysis. Following sonication for 5 min with 30 sec ON/30 sec OFF settings, the cell lysates were centrifuged at 14,000g for 15 min at 4°C. The supernatant was collected and stored at -80°C until further use.

For mass spectrometry analysis, 125 µg of each cell lysate were reduced with 5 mM DTT for 1 h at 37°C and alkylated with 10 mM iodoacetamide for 30 min at room temperature in the dark. The samples were then diluted in 100 mM ammonium bicarbonate to reach a final concentration of 1 M urea and digested with Trypsin/Lys-C (Promega CAT#: V5071) at 37 °C in an enzyme/protein ratio of 1:50. After overnight digestion, enzymatic activity was quenched by acidifying the lysates using formic acid (FA) at a final concentration of 5%. Precipitates were removed by centrifugation at 4000 rpm and then loaded onto homemade SepPak C18 Tips packed by stacking ten AttractSPE disk (#SPE-Disks-Bio-C18-100.47.20 Affinisep) into a 200µL micropipette tip for desalting. Peptide were eluted and 90% of the starting material was enriched using Titansphere^TM^ Phos-TiO kit centrifuge columns (#5010-21312, GL Sciences) as described by the manufacturer. After elution from the Spin tips, the phospho-peptides and the remaining 10% eluted peptides were vacuum concentrated to dryness and reconstituted in 0.1% FA prior to liquid chromatography-tandem mass spectrometry (LC-MS/MS) phosphoproteome and proteome analysis.

Phosphoproteome analyses were performed with a RSLCnano system (Ultimate 3000, Thermo Scientific) coupled online to an Exploris 480 mass spectrometer (Thermo Scientific). Peptides were trapped on a 2 cm nanoviper Precolumn (i.d. 75 μm, C18 Acclaim PepMap^TM^ 100, Thermo Scientific) at a flow rate of 2.5 µL/min in buffer A (2/98 CH_3_CN/H_2_O in 0.1% FA) for 4 min to desalt and concentrate the samples. Separation was performed on a 50 cm x 75 μm C18 column (nanoViper Acclaim PepMap^TM^ RSLC, 2 μm, 100Å, Thermo Scientific) regulated to a temperature of 50°C with a linear gradient of 2% to 30% buffer B (100% CH_3_CN in 0.1% FA) or 2% to 25% buffer B (for the 48h PI samples) at a flow rate of 300 nL/min over 91 min. For proteome analyses, an Exploris 480 or an Orbitrap Eclipse mass spectrometer (Thermo Scientific) were used, both coupled to a RSLCnano system as described previously, with some minor modifications. Peptide trapping was at 3.0 µL/min for the 48h PI samples and separation used a linear gradient of 2% to 35% buffer B or 2% to 30% (for the 48 PI samples) over 211 min.

Exploris MS full scans were performed in the ultrahigh-field Orbitrap mass analyzer in ranges m/z 375–1500 with a resolution of 120 000 (at m/z 200). The top 20 most intense ions were subjected to Orbitrap for further fragmentation via high energy collision dissociation (HCD) activation and a resolution of 15 000 with the normalized AGC target set at 100%. We selected ions with charge state from 2+ to 6+ for screening. Normalized collision energy (NCE) was set at 30 and the dynamic exclusion to 40s.

Orbitrap Eclipse MS1 data were collected in the Orbitrap (120,000 resolution; maximum injection time 60 ms; AGC 4 x 10^5^). Charges states between 2 and 5 were required for MS2 analysis, and a 45 s dynamic exclusion window was used. MS2 scan were performed in the ion trap in rapid mode with HCD fragmentation (isolation window 1.2 Da; NCE 30%; maximum injection time 60 ms; AGC 10^4^)

### Phosphoproteomic and proteomic data analysis

For identification, the data were searched against the *Mus Musculus* (UP000000589) UniProt database by adding or not (for the 48h PI phosphoproteome samples) the L-CK1.2 sequence, and the Ld1S database for the 48h PI proteome samples using Sequest HT through Proteome Discoverer (version 2.4). Enzyme specificity was set to trypsin and a maximum of two missed cleavage sites were allowed. Oxidized methionine, N-terminal acetylation, methionine loss and methionine acetylation loss were set as variable modifications. Phospho serine, threonine and tyrosines were also set as variable modifications in phosphoproteome analyses. Maximum allowed mass deviation was set to 10 ppm for monoisotopic precursor ions and 0.02 Da for MS/MS peaks from the Orbitrap Exploris 480 instrument and 0.6 Da for MS/MS peaks from the Orbitrap Eclipse instrument. The resulting files were further processed using myProMS [^26^; https://github.com/bioinfo-pf-curie/myproms] v.3.10. False-discovery rate (FDR) was calculated using Percolator ^109^ and was set to 1% at the peptide level for the whole study. Label-free quantification was performed using peptide extracted ion chromatograms (XICs), computed with MassChroQ ^110^ v.2.2.1. For protein quantification, XICs from proteotypic peptides shared between compared conditions (TopN matching for proteome setting and simple ratios for phosphoproteome) with missed cleavages were used. Median and scale normalization at peptide level was applied on the total signal to correct the XICs for each biological replicate (N=5). The phosphosite localization accuracy was estimated by using the PtmRS node in PD, in PhosphoRS mode only. Phosphosites with a localization site probability greater than 75% were quantified at the peptide level. To estimate the significance of the change in protein abundance, a linear model (adjusted on peptides and biological replicates) was performed, and p-values were adjusted using the Benjamini–Hochberg FDR procedure. The mass spectrometry proteomics raw data have been deposited to the ProteomeXchange Consortium via the PRIDE ^111^ partner repository with the dataset identifier PXD070745.

### Kinase enrichment analysis

The kinase enrichment was performed using the method proposed for serine and threonine kinases by Johnson *et al.* ^29^ and later extended by Yaron-Barir *et al.* for tyrosine kinases ^112^, using the implementation provided by the KINference R tool ^113^. For each enrichment of a phosphoproteomic experiment P_exp_, we used the set of all phosphorylation sites P = [p ∈ P_exp_ | (|log2-Ratio(p)| ≥ 1 ∧ adj_pval_p_ ≤ 0.05) ∨ (|log2-Ratio(p)| < 1)] as input, where the absolute log2-Ratio was greater than 1.0 and the adjusted p-value was lower than 0.05 or the absolute log2-Ratio was lower than 1.0. The method of Johnson *et al.* and Yaron-Barir *et al.* uses curated 303 serine/threonine and 93 tyrosine kinase motifs to score the likelihood of a kinase targeting a phosphorylation site p ∈ P. Afterwards, for each kinase, the likely targeted phosphorylation sites of the respective kinase are split into a background, downregulated and upregulated set based on the log2-Ratio. The upregulated set consists of all phosphorylation sites p with log2-Ratio(p) ≥ 1 and the downregulated set of all phosphorylation sites p with log2-Ratio(p) ≤ -1. The background set consists of all phosphorylation sites p with -1 ≤ log2-Ratio(p) ≤ 1. Then, the activity of each kinase is determined utilizing a one-sided Fisher’s exact test to compute the significance of the up- or downregulated set compared to the background set respectively. For further details, see the Methods section of Johnson *et al.* ^29^.

### Immunofluorescence assay

In 24 well plates, coverslips were added and cells were seeded at 1.5 X 10^5^ cells per well. These cells were either infected or not with *L. donovani* amastigotes. Following infection, appropriate treatments were done and the cells were washed using 1X PBS and then fixed at the appropriate time points using 4% paraformaldehyde. These fixed coverslips were then used to perform immunofluorescence assays. The cells were first blocked with 50mM NH_4_Cl (in 1XPBS) for 20 minutes. Following this they were incubated with a solution of 0.1%TritonX100 and 10% Donkey or Goat serum for 15 minutes. The coverslips were then incubated with the optimised concentrations for the appropriate primary antibodies for two hours at room temperature in dark. After washing the coverslips with 1XPBS to remove any remaining primary antibody, secondary antibody incubation was performed for one hour at room temperature in dark. The secondary antibody was then washed off and the cells were incubated in Hoechst (1:1000) for 10-15 minutes at room temperature in dark. The Hoechst was then first washed with 1XPBS and then 2 times with distilled water and mounted on SuperFrost microscopy slides using SlowFade gold antifade mounting medium.

### Counting infection

Infected cells in 24 well plates were fixed with 4% PFA at the timepoint required for counting the infection. Immunofluorescence assay was performed to stain the parasites by using the antibody Hamster immunoserum for parasite detection (1:400) followed by secondary antibody staining using Texas Red conjugated anti-Syrian Hamster IgG (5µg/mL). The cells were then stained with Hoechst 33342 and mounted on slides. Images were taken in Carl Zeiss Apotome inverted microscopes (excitation lasers – 405nm and 594nm) using 20X oil objective. For parasite load, parasites per 100 cells were counted for three biological replicates and for around 600 cells per replicate. For percent infection, images of multiple fields were taken in Evos microscope (Thermo Fisher Scientific) with the 405nm excitation laser and 40X objective. Percentage of infected cells was calculated for each field and for 3 biological replicates.

### Western blot

Total cell protein lysates were prepared by first lysing the cells using RIPA lysis buffer (1XRIPA buffer, 1mM PMSF, Benzonase, cOmplete EDTA-free protease inhibitor cocktail). The samples were then sonicated for 5 min with 30sec ON/30sec OFF setting and centrifuged at 14000g for 15 min at 4°C. The supernatant was collected and quantified using Bradford quantification. The samples were then resolved via SDS PAGE gel in MES running buffer and transferred via wet transfer system. The proteins were probed by incubating with the appropriate primary antibodies at 4°C overnight followed by the secondary antibodies for 1 hour at RT. The membranes were revealed via chemiluminescence using Cytvia’s Amersham ECL Western Blotting Detection Reagent.

### Quantification of Ftl1 localisation around parasites

BMDMs were plated in 24 well plates at 0.15 million cells per well and infected or not with *L. donovani* amastigotes at MOI 10. The cells were fixed at four hours, 48 hours and 6 days post infection. Immunofluorescence staining was performed for all the three timepoints by staining with the primary antibodies Ftl1 (Proteintech, 1:400), Hamster immunoserum for parasite detection (1:400). This was followed by secondary antibodies FITC-anti Rabbit and Texas Red-anti Hamster at 5µg/mL and Hoechst staining to stain the nuclei. For quantifying Ftl1 around parasites, the cells were imaged with Carl Zeiss Apotome inverted microscope (excitation lasers – 405nm, 594nm) using 63X oil objective. At least 100 cells per replicate, for total of three biological replicates, were counted to assess percentage of parasites per cell having Ftl1 around.

### Quantification of Fth1 localisation around parasites

BMDMs were plated in 24 well plates at 0.15 million cells per well and infected or not with *L. donovani* amastigotes at MOI 10. The cells were fixed at 48 hours post infection. Immunofluorescence staining was performed for Fth1 (Novus, 1:50) and Hamster immunoserum for parasite detection (1:400). This was followed by secondary antibodies FITC-anti Rabbit and Texas Red-anti Hamster at 5µg/mL and Hoechst staining to stain the nuclei. For quantifying Ftl1 around parasites, the cells were imaged with Leica SP8 at 63X magnification (excitation lasers – 405nm, 594nm) using 63X oil objective. Atleast 100 cells were counted per replicate (from three biological replicates in total) to assess percentage of parasites per cell having Fth1 around them.

### Sample preparation and immunolabelling by Tokuyasu Technique for transmission electron microscopy

Mouse bone marrow derived macrophages were infected or not by *L. donovani* amastigotes fixed with 0.1% glutaraldehyde (Sigma G5882) and 2% paraformaldehyde (EMS 15714) in DMEM medium 10 min at room temperature, then 2h at room temperature with 0.1% glutaraldehyde (Sigma G5882) and 2% paraformaldehyde (EMS 15714) in DPBS1X (STEMCELL 37354) and overnight at 4°C. The fixative is replaced by a solution of 1% paraformaldehyde in DPBS1X and store at 4°C until use. After three wash in DPBS1X sample are incubate 15 min. in Ammonium Chloride 50mM at room temperature (RT) and resuspended in PBS with 10% gelatin (Sigma G2500), incubate 10 min at 37°C. The samples were allowed to solidify on ice and infiltrated with 2.3M sucrose at 4°C overnight. Following that, they were mounted onto sample pins and frozen in liquid nitrogen. Subsequently, the samples were cryo-sectioned (60nM thickness) using FC6/UC6- cryo-ultramicrotome (Leica) and a 35° diamond knife (Diatome). The sections were picked up using a 1:1 mixture of 2% methyl cellulose (Sigma Aldrich M-6385) and 2.3M sucrose. Samples were quenched with 50 mM NH_4_Cl, blocked in PBS with 1% BSA and immunolabeled either with anti-Ferritin light chain Polyclonal antibody in PBS with 1% BSA (d=1/50) and protein A-gold (15 nm d= 1/50) treatment for 15 min or with anti-Ferritin Heavy Chain Antibody in PBS with 1% BSA (d=1/250) and protein A-gold (15 nm d= 1/10) treatment for 15 min. Finally, the samples were fixed with 1 % Glutaraldehyde in DPBS1X for 5 min. The sections were thawed and stained/embedded in 4% uranylacetate/2% methyl cellulose mixture (1:9). Images were recorded with TECHNAISPIRIT 120 Kv (with bottom-mounted RIO16 Camera 4Kx4K).

### Colocalization analysis of Ftl1, TAX1BP1 and Atg9 with NCOA4 using confocal microscopy

BMDMs were plated in 24 well plates at 0.15 million cells per well and infected or not with *L. donovani* amastigotes at MOI 10. The cells were fixed at four hours, 48 hours and 6 days post infection. Immunofluorescence staining was performed for all the three timepoints with the antibodies Hamster immunoserum for parasite detection (1:400) and NCOA4 (Abnova, 1:50) along with either Ftl1 (Proteintech, 1:400), TAX1BP1 (Proteintech, 1:50) or Atg9A (Proteintech, 1:50). This was then followed by secondary antibodies FITC-anti Rabbit, Texas Red-anti Hamster and Alexa Fluor® 647-anti Mouse IgG1 all at 5µg/mL and Hoechst staining to stain the nuclei. For colocalisation analysis, between Ftl1/TAX1BP1/Atg9A and NCOA4, the immunostained cells were imaged using Leica SP8 at 63X magnification with the excitation lasers 405nm, 594nm and 647nm and optimal XY resolution. Deconvolution and colocalization analysis were done on the images using Huygens Professionals software. The images were first deconvolved using the pipeline ‘Deconvolution Wizard’. Threshold was calculated by the auto mode and all other settings were kept default. Post deconvolution, colocalization analysis was performed using the pipeline ‘Colocalisation Wizard’. Thresholding was done by Costes method after which Pearson’s coefficient was calculated.

### Gene knock-down by small RNA interfering (siRNA)

Knockdown of *Ftl1* transcripts was performed using the predesigned Ftl1 siRNA (Sigma). BMDMs were seeded in 6 well plates at 3X10^6^ cells per well or in 24 wells with coverslips at 1.5 X 10^5^ cells per well and incubated overnight. Cells were then transfected using the GenMute™ siRNA Transfection reagent (SignaGen Laboratories, SL100568-PMG) with 70nM of Ftl1 siRNA or Universal negative control siRNA (SIC001) and incubated at 37°C. After 4 hours of incubation, the reagent was washed off using prewarmed 1XPBS and fresh media was added. The BMDMs were then infected or not with purified *L. donovani* amastigotes at MOI 10. After 48 hours infection, total protein and RNA was extracted, and the knockdown was confirmed using qPCR and western blot methods.

### RT-qPCR analysis

Total RNA isolation was performed using NucleoSpin® RNA plus kit (Macherey-Nagel) following the manufacturer’s instructions followed by quality control using Nanodrop and reverse transcription to obtain cDNA. The list of primers used for the RT-qPCR analyses is provided below:

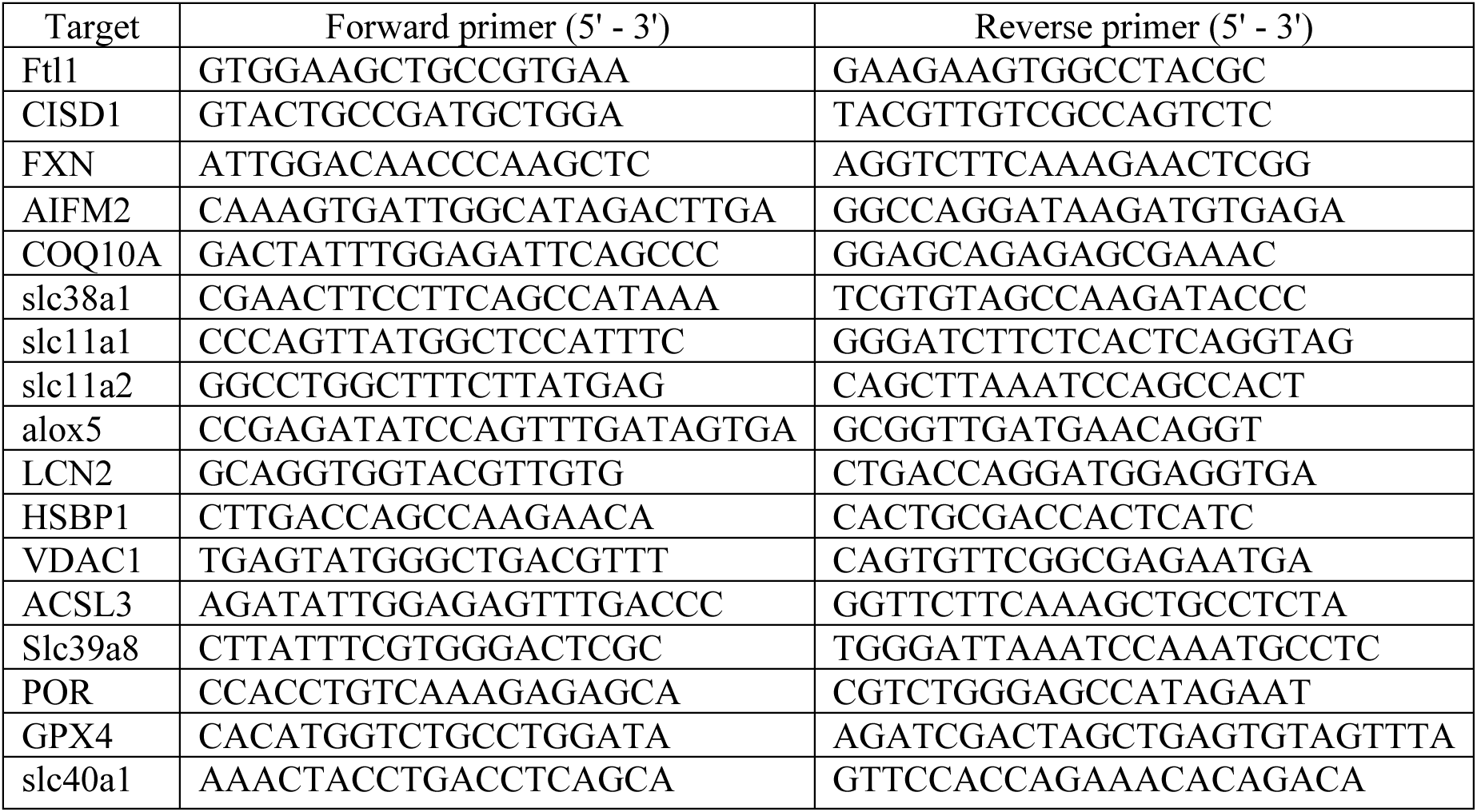

### BODIPY C11 staining and quantification

Ftl1 knockdown using si-Ftl1 was performed in uninfected and *L. donovani* infected macrophages according to the protocol given. 48 hours after siRNA treatment, the cells were washed twice with 1XPBS and then incubated for 15 min at RT with 4µM BODIPY™ 581/591 C11 + 10µg/mL Hoechst 33342 in 1XPBS. The solution was washed twice with 1XPBS and DMEM (without FCS) was added to the cells. The cells were imaged by Evos microscope (Thermo Fisher Scientific) with the 405nm, 488nm and 594nm excitation lasers and 20X objective.

To analysis the level of lipid peroxidation, the cells were first segmented using the machine learning software Ilastik. The software was trained for cell segmentation using 15 images from the red channel (594nm excitation). Following this, all the images were provided to the trained algorithm and the segmentation outputs were exported. These outputs were used in Fiji to create masks and the intensity in the green channel (488nm excitation) and red channel (594nm excitation) was measured for each segmented cell and the ratio of green/red was calculated which was then used for the comparisons.

### Expansion microscopy

The protocol used for the expansion microscopy was adapted from Louvel *et al*. ^114^. Coverslips fixed with 4% paraformaldehyde (PFA) for 30 minutes at RT were incubated in 4% acrylamide (BioRad) + 4% PFA solution overnight at RT. A gelation chamber was prepared by first gluing two 22*22mm glass coverslips together and making two such sets. The two stacks of 22*22mm coverslips were glued on a classic glass microscope slide with a distance of 14 mm one between them. Then on each 22*22mm stacks, another 22*22mm coverslip was glued on top of it, leaving enough space to add a coverslip between the two stacks to cover the chamber which acts as a sliding door that can be freely moved. Finally, 2 more 22*22mm coverslips were glued on the stacked coverslips and on top of the door to act as a top cover, taking care that the door can still slide in and out without resistance.

For the gelation, the coverslip having the fixed cells was glued on the slide and in the space between the two 22*22mm stacks with the cells side facing the top. The sliding door was placed on top to close the gelation chamber. Then ∼60µL of 1MS solution (19% sodium acrylate, 10% acrylamide (BioRad), 0.1% DHEBA (Sigma) in 1XPBS) with 0.25% APS and 0.25% TEMED was added in gap between the coverslip having fixed cells and the door by sliding the door slightly so that it was at the edge of the coverslip. Following this, the gelation chamber was kept for 15 min in a humid chamber (a closed box lined with wet tissues) that had been precooled at -20°C for 30 min. After this, the humid chamber was transferred to 37°C for 45 min. The gelation chamber was then disassembled to remove the formed gel from the coverslip. The gel was then placed in a 1.5mL Eppendorf having 1mL of denaturation buffer (200mM SDS, 200mM NaCl and 50mM TrisBase in water with pH=6.8) and heated at 85°C for 1.5 hours. Following this, the gel was expanded by performing three washes, 30 min each, with distilled water. These gels were then used for immunofluorescence. The quarter of the expanded gel was cut and 2 washes, 10 min each, with 1XPBS were performed. The gel piece was placed in 12 well plate and primary antibody mix prepared in 2%BSA – Ftl1 (Proteintech) at 1:300, GT335 (Kind gift from Philippe Bastin, ^105^) at 1:5000 was added. The primary antibody incubation was performed for 3 hours at 37°C with shaking. Following this, 3 washes with 1XPBS+0.1%Tween, 10 min per wash, were done. The secondary antibody mix was prepared with the final concentrations of 5µg/mL in 2% BSA. This mix was added to the gels and incubated for 3 hours at 37°C with shaking. Following this, the secondary antibody was washed off using 3 washes with 1XPBS+0.1%Tween, 10 min per wash. The gel was then stained using Hoechst 33342 in 1XPBS for 15 min and 3 washes of 1XPBS, 10 min per wash, were done. The gel was then expanded with 3 washes of 30 min each in distilled water. A small piece of the gel was cut and placed on Ibidi µ-Dish 35 mm. After ensuring that the cell side is facing the bottom of the dish, the gels was fixed on the dish by adding 1:1 mix of Picodent Silicone (white and Lilac solutions) and letting it harden at RT for 5-6 hours. Following this, the sample was imaged in confocal settings.

### Statistical tools used

All the statistical analyses were performed with Graphpad Prism using unpaired t-test for p-value significance. Kinase enrichment analysis was performed with the KINference R tool.

## Supporting information

The supplementary figures are linked to the following text : Figure S1-S8 and Table S7

## Acknowledgments

The authors would like to thank Christine Girard-Blanc and Philippe Bastin, Institut Pasteur, for their help with expansion microscopy; the Unit of Technology and Service - Photonic BioImaging (UTechS PBI) from the Institut Pasteur and the Image Analysis Hub of the Institut Pasteur for the help with confocal microscopy and analyses of co-localisations, in particular Audrey Salles, Julien Fernandes, and Jean-Yves Tinevez. The authors would also like to thank Pascale Pescher and Eric Prina for their help with macrophage infections and Hervé Lecoeur for discussions and help with the Ferroptosis part of the project.

## Author contribution

NR conceived the study, SSS and NR designed the experiments. SSS, FD, NM, OG, CT performed the investigation and data acquisition. SSS and NR conducted the formal analysis and data interpretation. SSS and NR wrote the original draft. SSS, FD, NM, OG, CT, DBB, DL, AS and NR reviewed and edited the manuscript. NR supervised the research. AS, DBB and NR acquired funding. The final draft of the manuscript was approved by all co-authors.

## REFERENCES

1. Kaye P, Scott P. Leishmaniasis: complexity at the host-pathogen interface. Nature reviews Microbiology. 2011;9(8):604–15. doi: 10.1038/nrmicro2608. PubMed PMID: 21747391.

2. Bodhale N, Ohms M, Ferreira C, Mesquita I, Mukherjee A, Andre S, Sarkar A, Estaquier J, Laskay T, Saha B, Silvestre R. Cytokines and metabolic regulation: A framework of bidirectional influences affecting Leishmania infection. Cytokine. 2021;147:155267. doi: 10.1016/j.cyto.2020.155267. PubMed PMID: 32917471.

3. Fernandez-Garcia M, Mesquita I, Ferreira C, Araujo M, Saha B, Rey-Stolle MF, Garcia A, Silvestre R, Barbas C. Leishmania donovani Induces Multiple Dynamic Responses in the Metabolome Associated with Amastigote Differentiation and Maturation Inside the Human Macrophage. Journal of proteome research. 2023;22(7):2256–70. doi: 10.1021/acs.jproteome.2c00845. PubMed PMID: 37339249.

4. Majumder S, Dey R, Bhattacharjee S, Rub A, Gupta G, Bhattacharyya Majumdar S, Saha B, Majumdar S. Leishmania-induced biphasic ceramide generation in macrophages is crucial for uptake and survival of the parasite. J Infect Dis. 2012;205(10):1607–16. Epub 20120418. doi: 10.1093/infdis/jis229. PubMed PMID: 22517914.

5. Nandan D, Cherkasov A, Sabouti R, Yi T, Reiner NE. Molecular cloning, biochemical and structural analysis of elongation factor-1 alpha from Leishmania donovani: comparison with the mammalian homologue. Biochem Biophys Res Commun. 2003;302(4):646–52. PubMed PMID: Medline:12646217.

6. Silverman JM, Reiner NE. Leishmania exosomes deliver preemptive strikes to create an environment permissive for early infection. Frontiers in cellular and infection microbiology. 2011;1:26. Epub 20120109. doi: 10.3389/fcimb.2011.00026. PubMed PMID: 22919591; PMCID: PMC3417360.

7. Fernandes JCR, Zamboni DS. Mechanisms regulating host cell death during Leishmania infection. mBio. 2024;15(11):e0198023. doi: 10.1128/mbio.01980-23. PubMed PMID: 39392429; PMCID: 11559009.

8. Chaparro V, Graber TE, Alain T, Jaramillo M. Transcriptional profiling of macrophages reveals distinct parasite stage-driven signatures during early infection by Leishmania donovani. Sci Rep. 2022;12(1):6369. doi: 10.1038/s41598-022-10317-6. PubMed PMID: 35430587; PMCID: 9013368.

9. Ruvolo PP, Deng X, May WS. Phosphorylation of Bcl2 and regulation of apoptosis. Leukemia. 2001;15(4):515–22. doi: 10.1038/sj.leu.2402090. PubMed PMID: 11368354.

10. Singh AK, Pandey RK, Siqueira-Neto JL, Kwon YJ, Freitas-Junior LH, Shaha C, Madhubala R. Proteomic-based approach to gain insight into reprogramming of THP-1 cells exposed to Leishmania donovani over an early temporal window. Infect Immun. 2015;83(5):1853–68. Epub 20150217. doi: 10.1128/IAI.02833-14. PubMed PMID: 25690103; PMCID: PMC4399049.

11. Chakraborty S, Kumar A, Faheem MM, Katoch A, Kumar A, Jamwal VL, Nayak D, Golani A, Rasool RU, Ahmad SM, Jose J, Kumar R, Gandhi SG, Dinesh Kumar L, Goswami A. Vimentin activation in early apoptotic cancer cells errands survival pathways during DNA damage inducer CPT treatment in colon carcinoma model. Cell Death Dis. 2019;10(6):467. Epub 20190613. doi: 10.1038/s41419-019-1690-2. PubMed PMID: 31197132; PMCID: PMC6565729.

12. Marr AK, MacIsaac JL, Jiang R, Airo AM, Kobor MS, McMaster WR. Leishmania donovani infection causes distinct epigenetic DNA methylation changes in host macrophages. PLoS pathogens. 2014;10(10):e1004419. doi: 10.1371/journal.ppat.1004419. PubMed PMID: 25299267; PMCID: 4192605.

13. Kumari I, Lakhanpal D, Swargam S, Nath Jha A. Leishmaniasis: Omics Approaches to Understand its Biology from Molecule to Cell Level. Curr Protein Pept Sci. 2023;24(3):229–39. doi: 10.2174/1389203724666230210123147. PubMed PMID: 36809951.

14. Mann M, Ong SE, Gronborg M, Steen H, Jensen ON, Pandey A. Analysis of protein phosphorylation using mass spectrometry: deciphering the phosphoproteome. Trends Biotechnol. 2002;20(6):261–8. doi: 10.1016/s0167-7799(02)01944-3. PubMed PMID: 12007495.

15. Zhong Q, Xiao X, Qiu Y, Xu Z, Chen C, Chong B, Zhao X, Hai S, Li S, An Z, Dai L. Protein posttranslational modifications in health and diseases: Functions, regulatory mechanisms, and therapeutic implications. MedComm (2020). 2023;4(3):e261. Epub 20230502. doi: 10.1002/mco2.261. PubMed PMID: 37143582; PMCID: PMC10152985.

16. Torgersen KM, Vang T, Abrahamsen H, Yaqub S, Tasken K. Molecular mechanisms for protein kinase A-mediated modulation of immune function. Cell Signal. 2002;14(1):1–9. doi: 10.1016/s0898-6568(01)00214-5. PubMed PMID: 11747983.

17. Lim PS, Sutton CR, Rao S. Protein kinase C in the immune system: from signalling to chromatin regulation. Immunology. 2015;146(4):508–22. Epub 20151006. doi: 10.1111/imm.12510. PubMed PMID: 26194700; PMCID: PMC4693901.

18. Giorgione JR, Turco SJ, Epand RM. Transbilayer inhibition of protein kinase C by the lipophosphoglycan from Leishmania donovani. Proc Natl Acad Sci U S A. 1996;93(21):11634–9. doi: 10.1073/pnas.93.21.11634. PubMed PMID: 8876188; PMCID: PMC38110.

19. Shadab M, Ali N. Evasion of Host Defence by Leishmania donovani: Subversion of Signaling Pathways. Molecular biology international. 2011;2011:343961. doi: 10.4061/2011/343961. PubMed PMID: 22091401; PMCID: 3199940.

20. Shio MT, Hassani K, Isnard A, Ralph B, Contreras I, Gomez MA, Abu-Dayyeh I, Olivier M. Host cell signalling and leishmania mechanisms of evasion. Journal of tropical medicine. 2012;2012:819512. doi: 10.1155/2012/819512. PubMed PMID: 22131998; PMCID: 3216306.

21. Silverman JM, Clos J, de’Oliveira CC, Shirvani O, Fang Y, Wang C, Foster LJ, Reiner NE. An exosome-based secretion pathway is responsible for protein export from Leishmania and communication with macrophages. J Cell Sci. 2010;123(Pt 6):842–52. Epub 2010/02/18. doi: jcs.056465 [pii] 10.1242/jcs.056465. PubMed PMID: 20159964.

22. Rachidi N, Knippschild U, Späth GF. Dangerous Duplicity: The Dual Functions of Casein Kinase 1in Parasite Biology and Host Subversion. Frontiers in cellular and infection microbiology. 2021;11(230). doi: 10.3389/fcimb.2021.655700.

23. Douanne N, Dong G, Douanne M, Olivier M, Fernandez-Prada C. Unravelling the proteomic signature of extracellular vesicles released by drug-resistant Leishmania infantum parasites. PLoS neglected tropical diseases. 2020;14(7):e0008439. Epub 2020/07/07. doi: 10.1371/journal.pntd.0008439. PubMed PMID: 32628683; PMCID: PMC7365475.

24. Silvestre A, Shintre SS, Rachidi N. Released Parasite-Derived Kinases as Novel Targets for Antiparasitic Therapies. Front Cell Infect Microbiol. 2022;12:825458. Epub 20220217. doi: 10.3389/fcimb.2022.825458. PubMed PMID: 35252034; PMCID: PMC8893276.

25. Ueno N, Wilson ME. Receptor-mediated phagocytosis of Leishmania: implications for intracellular survival. Trends Parasitol. 2012;28(8):335–44. doi: 10.1016/j.pt.2012.05.002. PubMed PMID: 22726697; PMCID: PMC3399048.

26. Poullet P, Carpentier S, Barillot E. myProMS, a web server for management and validation of mass spectrometry-based proteomic data. Proteomics. 2007;7(15):2553–6. doi: 10.1002/pmic.200600784. PubMed PMID: 17610305.

27. Zhou Y, Zhou B, Pache L, Chang M, Khodabakhshi AH, Tanaseichuk O, Benner C, Chanda SK. Metascape provides a biologist-oriented resource for the analysis of systems-level datasets. Nat Commun. 2019;10(1):1523. doi: 10.1038/s41467-019-09234-6. PubMed PMID: 30944313; PMCID: PMC6447622.

28. Fernandes MC, Dillon LA, Belew AT, Bravo HC, Mosser DM, El-Sayed NM. Dual Transcriptome Profiling of Leishmania-Infected Human Macrophages Reveals Distinct Reprogramming Signatures. mBio. 2016;7(3). Epub 20160510. doi: 10.1128/mBio.00027-16. PubMed PMID: 27165796; PMCID: PMC4959658.

29. Johnson JL, Yaron TM, Huntsman EM, Kerelsky A, Song J, Regev A, Lin TY, Liberatore K, Cizin DM, Cohen BM, Vasan N, Ma Y, Krismer K, Robles JT, van de Kooij B, van Vlimmeren AE, Andrée-Busch N, Käufer NF, Dorovkov MV, Ryazanov AG, Takagi Y, Kastenhuber ER, Goncalves MD, Hopkins BD, Elemento O, Taatjes DJ, Maucuer A, Yamashita A, Degterev A, Uduman M, Lu J, Landry SD, Zhang B, Cossentino I, Linding R, Blenis J, Hornbeck PV, Turk BE, Yaffe MB, Cantley LC. An atlas of substrate specificities for the human serine/threonine kinome. Nature. 2023;613(7945):759–66. Epub 20230111. doi: 10.1038/s41586-022-05575-3. PubMed PMID: 36631611; PMCID: PMC9876800.

30. Calegari-Silva TC, Vivarini AC, Miqueline M, Dos Santos GR, Teixeira KL, Saliba AM, Nunes de Carvalho S, de Carvalho L, Lopes UG. The human parasite Leishmania amazonensis downregulates iNOS expression via NF-kappaB p50/p50 homodimer: role of the PI3K/Akt pathway. Open Biol. 2015;5(9):150118. doi: 10.1098/rsob.150118. PubMed PMID: 26400473; PMCID: PMC4593669.

31. Kumar V, Kumar A, Das S, Kumar A, Abhishek K, Verma S, Mandal A, Singh RK, Das P. Leishmania donovani Activates Hypoxia Inducible Factor-1alpha and miR-210 for Survival in Macrophages by Downregulation of NF-kappaB Mediated Pro-inflammatory Immune Response. Front Microbiol. 2018;9:385. Epub 20180308. doi: 10.3389/fmicb.2018.00385. PubMed PMID: 29568285; PMCID: PMC5852103.

32. Chakraborty A, Kurati SP, Mahata SK, Sundar S, Roy S, Sen M. Wnt5a Signaling Promotes Host Defense against Leishmania donovani Infection. J Immunol. 2017;199(3):992–1002. Epub 20170628. doi: 10.4049/jimmunol.1601927. PubMed PMID: 28659356.

33. Maity S, Chakraborty A, Mahata SK, Roy S, Das AK, Sen M. Wnt5A Signaling Blocks Progression of Experimental Visceral Leishmaniasis. Front Immunol. 2022;13:818266. Epub 20220207. doi: 10.3389/fimmu.2022.818266. PubMed PMID: 35197983; PMCID: PMC8859155.

34. Barik S, Goswami S, Nanda PK, Sarkar A, Saha B, Sarkar A, Bhattacharjee S. TGF-beta plays dual roles in immunity and pathogenesis in leishmaniasis. Cytokine. 2025;187:156865. Epub 20250127. doi: 10.1016/j.cyto.2025.156865. PubMed PMID: 39874938.

35. Barral-Netto M, Barral A, Brownell CE, Skeiky YA, Ellingsworth LR, Twardzik DR, Reed SG. Transforming growth factor-beta in leishmanial infection: a parasite escape mechanism. Science. 1992;257(5069):545–8. doi: 10.1126/science.1636092. PubMed PMID: 1636092.

36. Ruhland A, Leal N, Kima PE. Leishmania promastigotes activate PI3K/Akt signalling to confer host cell resistance to apoptosis. Cell Microbiol. 2007;9(1):84–96. Epub 20060802. doi: 10.1111/j.1462-5822.2006.00769.x. PubMed PMID: 16889626.

37. Singhal J, Madan E, Chaurasiya A, Srivastava P, Singh N, Kaushik S, Kahlon AK, Maurya MK, Marothia M, Joshi P, Ranganathan A, Singh S. Host SUMOylation Pathway Negatively Regulates Protective Immune Responses and Promotes Leishmania donovani Survival. Frontiers in cellular and infection microbiology. 2022;12:878136. doi: 10.3389/fcimb.2022.878136. PubMed PMID: 35734580; PMCID: 9207379.

38. Gupta AK, Das S, Kamran M, Ejazi SA, Ali N. The pathogenicity and virulence of Leishmania - interplay of virulence factors with host defenses. Virulence. 2022;13(1):903–35. doi: 10.1080/21505594.2022.2074130. PubMed PMID: 35531875; PMCID: PMC9154802.

39. Yang Z, Mosser DM, Zhang X. Activation of the MAPK, ERK, following Leishmania amazonensis infection of macrophages. J Immunol. 2007;178(2):1077–85. doi: 10.4049/jimmunol.178.2.1077. PubMed PMID: 17202371; PMCID: PMC2643020.

40. Junghae M, Raynes JG. Activation of p38 mitogen-activated protein kinase attenuates Leishmania donovani infection in macrophages. Infect Immun. 2002;70(9):5026–35. doi: 10.1128/IAI.70.9.5026-5035.2002. PubMed PMID: 12183549; PMCID: PMC128247.

41. Kumar GA, Karmakar J, Mandal C, Chattopadhyay A. Leishmania donovani Internalizes into Host Cells via Caveolin-mediated Endocytosis. Sci Rep. 2019;9(1):12636. doi: 10.1038/s41598-019-49007-1. PubMed PMID: 31477757; PMCID: PMC6718660.

42. Verma JK, Rastogi R, Mukhopadhyay A. Leishmania donovani resides in modified early endosomes by upregulating Rab5a expression via the downregulation of miR-494. PLoS Pathog. 2017;13(6):e1006459. doi: 10.1371/journal.ppat.1006459. PubMed PMID: 28650977; PMCID: PMC5501680.

43. Azevedo E, Oliveira LT, Castro Lima AK, Terra R, Dutra PM, Salerno VP. Interactions between Leishmania braziliensis and Macrophages Are Dependent on the Cytoskeleton and Myosin Va. J Parasitol Res. 2012;2012:275436. doi: 10.1155/2012/275436. PubMed PMID: 22792440; PMCID: PMC3391898.

44. Young J, Kima PE. The Leishmania Parasitophorous Vacuole Membrane at the Parasite-Host Interface. Yale J Biol Med. 2019;92(3):511–21. PubMed PMID: 31543712; PMCID: PMC6747952.

45. Pucadyil TJ, Tewary P, Madhubala R, Chattopadhyay A. Cholesterol is required for Leishmania donovani infection: implications in leishmaniasis. Mol Biochem Parasitol. 2004;133(2):145–52. doi: 10.1016/j.molbiopara.2003.10.002. PubMed PMID: 14698427.

46. Zhang H. Lysosomal acid lipase and lipid metabolism: new mechanisms, new questions, and new therapies. Curr Opin Lipidol. 2018;29(3):218–23. doi: 10.1097/MOL.0000000000000507. PubMed PMID: 29547398; PMCID: PMC6215475.

47. Desjardins M, Descoteaux A. Inhibition of phagolysosomal biogenesis by the Leishmania lipophosphoglycan. J Exp Med. 1997;185(12):2061–8. doi: 10.1084/jem.185.12.2061. PubMed PMID: 9182677; PMCID: PMC2196352.

48. Scianimanico S, Desrosiers M, Dermine JF, Meresse S, Descoteaux A, Desjardins M. Impaired recruitment of the small GTPase rab7 correlates with the inhibition of phagosome maturation by Leishmania donovani promastigotes. Cell Microbiol. 1999;1(1):19–32. doi: 10.1046/j.1462-5822.1999.00002.x. PubMed PMID: 11207538.

49. Michels AA, Bensaude O. Hexim1, an RNA-controlled protein hub. Transcription. 2018;9(4):262–71. Epub 20180223. doi: 10.1080/21541264.2018.1429836. PubMed PMID: 29345523; PMCID: PMC6104690.

50. Klein DJ, Moore PB, Steitz TA. The roles of ribosomal proteins in the structure assembly, and evolution of the large ribosomal subunit. J Mol Biol. 2004;340(1):141–77. doi: 10.1016/j.jmb.2004.03.076. PubMed PMID: 15184028.

51. Alkhateeb AA, Connor JR. Nuclear ferritin: A new role for ferritin in cell biology. Biochim Biophys Acta. 2010;1800(8):793–7. doi: 10.1016/j.bbagen.2010.03.017. PubMed PMID: 20347012.

52. Plays M, Muller S, Rodriguez R. Chemistry and biology of ferritin. Metallomics. 2021;13(5). doi: 10.1093/mtomcs/mfab021. PubMed PMID: 33881539; PMCID: PMC8083198.

53. Andrews SC, Arosio P, Bottke W, Briat JF, von Darl M, Harrison PM, Laulhere JP, Levi S, Lobreaux S, Yewdall SJ. Structure, function, and evolution of ferritins. J Inorg Biochem. 1992;47(3-4):161–74. doi: 10.1016/0162-0134(92)84062-r. PubMed PMID: 1431878.

54. Ru Q, Li Y, Chen L, Wu Y, Min J, Wang F. Iron homeostasis and ferroptosis in human diseases: mechanisms and therapeutic prospects. Signal Transduct Target Ther. 2024;9(1):271. doi: 10.1038/s41392-024-01969-z. PubMed PMID: 39396974; PMCID: PMC11486532.

55. Malafaia G, Marcon Lde N, Pereira Lde F, Pedrosa ML, Rezende SA. Leishmania chagasi: effect of the iron deficiency on the infection in BALB/c mice. Exp Parasitol. 2011;127(3):719–23. Epub 20101124. doi: 10.1016/j.exppara.2010.11.010. PubMed PMID: 21110973.

56. Vale-Costa S, Gomes-Pereira S, Teixeira CM, Rosa G, Rodrigues PN, Tomas A, Appelberg R, Gomes MS. Iron overload favors the elimination of Leishmania infantum from mouse tissues through interaction with reactive oxygen and nitrogen species. PLoS neglected tropical diseases. 2013;7(2):e2061. Epub 20130214. doi: 10.1371/journal.pntd.0002061. PubMed PMID: 23459556; PMCID: PMC3573095.

57. Torti FM, Torti SV. Regulation of ferritin genes and protein. Blood. 2002;99(10):3505–16. doi: 10.1182/blood.v99.10.3505. PubMed PMID: 11986201.

58. Asano T, Komatsu M, Yamaguchi-Iwai Y, Ishikawa F, Mizushima N, Iwai K. Distinct mechanisms of ferritin delivery to lysosomes in iron-depleted and iron-replete cells. Mol Cell Biol. 2011;31(10):2040–52. doi: 10.1128/MCB.01437-10. PubMed PMID: 21444722; PMCID: PMC3133360.

59. Zhang Y, Mikhael M, Xu D, Li Y, Soe-Lin S, Ning B, Li W, Nie G, Zhao Y, Ponka P. Lysosomal proteolysis is the primary degradation pathway for cytosolic ferritin and cytosolic ferritin degradation is necessary for iron exit. Antioxid Redox Signal. 2010;13(7):999–1009. doi: 10.1089/ars.2010.3129. PubMed PMID: 20406137.

60. Antoine JC, Prina E, Lang T, Courret N. The biogenesis and properties of the parasitophorous vacuoles that harbour Leishmania in murine macrophages. Trends Microbiol. 1998;6(10):392–401. doi: 10.1016/s0966-842x(98)01324-9. PubMed PMID: 9807783.

61. Surguladze N, Thompson KM, Beard JL, Connor JR, Fried MG. Interactions and reactions of ferritin with DNA. The Journal of biological chemistry. 2004;279(15):14694–702. Epub 20040120. doi: 10.1074/jbc.M313348200. PubMed PMID: 14734543.

62. Pountney D, Trugnan G, Bourgeois M, Beaumont C. The identification of ferritin in the nucleus of K562 cells, and investigation of a possible role in the transcriptional regulation of adult beta-globin gene expression. J Cell Sci. 1999;112 (Pt 6):825–31. doi: 10.1242/jcs.112.6.825. PubMed PMID: 10036232.

63. Srivastava AK, Reutovich AA, Hunter NJ, Arosio P, Bou-Abdallah F. Ferritin microheterogeneity, subunit composition, functional, and physiological implications. Sci Rep. 2023;13(1):19862. doi: 10.1038/s41598-023-46880-9. PubMed PMID: 37963965; PMCID: PMC10646083.

64. Ahmad S, Moriconi F, Naz N, Sultan S, Sheikh N, Ramadori G, Malik IA. Ferritin L and Ferritin H are differentially located within hepatic and extra hepatic organs under physiological and acute phase conditions. Int J Clin Exp Pathol. 2013;6(4):622–9. PubMed PMID: 23573308; PMCID: PMC3606851.

65. Tofighi Naeem A, Mahmoudi S, Saboui F, Hajjaran H, Pourakbari B, Mohebali M, Zarkesh MR, Mamishi S. Clinical Features and Laboratory Findings of Visceral Leishmaniasis in Children Referred To Children Medical Center Hospital, Tehran, Iran during 2004-2011. Iranian journal of parasitology. 2014;9(1):1–5. PubMed PMID: 25642253; PMCID: 4289866.

66. Garcia MSA, Oliveira Filho VA, Brioschi MBC, Minori K, Miguel DC. Leishmania (Leishmania) amazonensis. Trends Parasitol. 2025;41(1):66-7. doi: 10.1016/j.pt.2024.10.023. PubMed PMID: 39572329.

67. Russell DG, Xu S, Chakraborty P. Intracellular trafficking and the parasitophorous vacuole of Leishmania mexicana-infected macrophages. J Cell Sci. 1992;103 (Pt 4):1193–210. doi: 10.1242/jcs.103.4.1193. PubMed PMID: 1487496.

68. Dowdle WE, Nyfeler B, Nagel J, Elling RA, Liu S, Triantafellow E, Menon S, Wang Z, Honda A, Pardee G, Cantwell J, Luu C, Cornella-Taracido I, Harrington E, Fekkes P, Lei H, Fang Q, Digan ME, Burdick D, Powers AF, Helliwell SB, D’Aquin S, Bastien J, Wang H, Wiederschain D, Kuerth J, Bergman P, Schwalb D, Thomas J, Ugwonali S, Harbinski F, Tallarico J, Wilson CJ, Myer VE, Porter JA, Bussiere DE, Finan PM, Labow MA, Mao X, Hamann LG, Manning BD, Valdez RA, Nicholson T, Schirle M, Knapp MS, Keaney EP, Murphy LO. Selective VPS34 inhibitor blocks autophagy and uncovers a role for NCOA4 in ferritin degradation and iron homeostasis in vivo. Nat Cell Biol. 2014;16(11):1069–79. doi: 10.1038/ncb3053. PubMed PMID: 25327288.

69. Mancias JD, Wang X, Gygi SP, Harper JW, Kimmelman AC. Quantitative proteomics identifies NCOA4 as the cargo receptor mediating ferritinophagy. Nature. 2014;509(7498):105–9. doi: 10.1038/nature13148. PubMed PMID: 24695223; PMCID: PMC4180099.

70. Mancias JD, Pontano Vaites L, Nissim S, Biancur DE, Kim AJ, Wang X, Liu Y, Goessling W, Kimmelman AC, Harper JW. Ferritinophagy via NCOA4 is required for erythropoiesis and is regulated by iron dependent HERC2-mediated proteolysis. Elife. 2015;4. doi: 10.7554/eLife.10308. PubMed PMID: 26436293; PMCID: PMC4592949.

71. Le Y, Liu Q, Yang Y, Wu J. The emerging role of nuclear receptor coactivator 4 in health and disease: a novel bridge between iron metabolism and immunity. Cell Death Discov. 2024;10(1):312. doi: 10.1038/s41420-024-02075-3. PubMed PMID: 38961066; PMCID: PMC11222541.

72. Goodwin JM, Dowdle WE, DeJesus R, Wang Z, Bergman P, Kobylarz M, Lindeman A, Xavier RJ, McAllister G, Nyfeler B, Hoffman G, Murphy LO. Autophagy-Independent Lysosomal Targeting Regulated by ULK1/2-FIP200 and ATG9. Cell Rep. 2017;20(10):2341–56. doi: 10.1016/j.celrep.2017.08.034. PubMed PMID: 28877469; PMCID: PMC5699710.

73. Kuno S, Fujita H, Tanaka YK, Ogra Y, Iwai K. Iron-induced NCOA4 condensation regulates ferritin fate and iron homeostasis. EMBO Rep. 2022;23(5):e54278. doi: 10.15252/embr.202154278. PubMed PMID: 35318808; PMCID: PMC9066066.

74. Kidane TZ, Sauble E, Linder MC. Release of iron from ferritin requires lysosomal activity. Am J Physiol Cell Physiol. 2006;291(3):C445–55. doi: 10.1152/ajpcell.00505.2005. PubMed PMID: 16611735.

75. Goto Y, Ito T, Ghosh S, Mukherjee B. Access and utilization of host-derived iron by Leishmania parasites. J Biochem. 2023;175(1):17–24. doi: 10.1093/jb/mvad082. PubMed PMID: 37830941; PMCID: PMC10771036.

76. Stockwell BR, Friedmann Angeli JP, Bayir H, Bush AI, Conrad M, Dixon SJ, Fulda S, Gascon S, Hatzios SK, Kagan VE, Noel K, Jiang X, Linkermann A, Murphy ME, Overholtzer M, Oyagi A, Pagnussat GC, Park J, Ran Q, Rosenfeld CS, Salnikow K, Tang D, Torti FM, Torti SV, Toyokuni S, Woerpel KA, Zhang DD. Ferroptosis: A Regulated Cell Death Nexus Linking Metabolism, Redox Biology, and Disease. Cell. 2017;171(2):273–85. doi: 10.1016/j.cell.2017.09.021. PubMed PMID: 28985560; PMCID: PMC5685180.

77. Stockwell BR. Ferroptosis turns 10: Emerging mechanisms, physiological functions, and therapeutic applications. Cell. 2022;185(14):2401–21. doi: 10.1016/j.cell.2022.06.003. PubMed PMID: 35803244; PMCID: PMC9273022.

78. Ben-Othman R, Flannery AR, Miguel DC, Ward DM, Kaplan J, Andrews NW. Leishmania-mediated inhibition of iron export promotes parasite replication in macrophages. PLoS pathogens. 2014;10(1):e1003901. Epub 20140130. doi: 10.1371/journal.ppat.1003901. PubMed PMID: 24497831; PMCID: PMC3907422.

79. Sun X, Ou Z, Xie M, Kang R, Fan Y, Niu X, Wang H, Cao L, Tang D. HSPB1 as a novel regulator of ferroptotic cancer cell death. Oncogene. 2015;34(45):5617–25. Epub 20150302. doi: 10.1038/onc.2015.32. PubMed PMID: 25728673; PMCID: PMC4640181.

80. Chen H, Zheng C, Zhang Y, Chang YZ, Qian ZM, Shen X. Heat shock protein 27 downregulates the transferrin receptor 1-mediated iron uptake. Int J Biochem Cell Biol. 2006;38(8):1402–16. Epub 20060307. doi: 10.1016/j.biocel.2006.02.006. PubMed PMID: 16546437.

81. Forget G, Gregory DJ, Whitcombe LA, Olivier M. Role of host protein tyrosine phosphatase SHP-1 in Leishmania donovani-induced inhibition of nitric oxide production. Infect Immun. 2006;74(11):6272–9. doi: 10.1128/IAI.00853-05. PubMed PMID: 17057094; PMCID: PMC1695482.

82. Forget G, Gregory DJ, Olivier M. Proteasome-mediated degradation of STAT1alpha following infection of macrophages with Leishmania donovani. The Journal of biological chemistry. 2005;280(34):30542–9. Epub 20050627. doi: 10.1074/jbc.M414126200. PubMed PMID: 15983048.

83. Matte C, Descoteaux A. Leishmania donovani amastigotes impair gamma interferon-induced STAT1alpha nuclear translocation by blocking the interaction between STAT1alpha and importin-alpha5. Infect Immun. 2010;78(9):3736–43. doi: 10.1128/IAI.00046-10. PubMed PMID: 20566692; PMCID: PMC2937469.

84. Descoteaux A, Matlashewski G, Turco SJ. Inhibition of macrophage protein kinase C-mediated protein phosphorylation by Leishmania donovani lipophosphoglycan. J Immunol. 1992;149(9):3008–15. PubMed PMID: 1383336.

85. van Bruggen R, Drewniak A, Tool AT, Jansen M, van Houdt M, Geissler J, van den Berg TK, Chapel H, Kuijpers TW. Toll-like receptor responses in IRAK-4-deficient neutrophils. J Innate Immun. 2010;2(3):280–7. Epub 20091216. doi: 10.1159/000268288. PubMed PMID: 20375545.

86. Atayde VD, Aslan H, Townsend S, Hassani K, Kamhawi S, Olivier M. Exosome Secretion by the Parasitic Protozoan Leishmania within the Sand Fly Midgut. Cell Rep. 2015;13(5):957–67. doi: 10.1016/j.celrep.2015.09.058. PubMed PMID: 26565909; PMCID: 4644496.

87. Forrest DM, Batista M, Marchini FK, Tempone AJ, Traub-Csekö YM. Proteomic analysis of exosomes derived from procyclic and metacyclic-like cultured Leishmania infantum chagasi. Journal of Proteomics. 2020;227:103902. doi: 10.1016/j.jprot.2020.103902.

88. Liu J, Carvalho LP, Bhattachariya S, Carbone CJ, Kumar KG, Leu NA, Yau PM, Donald RG, Weiss MJ, Baker DP, McLaughlin KJ, Scott P, Fuchs SY. Mammalian casein kinase 1alpha and its leishmanial ortholog regulate stability of IFNAR1 and Type I interferon signaling. Mol Cell Biol. 2009;29(24):6401–12. Epub 2009/10/07. doi: MCB.00478-09 [pii] 10.1128/MCB.00478-09. PubMed PMID: 19805514.

89. Smirlis D, Dingli F, Sabatet V, Roth A, Knippschild U, Loew D, Spath GF, Rachidi N. Identification of the Host Substratome of Leishmania-Secreted Casein Kinase 1 Using a SILAC-Based Quantitative Mass Spectrometry Assay. Front Cell Dev Biol. 2021;9:800098. Epub 20220103. doi: 10.3389/fcell.2021.800098. PubMed PMID: 35047509; PMCID: PMC8762337.

90. Nandan D, Yi T, Lopez M, Lai C, Reiner NE. Leishmania EF-1alpha activates the Src homology 2 domain containing tyrosine phosphatase SHP-1 leading to macrophage deactivation. The Journal of biological chemistry. 2002;277(51):50190–7. Epub 20021015. doi: 10.1074/jbc.M209210200. PubMed PMID: 12384497.

91. Abu-Dayyeh I, Shio MT, Sato S, Akira S, Cousineau B, Olivier M. Leishmania-induced IRAK-1 inactivation is mediated by SHP-1 interacting with an evolutionarily conserved KTIM motif. PLoS Negl Trop Dis. 2008;2(12):e305. doi: 10.1371/journal.pntd.0000305. PubMed PMID: 19104650; PMCID: PMC2596967.

92. Chaparro V, Leroux LP, Masvidal L, Lorent J, Graber TE, Zimmermann A, Arango Duque G, Descoteaux A, Alain T, Larsson O, Jaramillo M. Translational profiling of macrophages infected with Leishmania donovani identifies mTOR- and eIF4A-sensitive immune-related transcripts. PLoS Pathog. 2020;16(6):e1008291. doi: 10.1371/journal.ppat.1008291. PubMed PMID: 32479529; PMCID: PMC7310862.

93. Jaramillo M, Gomez MA, Larsson O, Shio MT, Topisirovic I, Contreras I, Luxenburg R, Rosenfeld A, Colina R, McMaster RW, Olivier M, Costa-Mattioli M, Sonenberg N. Leishmania repression of host translation through mTOR cleavage is required for parasite survival and infection. Cell Host Microbe. 2011;9(4):331–41. doi: 10.1016/j.chom.2011.03.008. PubMed PMID: 21501832.

94. Das NK, Sandhya S, G VV, Kumar R, Singh AK, Bal SK, Kumari S, Mukhopadhyay CK. Leishmania donovani inhibits ferroportin translation by modulating FBXL5-IRP2 axis for its growth within host macrophages. Cell Microbiol. 2018;20(7):e12834. doi: 10.1111/cmi.12834. PubMed PMID: 29470856.

95. Levi S, Santambrogio P, Cozzi A, Rovida E, Corsi B, Tamborini E, Spada S, Albertini A, Arosio P. The role of the L-chain in ferritin iron incorporation. Studies of homo and heteropolymers. J Mol Biol. 1994;238(5):649–54. doi: 10.1006/jmbi.1994.1325. PubMed PMID: 8182740.

96. Aolymat I, Hatmal MM, Olaimat AN. The Emerging Role of Heat Shock Factor 1 (HSF1) and Heat Shock Proteins (HSPs) in Ferroptosis. Pathophysiology. 2023;30(1):63–82. doi: 10.3390/pathophysiology30010007. PubMed PMID: 36976734; PMCID: PMC10057451.

97. Murdoch CC, Skaar EP. Nutritional immunity: the battle for nutrient metals at the host-pathogen interface. Nature reviews Microbiology. 2022;20(11):657–70. Epub 20220531. doi: 10.1038/s41579-022-00745-6. PubMed PMID: 35641670; PMCID: PMC9153222.

98. Ullah I, Lang M. Key players in the regulation of iron homeostasis at the host-pathogen interface. Front Immunol. 2023;14:1279826. Epub 20231024. doi: 10.3389/fimmu.2023.1279826. PubMed PMID: 37942316; PMCID: PMC10627961.

99. Shankar G, Akhter Y. Stealing survival: Iron acquisition strategies of Mycobacteriumtuberculosis. Biochimie. 2024;227(Pt A):37–60. doi: 10.1016/j.biochi.2024.06.006. PubMed PMID: 38901792.

100. Ryndak MB, Wang S, Smith I, Rodriguez GM. The Mycobacterium tuberculosis high-affinity iron importer, IrtA, contains an FAD-binding domain. J Bacteriol. 2010;192(3):861–9. doi: 10.1128/JB.00223-09. PubMed PMID: 19948799; PMCID: PMC2812465.

101. Takemura K, Kolasinski V, Del Poeta M, Vieira de Sa NF, Garg A, Ojima I, Del Poeta M, Pereira de Sa N. Iron acquisition strategies in pathogenic fungi. mBio. 2025;16(6):e0121125. Epub 20250520. doi: 10.1128/mbio.01211-25. PubMed PMID: 40391928; PMCID: PMC12153323.

102. Almeida RS, Brunke S, Albrecht A, Thewes S, Laue M, Edwards JE, Filler SG, Hube B. the hyphal-associated adhesin and invasin Als3 of Candida albicans mediates iron acquisition from host ferritin. PLoS pathogens. 2008;4(11):e1000217. Epub 20081121. doi: 10.1371/journal.ppat.1000217. PubMed PMID: 19023418; PMCID: PMC2581891.

103. Galvez N, Ruiz B, Cuesta R, Colacio E, Dominguez-Vera JM. Release of iron from ferritin by aceto- and benzohydroxamic acids. Inorg Chem. 2005;44(8):2706–9. doi: 10.1021/ic048840s. PubMed PMID: 15819556.

104. Flannery AR, Huynh C, Mittra B, Mortara RA, Andrews NW. LFR1 ferric iron reductase of Leishmania amazonensis is essential for the generation of infective parasite forms. The Journal of biological chemistry. 2011;286(26):23266–79. Epub 20110510. doi: 10.1074/jbc.M111.229674. PubMed PMID: 21558274; PMCID: PMC3123093.

105. Wolff A, de Nechaud B, Chillet D, Mazarguil H, Desbruyeres E, Audebert S, Edde B, Gros F, Denoulet P. Distribution of glutamylated alpha and beta-tubulin in mouse tissues using a specific monoclonal antibody, GT335. Eur J Cell Biol. 1992;59(2):425-32. PubMed PMID: 1493808.

106. Schindelin J, Arganda-Carreras I, Frise E, Kaynig V, Longair M, Pietzsch T, Preibisch S, Rueden C, Saalfeld S, Schmid B, Tinevez JY, White DJ, Hartenstein V, Eliceiri K, Tomancak P, Cardona A. Fiji: an open-source platform for biological-image analysis. Nature methods. 2012;9(7):676–82. doi: 10.1038/nmeth.2019. PubMed PMID: 22743772; PMCID: 3855844.

107. Pescher P, Blisnick T, Bastin P, Spath GF. Quantitative proteome profiling informs on phenotypic traits that adapt Leishmania donovani for axenic and intracellular proliferation. Cell Microbiol. 2011;13(7):978–91. doi: 10.1111/j.1462-5822.2011.01593.x. PubMed PMID: 21501362.

108. Prieto Barja P, Pescher P, Bussotti G, Dumetz F, Imamura H, Kedra D, Domagalska M, Chaumeau V, Himmelbauer H, Pages M, Sterkers Y, Dujardin JC, Notredame C, Spath GF. Haplotype selection as an adaptive mechanism in the protozoan pathogen Leishmania donovani. Nat Ecol Evol. 2017;1(12):1961–9. doi: 10.1038/s41559-017-0361-x. PubMed PMID: 29109466.

109. 109. The M, MacCoss MJ, Noble WS, Kall L. Fast and Accurate Protein False Discovery Rates on Large-Scale Proteomics Data Sets with Percolator 3.0. J Am Soc Mass Spectrom. 2016;27(11):1719–27. Epub 20160829. doi: 10.1007/s13361-016-1460-7. PubMed PMID: 27572102; PMCID: PMC5059416.

110. Valot B, Langella O, Nano E, Zivy M. MassChroQ: a versatile tool for mass spectrometry quantification. Proteomics. 2011;11(17):3572–7. Epub 20110804. doi: 10.1002/pmic.201100120. PubMed PMID: 21751374.

111. Perez-Riverol Y, Bandla C, Kundu DJ, Kamatchinathan S, Bai J, Hewapathirana S, John NS, Prakash A, Walzer M, Wang S, Vizcaino JA. The PRIDE database at 20 years: 2025 update. Nucleic acids research. 2025;53(D1):D543-D53. doi: 10.1093/nar/gkae1011. PubMed PMID: 39494541; PMCID: PMC11701690.

112. Yaron-Barir TM, Joughin BA, Huntsman EM, Kerelsky A, Cizin DM, Cohen BM, Regev A, Song J, Vasan N, Lin TY, Orozco JM, Schoenherr C, Sagum C, Bedford MT, Wynn RM, Tso SC, Chuang DT, Li L, Li SS, Creixell P, Krismer K, Takegami M, Lee H, Zhang B, Lu J, Cossentino I, Landry SD, Uduman M, Blenis J, Elemento O, Frame MC, Hornbeck PV, Cantley LC, Turk BE, Yaffe MB, Johnson JL. The intrinsic substrate specificity of the human tyrosine kinome. Nature. 2024;629(8014):1174–81. doi: 10.1038/s41586-024-07407-y. PubMed PMID: 38720073; PMCID: PMC11136658 and is a founder and receives research support from Petra Pharmaceuticals; is listed as an inventor on a patent (WO2019232403A1, Weill Cornell Medicine) for combination therapy for PI3K-associated disease or disorder, and the identification of therapeutic interventions to improve response to PI3K inhibitors for cancer treatment; is a co-founder and shareholder in Faeth Therapeutics; has equity in and consults for Cell Signaling Technologies, Volastra, Larkspur and 1 Base Pharmaceuticals; and consults for Loxo-Lilly. J.L.J. has received consulting fees from Scorpion Therapeutics and Volastra Therapeutics. T.M.Y.-B. is a co-founder of DeStroke. N.V. reports consulting activities for Novartis and is on the scientific advisory board of Heligenics. O.E. is a founder and equity holder of Volastra Therapeutics and OneThree Biotech; is a member of the scientific advisory board of Owkin, Freenome, Genetic Intelligence, Acuamark and Champions Oncology; and receives research support from Eli Lilly, Janssen and Sanofi. M.T.B. is the co-founder of EpiCypher. The other authors declare no competing interests.

113. Meyerhofer N, Krogan NJ, Polacco BJ, Blumenthal DB. Inference of differential kinase interaction networks with KINference. Bioinformatics. 2025;41(7). doi: 10.1093/bioinformatics/btaf349. PubMed PMID: 40579228; PMCID: PMC12270260.

114. Louvel V, Haase R, Mercey O, Laporte MH, Eloy T, Baudrier E, Fortun D, Soldati-Favre D, Hamel V, Guichard P. iU-ExM: nanoscopy of organelles and tissues with iterative ultrastructure expansion microscopy. Nat Commun. 2023;14(1):7893. doi: 10.1038/s41467-023-43582-8. PubMed PMID: 38036510; PMCID: PMC10689735.

