## Supplementary material for "Dynamic phosphoproteomics and proteomics uncover *Leishmania donovani*-driven ferritin hijacking, contributing to the control of iron homeostasis and iron-related oxidative stress": The supplementary figures are linked to the following text : Figure S1-S8 and Table S7

### Figure S1

A

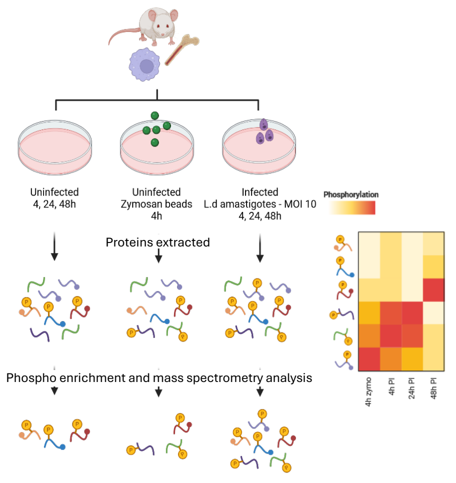


Figure 1: Experimental workflow used for phosphoproteomic analysis.

B

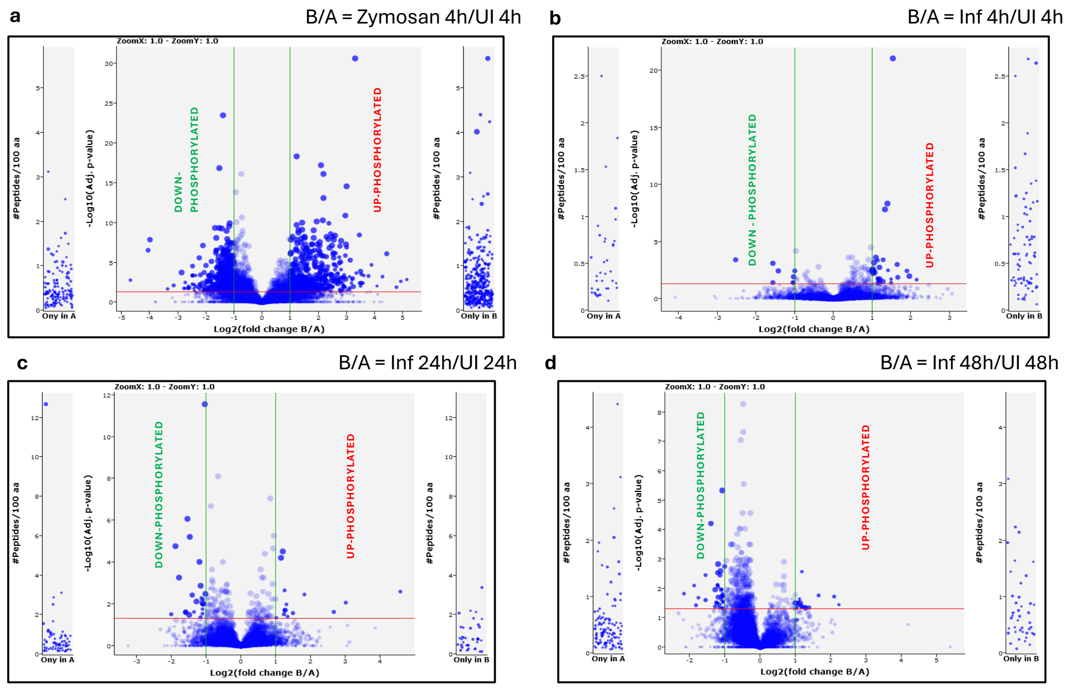


**Figure S1**: **A.** Experimental workflow used for phosphoproteomic analysis. **B.** Volcano plots for assessing the impact of phagocytosis and infection on global phosphorylation profiles. Plots for -$\log_{10}$ adjusted p-value vs $\log_2$Fold change (FC) are given for a comparison B/A. For each plot, for the central panel red line is for p-value = 0.05, green lines are for $\log_2$FC of 1 and -1,in the side panels are the phosphosites detected only in condition B (right panel) or only in condition A (left panel). For the subsections: (a) 4h Zymosan, (b) 4hpi, (c) 24hpi, (d) 48hpi.

### Figure S2

**A**

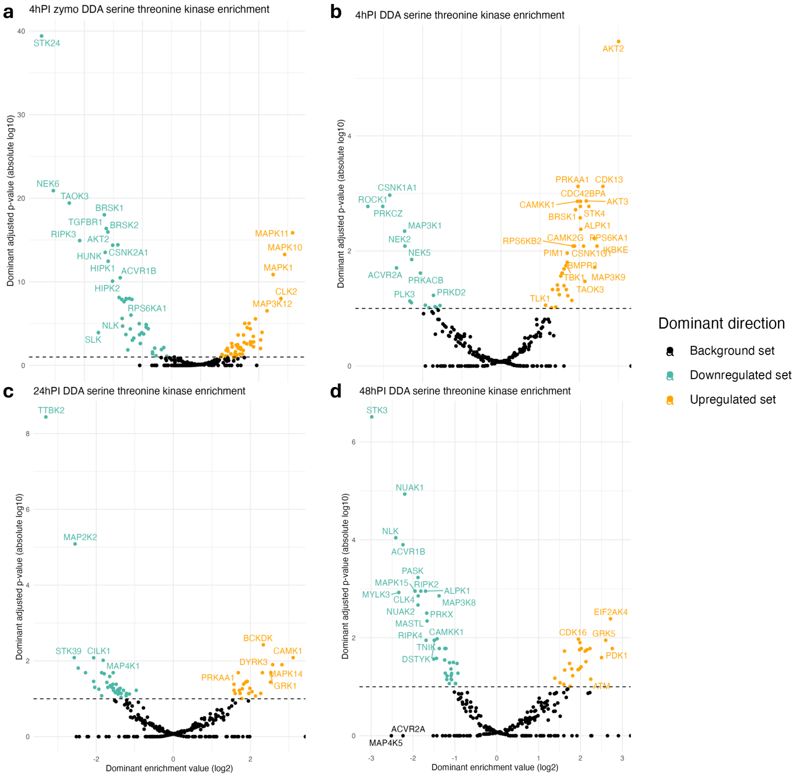


**B**

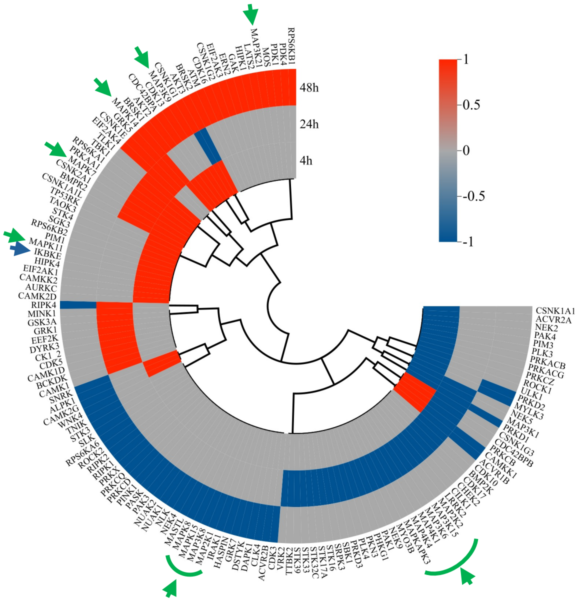


**Figure S2**: Volcano plots for the host kinase enrichment for **A.** 4h Zymosan, **B.** 4hpi, **C.** 24hpi, **D.** 48hpi. The volcano plots are plotted for absolute $\log_{10}$ adjusted p-value versus $\log_2$(enrichment value). The dotted black line is drawn at p-value=0.05. Upregulated set are the enriched active kinases, downregulated set are the enriched inactive kinases and background set are the kinases not enriched in the analysis. **E.** Modulation of all the kinase activities during infection.

### Figure S3

A

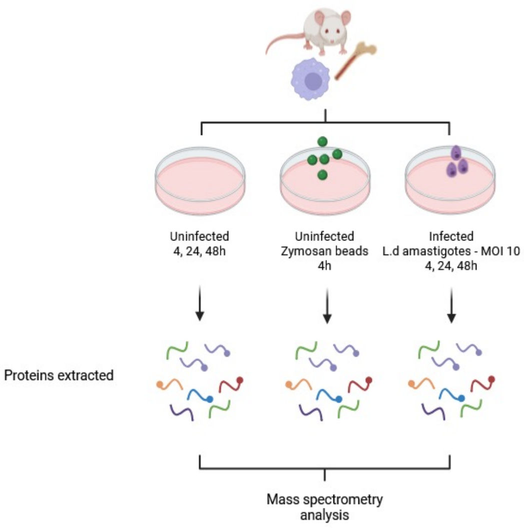


B

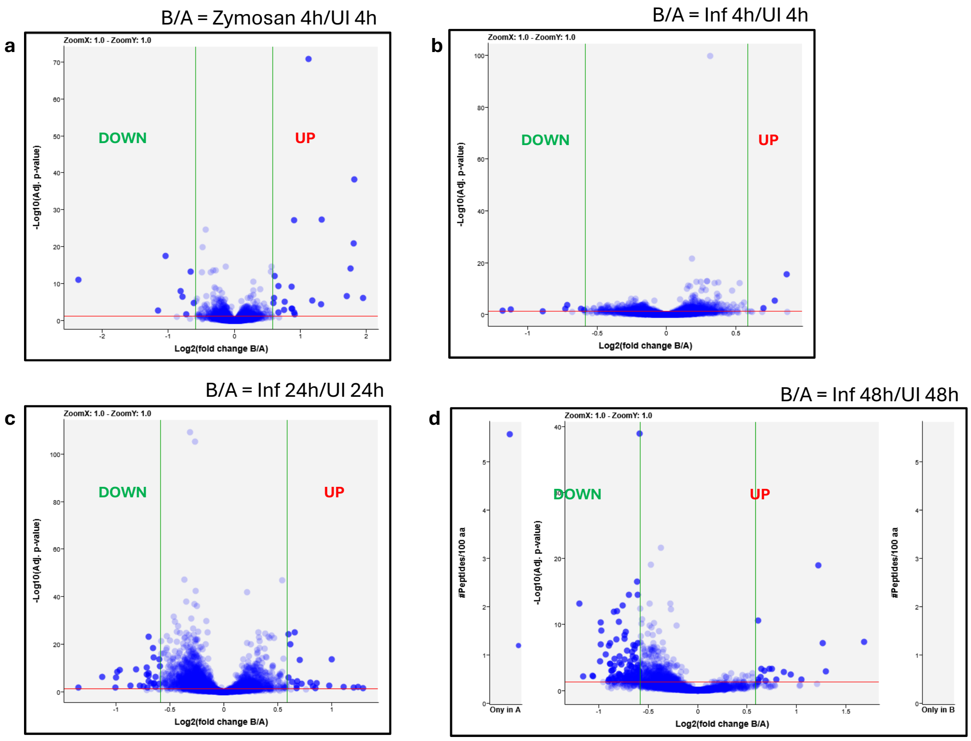


**Figure S3**: **A.** Experimental workflow used for proteomic analysis. **B.** Volcano plots assessing the impact of phagocytosis and infection on the global host protein levels. Plots for -$\log_{10}$ adjusted p-value vs $\log_2$Fold change ($\log_2$FC) are given for a comparison B/A. For each plot, for the central panel, red line is for p value = 0.05, green lines are for $\log_2$FC of 0.58 and -0.58, in the side panels are the proteins detected only in condition B (right panel) or only in condition A (left panel). For the subsections : (a) 4h Zymosan, (b) 4hpi, (c) 24hpi, (d) 48hpi.

### Figure S4

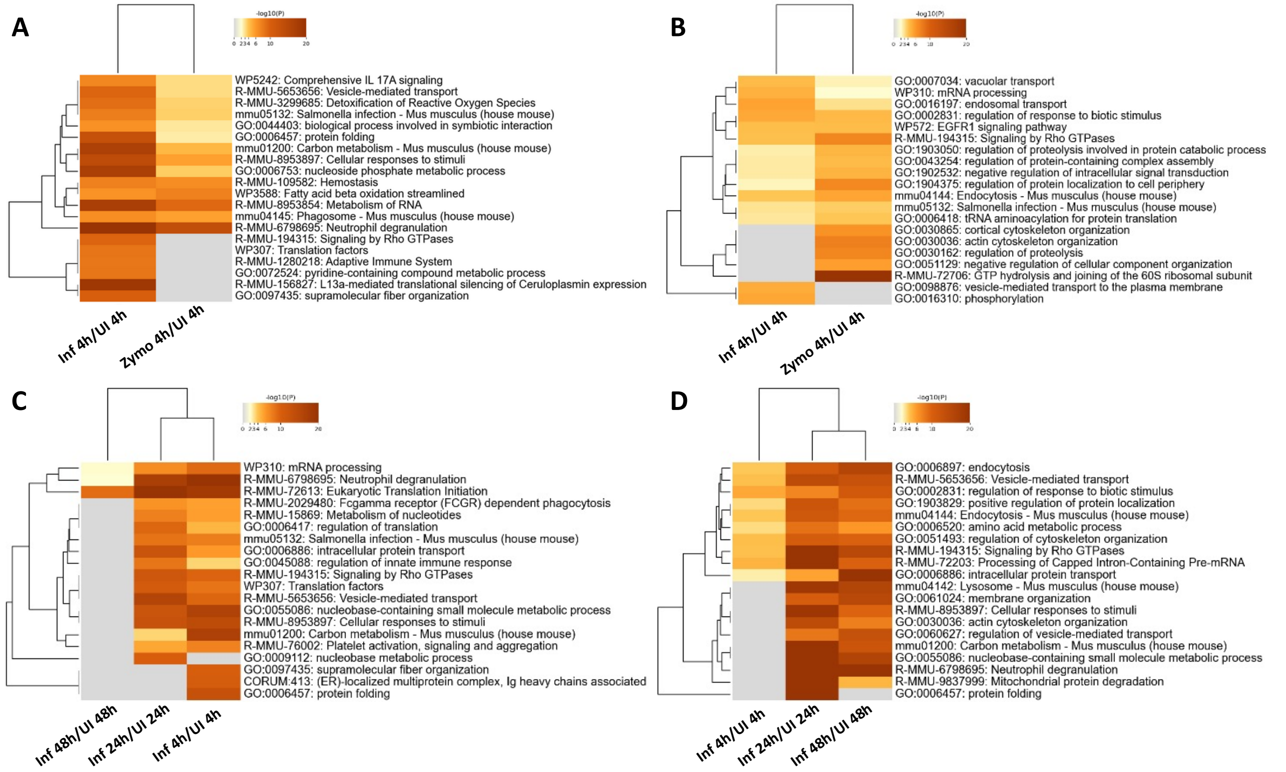


**Fig S4**: Comparing the impact of 4h Zymosan with 4hpi **A.** In upregulated proteins **B.** In downregulated proteins. Analysing the temporal modulation of host pathway during infection at 4hpi, 24hpi and 48hpi **C.** In upregulated proteins **D.** In downregulated proteins. The intensity of orange provides the level of significance of the enrichment. Grey indicates that the pathway was not enriched.

### Figure S5

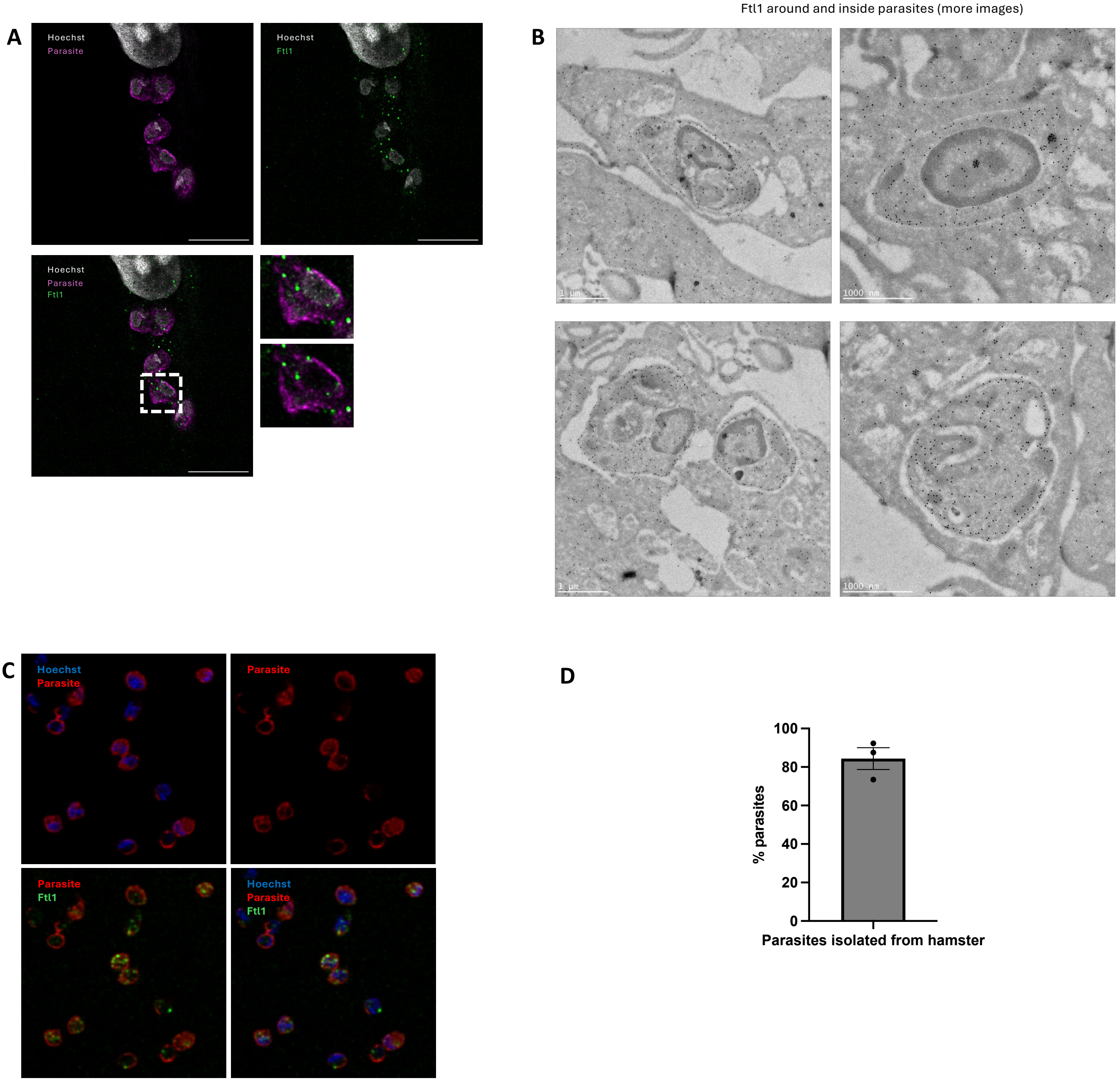


**Figure S5**: Localisation of Ftl1. **A.** Representative image from 4X immunofluorescence expansion microscopy showing Hoechst (grey), parasite using GT335 antibody (magenta), Ftl1 (green). One parasite (white box) is zoomed in to show the localisation of Ftl1 inside the parasite. **B.** More EM images showing Ftl1 localisation inside the parasites and presence of aggregates in some parasites. **C.** Representative immunofluorescence images showing Hoechst (blue), Ftl1 (green), parasite (red) for amastigotes isolated from infected liver and spleen of hamsters that were fixed and immunostained. **D.** Ftl1 localisation in purified amastigotes from hamster. Quantification of % parasites purified from hamster containing host Ftl1. Quantifications were performed by counting around 100 cells in three biological replicates. The error bars represent standard error.

### Figure S6

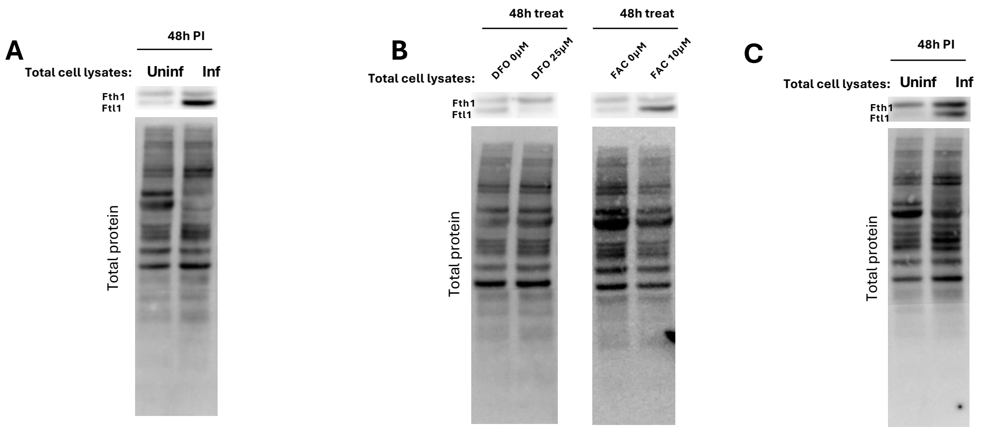


(attached to Fig 10)

**Figure S6**: Western blots for Ftl1 and Fth1 protein levels. The total protein lane intensity was used for normalising the corresponding band intensities before performing any further analysis. **A.** For BMDMs infected or not with *L. donovani* for 4hpi or 48hpi. **B.** Ftl1/Fth1 ratio for BMDMs treated or not with 25 µM DFO or 10 µM FAC for 48 hours. **C.** Ftl1/Fth1 ratio for BMDMs infected or not with *L. amazonensis* for 48h.

### Figure S7

**A**

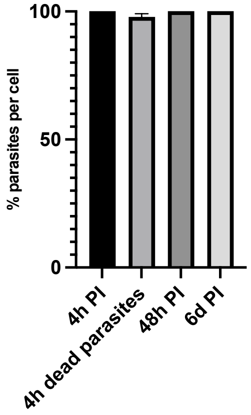


**B**

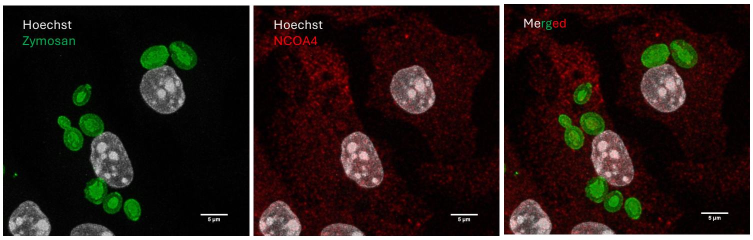


(attached to fig 10)

**Figure S7**: Localisation of NCOA4 in 4h Zymosan treatment. **A.** Localisation of NCOA4 around parasites during *Leishmania* infection at 4 hpi, 24 hpi, and 48 hpi. **B.** Representative immunofluorescence showing Hoechst (grey), zymosan (green) and NCOA4 (red). **C.** Quantification of number of zymosan beads per cell and the number of zymosan beads per cell having NCOA4 around. The quantification was done by counting 38 cells and in three biological replicates. **D.** Quantification off the percent parasites/cell having NCOA4 around them and comparing within 4h PI for dead and live parasites. The quantification was done by counting 41 cells (for infection with dead parasites) and 54 cells (for infection with live parasites) in three biological replicates.

### Figure S8

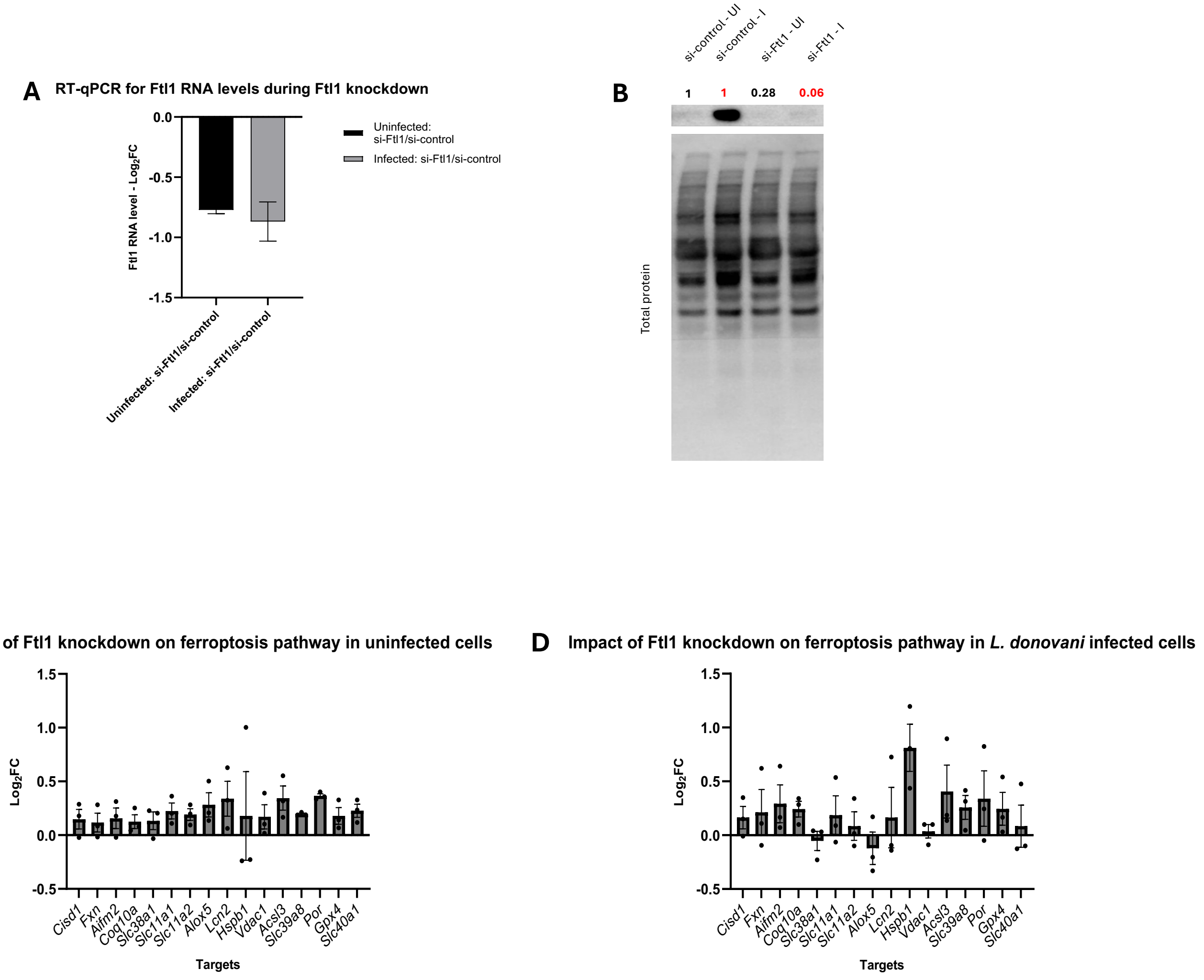


**Figure S8**: Validation of Ftl1 knockdown. **A.** RT-qPCR analysis for change in Ftl1 RNA levels upon Ftl1 KD in uninfected and infected BMDMs. **B.** Western blot for Ftl1 level in si-control uninfected, si-control infected, si-Ftl1 uninfected and si-Ftl1 infected. Comparison of si-Ftl1 uninfected with si-control uninfected (black numbers) and si-Ftl1 infected with si-control infected (red numbers) indicate the fold changes (FC) in the Ftl1 levels upon knockdown in uninfected (0.28 FC) and infected (0.06 FC). Impact of Ftl1 knockdown on the expression levels of the 16 genes in the ferroptosis pathway calculated by RT-qPCR in **C.** Uninfected BMDMs **D.** Infected BMDMs. The error bars represent standard error. Three biological replicates were used for analysis. The details of the genes mentioned is given in Supplementary Table S7. $Log_2FC$ is by comparing si-Ftl1 with si-control.

Supplementary Table S7: Genes representing various aspects of the iron homeostasis pathway

| Target | Name | Role | Inhibit/promote lipid peroxidation | Literature |
|---|---|---|---|---|
| CISD1 | CDGSH Iron Sulfur Domain 1 | Mitochondrial iron flux | Inhibit | (Yuan, Li et al. 2016) |
| FXN | Frataxin | Mitochondrial iron storage | Inhibit | (Du, Zhou et al. 2020) |
| AIFM2 | Ferroptosis suppressor protein 1 | NAD(P)H-dependent oxidoreductase - inhibitor of ferroptosis | Inhibit | (Tang, Chen et al. 2021) |
| COQ10A | Coenzyme Q | Membrane localised antioxidant protecting against lipid peroxidation | Inhibit | (Tang, Chen et al. 2021) |
| slc38a1 | Solute carrier family 38 member 1 | Glutamine transporter | Promote | (Tang, Chen et al. 2021) |
| slc11a1 | NRAMP1 | Iron transporter, on the parasitophorous vacuole membrane | Promote | (Montalbetti, Simonin et al. 2013) |
| slc11a2 | DMT1 | Iron importer | Promote | (Montalbetti, Simonin et al. 2013) |
| alox5 | 5-lipoxygenase | Promotes polyunsaturated fatty acid (PUFA) peroxidation | Promote | (Tang, Chen et al. 2021) |
| LCN2 | Lipocalin-2 | Iron sequestration and trafficking | Inhibit | (Xiao, Yeoh et al. 2017) |
| HSBP1 | Heat shock protein 25 | Inhibit Fenton reaction | Inhibit | (Sun, Ou et al. 2015) |
| VDAC1 | Voltage-dependent anion channel 1 | Iron import into mitochondria | Inhibit | (Zhou, Tang et al. 2023) |
| ACSL3 | Acyl CoA synthetase | Produces monounsaturated fatty acid co-A thus reducing amount of PUFAs | Inhibit | (Yang, Zhu et al. 2022) |
| Slc39a8 | ZIP8 | Iron import | Promote | (Zhang, Wang et al. 2024) |
| POR | P450 oxidoreductase | Promotes PUFA peroxidation | Promote | (Tang, Chen et al. 2021) |
| GPX4 | Glutathione peroxidase 4 | Detoxification of lipid peroxidation | Inhibit | (Tang, Chen et al. 2021) |
| slc40a1 | Solute carrier family 40 member 1/Ferroportin | Iron exporter from cytosol | Inhibit | (Montalbetti, Simonin et al. 2013) |
